# Conformational changes of baseplate regulating tail contraction of *Staphylococcus* phage 812

**DOI:** 10.1101/2024.09.19.613683

**Authors:** Ján Bíňovský, Marta Šiborová, Maryna Zlatohurska, Jiří Nováček, Pavol Bárdy, Roman Baška, Karel Škubník, Tibor Botka, Martin Benešík, Roman Pantůček, Konstantinos Tripsianes, Pavel Plevka

## Abstract

Phages with contractile tails employ elaborate mechanisms to penetrate bacterial cell walls and deliver their genomes into the host cytoplasm. Here, we used cryo-EM to show that the baseplate of phage 812, a member of the *Kayvirus* genus, which infects Gram-positive *Staphylococcus* strains, is formed of a core, wedge modules, and baseplate arms carrying receptor-binding proteins 1 and 2 and tripod complexes. Upon binding to a host cell, the receptor-binding proteins of phage 812 baseplate reorient and undergo conformational changes. The changes to the tripod complexes trigger the release of the central spike and weld proteins, which expose peptidoglycan-degrading domains of the hub proteins. Changes in the positions of baseplate arms are transmitted through wedge modules to tail sheath initiator proteins. The ring of the tail sheath initiator proteins expands and triggers the contraction of the tail sheath, which shortens to 50% and pushes the tail tube 10-30 nm into the bacterial cytoplasm. Homologous molecular mechanisms are probably shared by phages of the *Herelleviridae* family with contractile tails to infect Gram-positive bacteria.

## Introduction

Tailed phages use baseplates at the end of their tails to attach to the host cell surface to initiate infection (Nobrega *et al*, 2018). The cell binding of phages with contractile tails is coupled with a conformational change to the baseplate that triggers tail sheath contraction (Leiman & Shneider, 2012). The tail tube penetrates the cell wall and enables the ejection of the phage genome into the bacterial cytoplasm. Most of the phages with contractile tails and contractile injection systems (CISs) structurally characterized to date attack Gram-negative bacteria (Taylor *et al*, 2016; Li *et al*, 2023; Wang *et al*, 2023b; Yang *et al*, 2023a, 2023b; Yu *et al*, 2024; Sonani *et al*, 2024; Marín-Arraiza *et al*, 2025). Current knowledge regarding the mechanisms employed to breach the cell walls of Gram-positive bacteria is limited.

Phages of the genus *Kayvirus* of the family *Herelleviridae* (Barylski *et al*, 2020) exhibit high lytic activity against the Gram-positive bacterium *Staphylococcus aureus* (Botka *et al*, 2019; Łobocka *et al*, 2012; Vandersteegen *et al*, 2011). This human pathogen can cause skin and epithelial infections, as well as bacteremia that can develop into life-threatening sepsis (Kwiecinski & Horswill, 2020; Grunenwald *et al*, 2018). *S. aureus* is frequently resistant to antibiotics, which has renewed interest in phage therapy (Venturini *et al*, 2022; Petrovic Fabijan *et al*, 2020; Uyttebroek *et al*, 2022). The best studied *Kayvirus*, phage K, shares 98.5% of the genome identity with a polyvalent phage 812. Host range mutants of phage 812 were shown to infect more than 90 % of *S. aureus* isolates (Pantůček *et al*, 1998; Botka *et al*, 2019). Major components of the *S. aureus* cell wall are peptidoglycan and teichoic acids (Schaechter, 2009; van Dalen *et al*, 2020). Phage 812 and related *Kayvirus* phages recognize the backbone and side-chain moieties of the wall teichoic acid (Xia *et al*, 2011; Takeuchi *et al*, 2016; Krusche *et al*, 2025).

It has been shown that a virion of 812 is formed by an icosahedral head with a diameter of 90 nm and a 240-nm-long tail that ends with a lentil-shaped baseplate (Nováček *et al*, 2016). Upon binding to the *S. aureus* cell wall, the baseplate transforms into a double-layered disc-shaped conformation. Furthermore, the structure of phage A511 of the *Herelleviridae* family, which infects Gram-positive *Listeria* and is morphologically similar to 812, has also been determined (Guerrero-Ferreira *et al*, 2019). However, the resolutions of the previous studies of 812 and A511 were limited to 13 and 14 Å, respectively, which prevented the identification of some of the baseplate components and characterization of their conformational changes that enable cell wall binding, tail contraction, and genome delivery.

The naming convention for baseplate components was established in the pioneering studies of phage T4. The core of the baseplate is formed by the central hub (Taylor *et al*, 2016, 2018), which serves as a centerpiece for the baseplate assembly and a platform for the tail tube polymerization. Six wedge complexes encircle the core and provide the foundation for tail sheath assembly (Taylor *et al*, 2016, 2018). Baseplate core proteins of contractile-tailed phages exhibit high structural conservation (Taylor *et al*, 2016, 2018; Veesler & Cambillau, 2011). In addition, the structures of the baseplate core proteins are shared by CISs (Park *et al*, 2018; Ge *et al*, 2020; Weiss *et al*, 2022; Xu *et al*, 2022; Desfosses *et al*, 2019; Jiang *et al*, 2019; Cai *et al*, 2024). The outer baseplate of contractile-tailed phages can be either simple, containing one or two tail fibers to bind to host cellular receptors (Li *et al*, 2023; Wang *et al*, 2023b; Yang *et al*, 2023a, 2023b; Yu *et al*, 2024; Sonani *et al*, 2024), or more intricate, such as that of T4 (Taylor *et al*, 2016).

Here, we used a combination of cryogenic electron microscopy (cryo-EM), X-ray crystallography, nuclear magnetic resonance spectroscopy (NMR), and AlphaFold2 predictions to build the molecular structure of the tail and baseplate in extended and contracted conformations of a *Kayvirus* representative, 812. Based on a comparison of the pre- and post-contraction structures, we propose a sequence of conformational changes that propagate from the baseplate periphery to the core and trigger the tail sheath contraction.

## Results and Discussion

### The structure of baseplate of 812 with extended tail

The structure of the pre-contraction 812 baseplate with imposed sixfold symmetry was previously determined to a resolution of 13 Å using cryo-EM (Nováček *et al*, 2016). Here, we show that the 812 baseplate, similar to that of A511 (Guerrero-Ferreira *et al*, 2019), is only approximately sixfold symmetric. An initial asymmetric reconstruction with distinct three-fold symmetric features enabled the determination of the overall baseplate structure to a resolution of 5.9 Å after imposing the C3 symmetry (Fig. 1, Appendix Figure S1A, B, Appendix Table S1). The baseplate of 812 is lentil-shaped with a diameter of 655 Å and height of 310 Å, and its constituent proteins can be assigned to a core, six wedge modules, and six arms decorated with three types of putative receptor-binding proteins (Fig. 1, Appendix Table S2).

**Fig. 1.**
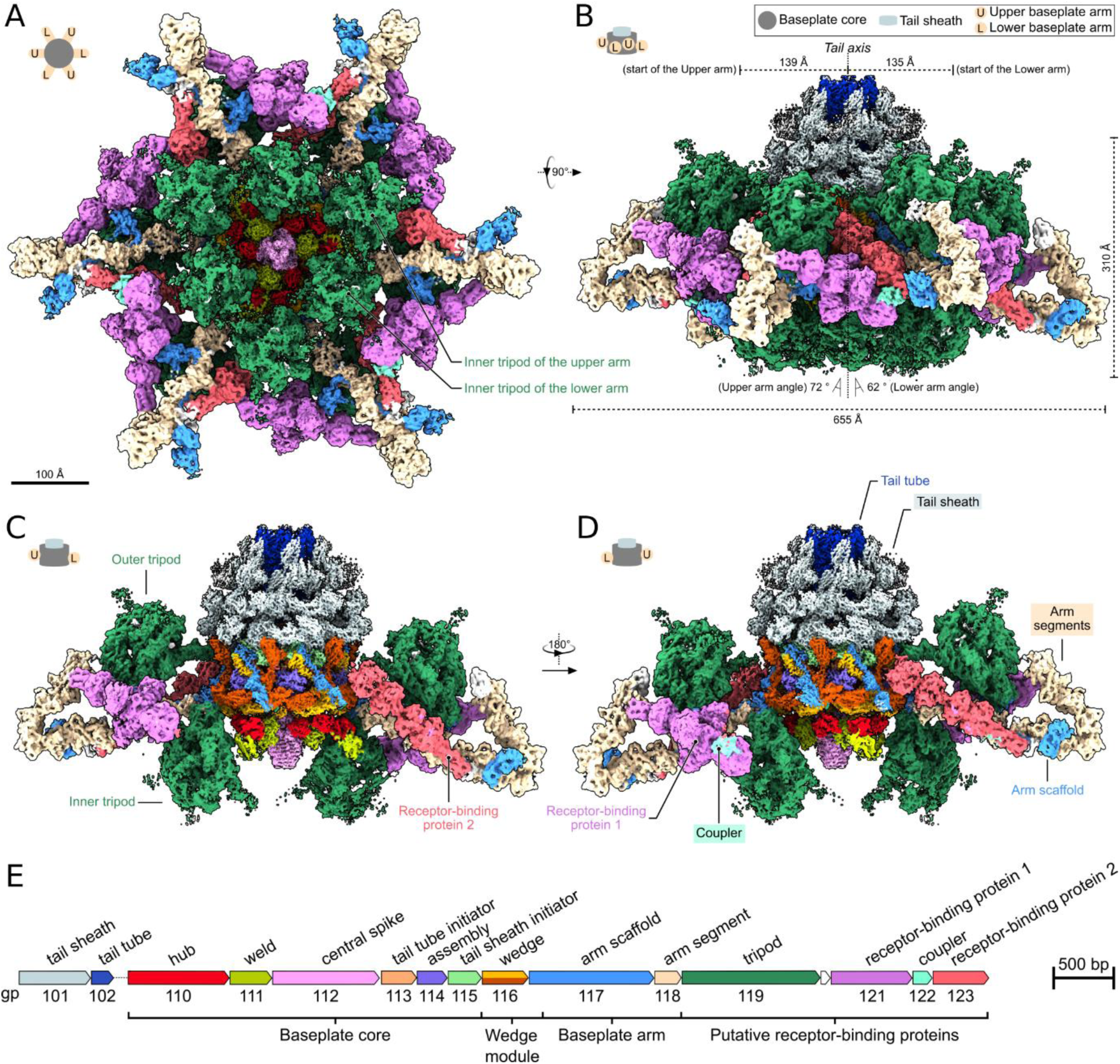
Structure of phage 812 baseplate. **(A-D)** The composite cryo-EM map of the 812 baseplate is viewed along (A) and perpendicular (B-D) to the tail axis. Protein components of the baseplate and tail are colored according to the scheme depicted in panel E. Pictograms indicate the upper and lower baseplate arms. The height and width of the baseplate, tail axis, angles of baseplate arms relative to the tail axis, and distances of baseplate arms from the tail axis are indicated. **(C, D)** Side views of the 812 baseplate, with two front and two rear baseplate arms removed to display the wedge modules and baseplate core. **(E)** Scheme of part of the 812 genome encoding tail and baseplate genes.

Each of the baseplate arms is formed by the arm scaffold protein (GenBank: KJ206563.2, gp117) and the arm segment proteins (gp118) (Fig. 1). The baseplate arm provides attachment sites for proteins with putative receptor binding functions: two trimers of the tripod proteins (gp119), a trimer of receptor-binding protein 1 (RBP1, gp121), and a trimer of receptor-binding protein 2 (RBP2, gp123) (Fig. 1A-D). The trimer of tripod proteins located closer to the tail axis (inner tripod) is oriented with its receptor-binding domains pointing away from the phage head (Fig. 1A-D). In contrast, the receptor-binding domains of tripod proteins attached to the baseplate arm further away from the tail axis (outer tripod) point towards the phage head (Fig. 1B-D). Trimers of RBP1 and RBP2 are attached to the end of the baseplate arm (Fig. 1A-D). The RBP1 reaches from one baseplate arm to the neighboring one, where it interacts with RBP2 (Fig. 1A, B). RBP2 points towards the tail sheath (Fig. 1C, D). In every alternating baseplate arm, the RBP1-RBP2 interaction is mediated by a coupler protein (gp122) (Fig. 1A, D).

The threefold symmetry of the 812 baseplate is most apparent in the distinct conformations of the alternating baseplate arms and the positions of the putative receptor-binding proteins attached to them (Fig. 1A-D). The alternating arrangement of baseplate arms is caused by different angles at which the arms point away from the tail axis (Fig. 1B). The upper baseplate arms radiate from the tail axis at an angle of 72°, while the lower baseplate arms radiate at an angle of 62°. The differences in the baseplate arm angles position the six inner tripods to form a shape of an equilateral triangle when viewed along the tail axis towards the phage head (Fig. 1A). The distinct angles of baseplate arms also influence the interactions of tripod proteins with the tail sheath. The outer tripod attached to the upper baseplate arm interacts with the first tail sheath disc (Appendix Figure S2A). In contrast, the outer tripod bound to the lower baseplate arm does not interact with the tail sheath (Appendix Figure S2B).

Each baseplate arm protrudes radially from a wedge module. A localized reconstruction of the wedge modules and baseplate core reached a resolution of 3.0 Å (Fig. 2, Appendix Figure S1C, S3, S4). Each wedge module is a heterotrimer formed by two copies of wedge protein (gp116) and an N-terminal part of the arm scaffold protein (gp117) (Fig. 2, Appendix Figure S4). Two neighboring wedge modules that connect to the upper and lower baseplate arms assume distinct conformations (Appendix Figure S5A-C).

**Fig. 2.**
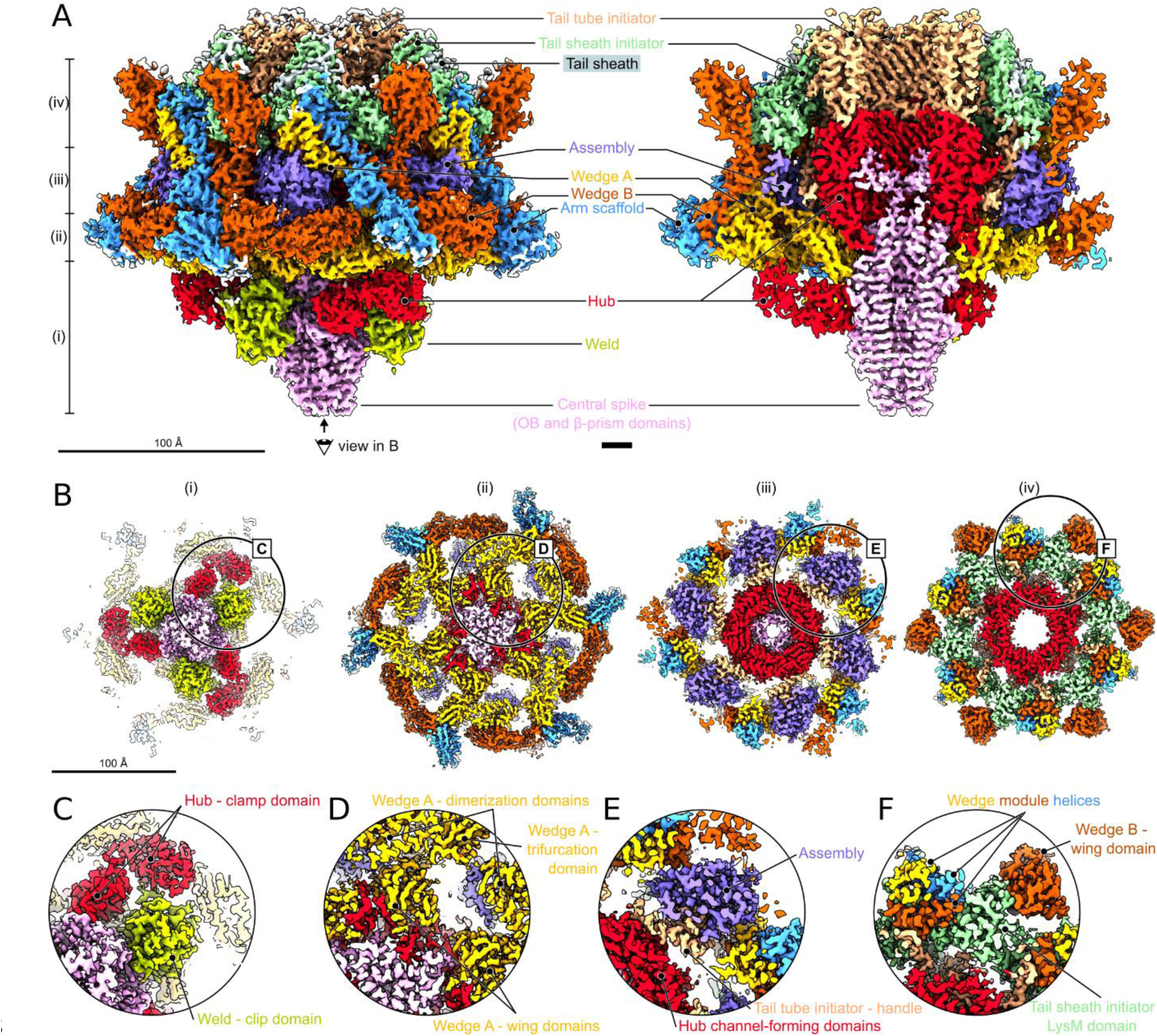
Structure of 812 wedge modules and baseplate core. **(A-F)** The cryo-EM map of wedge modules and baseplate core. Individual proteins are colored according to the scheme depicted in Fig. 1E. Names of selected proteins and protein domains are indicated. **(A)** Side view of wedge modules and baseplate core shown as a whole (left) and with front half removed (right). The positions of sections shown in panel B are indicated on the left. **(B)** Sections through the wedge modules and baseplate core viewed towards the phage head. Positions of close-up views of selected regions displayed in panels C-F are indicated as black circles. **(C-F)** Close-up views of selected regions from sections in panel B.

Six wedge modules encircle the baseplate core composed of tail tube initiator, tail sheath initiator, hub, assembly, central spike, and weld proteins (Fig. 2, Appendix Figure S5). The hexamer of tail tube initiator proteins (gp113) and trimer of hub proteins (gp110) form a channel, which extends the tail tube channel into the baseplate. The trimer of central spike proteins (gp112), bound to the three copies of weld protein (gp111), plugs the baseplate channel at the level of the hub proteins (Fig. 2A, B). The hexamer of tail tube initiator proteins is surrounded by six copies of each of the tail sheath initiator (gp115) and assembly proteins (gp114), which are placed above each other (Fig. 2A, Appendix Figure 5).

It is likely that the overall threefold symmetry of the 812 baseplate is determined during assembly by the asymmetric addition of baseplate proteins to the trimer of hub proteins. The structural differences propagate into the baseplate periphery, resulting in the alternating angles of the baseplate arms relative to the tail axis (Fig. 1B). The same mechanism is probably employed to establish the threefold symmetry of the A511 baseplate (Guerrero-Ferreira *et al*, 2019). In contrast, baseplate core proteins of structurally characterized phages were shown to be arranged with quasi-sixfold symmetry (Taylor *et al*, 2016; Li *et al*, 2023; Wang *et al*, 2023b; Yang *et al*, 2023a, 2023b; Yu *et al*, 2024; Sonani *et al*, 2024).

### Central spike protein blocks the baseplate channel and protrudes from the baseplate

The trimer of central spike proteins of 812 blocks the baseplate channel and protrudes more than 30 nm from the baseplate (Fig. 3A, B). The central spike protein can be divided into oligosaccharide-binding (OB), β-prism, coiled-coil, knob, and petal domains (Appendix Figure S6A, Appendix Table S3). The OB and β-prism domains were resolved in the reconstruction of the pre-contraction 812 baseplate (Fig. 2A). The structure of the coiled-coil domain was predicted using AlphaFold2 (Jumper *et al*, 2021) (Fig. 3B, Appendix Figure S6A), and structures of the knob and petal domains were determined by cryo-EM and single-particle reconstruction of a recombinantly produced central spike protein (Fig. 3B, Appendix Figure S7H, Appendix Table S1). The OB and β-prism domains have conserved structures shared by central spike proteins of other phages and CISs (Ge *et al*, 2020; Jiang *et al*, 2019; Xu *et al*, 2022; Li *et al*, 2023; Yang *et al*, 2023a, 2023b; Yu *et al*, 2024; Sonani *et al*, 2024; Desfosses *et al*, 2019; Weiss *et al*, 2022). The two domains bind to the inside of the channel formed by the hub proteins (Fig. 2A, B, Appendix Figure S4). The coiled-coil, knob, and petal domains protrude from the 812 baseplate and are not resolved in the cryo-EM reconstruction of the baseplate, due to the coiled-coil domain’s flexibility (Fig. 3A). Structural prediction indicates that the coiled-coil domains from the three central spike proteins form a 22-nm-long fiber, reminiscent of the tail needle from phage P22 (Bhardwaj *et al*, 2009) (Fig. 3B, Appendix Figure S6A). The knob domain has a tumor necrosis factor-like fold (Eck & Sprang, 1989), similar to that of the head domains of receptor-binding proteins from lactococcal phages Tuc2009 and 1358 (Legrand *et al*, 2016; Farenc *et al*, 2014; McCabe *et al*, 2015) and a lectin from *Burkholderia cenocepacia* (Šulák *et al*, 2010) which indicates that the knob domain may bind to saccharides from the bacterial cell wall. The petal domain of the central spike protein has an α/β-barrel fold structurally similar to phosphodiesterases, such as the GlpQ protein from *B. subtilis* that cleaves teichoic acid during phosphate starvation (Myers *et al*, 2016; Shi *et al*, 2008). Additionally, the petal domain includes two conserved histidines commonly found in active sites of phosphodiesterases, suggesting the domain’s putative cleavage activity (Fig. 3C).

**Fig. 3.**
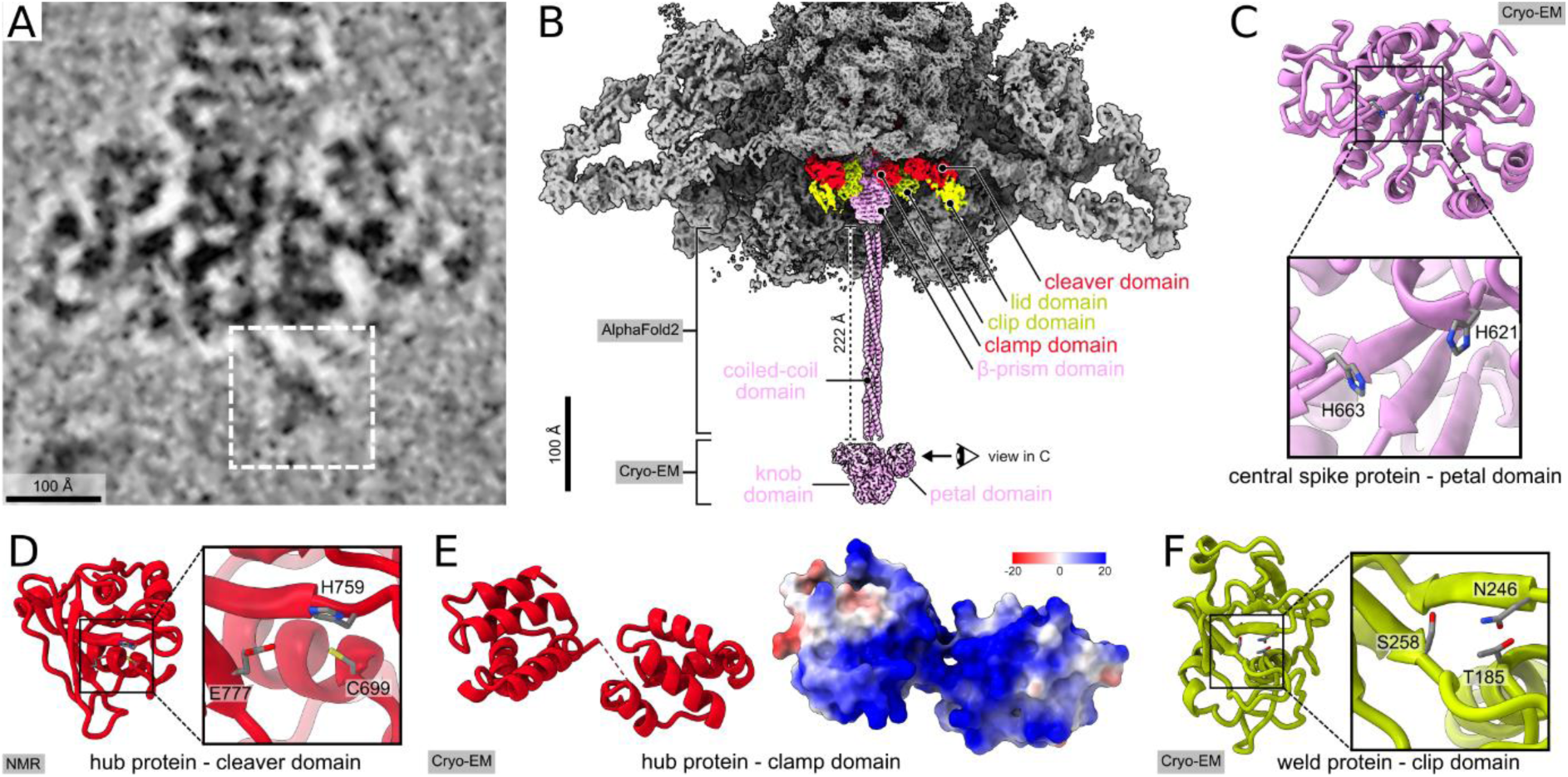
Central spike, hub, and weld proteins. **(A)** Cryo-electron micrograph of the 812 baseplate. The white dashed rectangle indicates the putative position of the coiled-coil, knob, and petal domains of the trimer of central spike proteins. **(B)** Composite cryo-EM map of 812 baseplate and cryo-EM map of the knob and petal domains of central spike protein trimer. The baseplate is shown in gray except for the central spike (pink), hub (red), and weld (green) proteins. The front part of the baseplate has been removed. The coiled-coil domains of the central spike proteins predicted using AlphaFold2 are shown in cartoon representation. **(C, D)** Cartoon representations of the petal domain of the central spike protein (C) and the cleaver domain of the hub protein (D). Side chains of residues forming putative hydrolytic sites are shown in stick representation. Insets show details of the putative hydrolytic sites. **(E)** Cartoon (left) and surface (right) representations of the clamp domain of the hub protein. The molecular surface is colored according to Coulombic electrostatic potential in the range from -20 to 20 kcal/(mol·*e^-^*) at 298 K. **(F)** Cartoon representation of the clip domain of the weld protein. Side chains of residues forming a putative receptor-binding site are shown as sticks. The structures shown in panels C-F were determined using cryo-EM or NMR, as indicated.

Phages with contractile tails and CISs attacking Gram-negative bacteria pierce the outer bacterial membranes using puncturing tips formed of proteins with β-prism or β-helix domains (Browning *et al*, 2012; Kanamaru *et al*, 2002; Shneider *et al*, 2013). In contrast, to infect Gram-positive *S. aureus*, phage 812 has to penetrate a 50-70-nm-thick layer of teichoic acids and peptidoglycan. Therefore, we propose that the knob and petal domains of the central spike of 812 bind to and degrade the host teichoic acids to expose the peptidoglycan layer for degradation by the cleaver domain of the hub protein (see below).

### Hub proteins facilitate penetration of the tail tube through the cell wall and cell membrane

The three hub proteins form a ring, extending the tail channel into the 812 baseplate (Fig. 2A-C). The hub protein can be divided into channel-forming domains I to IV and clamp and cleaver domains (Appendix Figure S6B, Appendix Table S3). The channel-forming domains I and III have beta-barrel folds and are positioned next to each other so that the head-proximal interface formed by the trimer of hub proteins has pseudo-sixfold symmetry for binding the hexamer of tube initiator proteins (Appendix Figure S4). In contrast, the channel-forming domains II and IV of the hub protein, positioned below domains I and III, respectively, are structurally distinct and form interfaces for binding the trimer of central spike proteins to the inside of the hub channel (Appendix Figure S4). The channel-forming domains I-IV are structurally similar to those of the gp27 of phage T4, for which the domain naming convention was established (Kanamaru *et al*, 2002). Homologs of the channel-forming domains can also be identified among phages with long non-contractile tails (Tal proteins) (Veesler & Cambillau, 2011; Kizziah *et al*, 2020) and CISs (Vgrg-like proteins) (Taylor *et al*, 2018; Büttner *et al*, 2016; Leiman *et al*, 2009) (Appendix Table S3).

The cleaver domain of the hub protein, the structure of which was determined using NMR (Fig. 3D, Appendix Figure S6B, Appendix Table S4), has the characteristic fold of the cysteine- and histidine-dependent amido-hydrolase/peptidase (CHAP) (Bateman & Rawlings, 2003), and contains the conserved catalytic triad Cys699-His759-Glu777 (Sanz-Gaitero *et al*, 2014) (Fig. 3D). Accordingly, zymogram analysis demonstrated that a recombinant cleaver domain degrades *S. aureus* cell walls (hence the name cleaver) (Appendix Figure S8), which agrees with previous functional studies of hub protein homologs from the related phage K (Becker *et al*, 2008; Paul *et al*, 2011). The clamp domain of the hub protein is formed by ten short α-helices that connect the central spike protein to weld proteins (Fig. 2B, C, and 3E). The clamp domain is structurally similar to pore-forming saposin-like proteins, which employ positively charged residues to interact with the negatively charged surfaces of lipid membranes (Willis *et al*, 2011; Lee *et al*, 2006) (Appendix Table S3). Since the clamp domain of the 812 hub protein contains a long positively charged patch of residues, as the saposin-like proteins do (Fig. 3E), we speculate that it may bind the *S. aureus* membrane to facilitate its penetration by a tail tube. In addition to gp27 of T4 featuring a lysozyme domain, more hub proteins equipped with enzymatic domains have been reported recently (Yu *et al*, 2024; Cai *et al*, 2024). Therefore, we propose that the hub proteins of 812 and related phages enable genome delivery by degrading peptidoglycan to facilitate penetration of the tail tube through the cell wall.

### Tail tube initiator, weld, assembly, and tail sheath initiator proteins complete the baseplate core

Three weld proteins bind to the central spike, hub, and three inner tripod proteins, connecting them together (Fig. 2A-C, Appendix Figure S4 and S9). Six assembly and six tail sheath initiator proteins surround the hub and tail tube initiator proteins to complete the structure of the baseplate core (Fig. 2A, B, E, F, Appendix Figure S4). The tube initiator and sheath initiator proteins are organized with sixfold symmetry. In contrast, the central spike, weld, hub, and assembly proteins are organized with threefold symmetry that determines the overall organization of the baseplate when the 812 tail is in extended conformation.

The weld protein of 812 can be divided into the N-terminal lid and C-terminal clip domains, which are connected by a 14-residue-long linker (Appendix Figure S6D). The lid domain interacts with an inner tripod and serves as a lid covering the peptidase site of the cleaver domain of the hub protein (Appendix Figure S9). Therefore, releasing the weld protein from the baseplate upon host cell binding exposes the active site of the cleaver domain of the hub protein to hydrolyze the cell wall peptidoglycan. The clip domain of the weld protein, located closer to the tail axis, binds the weld protein to the central spike protein (Fig. 2C). The clip domain has the CHAP protein fold similar to that of the cleaver domain of the hub protein (Gillespie *et al*, 2015; Wu *et al*, 2019) (Fig. 3F). However, the clip domain lacks the conserved catalytic residues Cys-His-Glu required for the peptidoglycan cleavage (Sanz-Gaitero *et al*, 2014) (Fig. 3F). Thus, the clip domain is likely enzymatically inactive, but may have retained peptidoglycan-binding ability, similar to the RipD protein from *M. tuberculosis* (Böth *et al*, 2014). There is no homolog of the weld protein in the baseplate of the related phage A511 (Guerrero-Ferreira *et al*, 2019; Habann *et al*, 2014). The weld protein may be unnecessary for A511 because of the differences in the cell wall composition between *Staphylococcus* and *Listeria* (Schaechter, 2009).

The assembly protein of 812 is formed by a compact bundle of nine α-helices. The assembly protein interacts with the wedge protein and the C-terminal handle of the tail tube initiator protein (Fig. 2E, Appendix Figure S4). There are proteins with a compact globular structure and in the equivalent position in the baseplates of pyocin R2 and phages Pam3, E217, A-1, and Milano (Appendix Table S3), indicating the structural role of these proteins in baseplate cores (Ge *et al*, 2020; Li *et al*, 2023; Yang *et al*, 2023a; Yu *et al*, 2024; Sonani *et al*, 2024).

The hexamer of tail tube initiator proteins binds to the trimer of hub proteins and, on the other side, to a hexamer of tail tube proteins (Fig. 2A). The tail tube initiator protein can be divided into the N-terminal β-barrel domain and the C-terminal α-helical handle, which stretches along the hub protein and reaches the assembly protein in the baseplate core (Appendix Figure S4, S6C). The tail tube initiator proteins from pyocin R2 and phages E217 and Pam3 contain extensions similar to the C-terminal handle of the 812 tail tube initiator protein, suggesting an evolutionary link between phage and pyocin baseplates (Leiman & Shneider, 2012; Veesler & Cambillau, 2011).

The tail sheath initiator protein of 812 is composed of the LysM and handshake domains (Appendix Figure S6E). The handshake domain, containing two β-strands, interacts with the N- and C-termini of tail sheath proteins from the first tail sheath disc and the C-terminus of tail sheath protein from the second tail sheath disc to form a five-stranded β-sheet (Appendix Figure S10C). Therefore, the interactions of the tail sheath and tail sheath initiator proteins can initiate assembly of the 812 tail sheath. Proteins containing domains with structures similar to the handshake domain are integral to the baseplates of most CISs and phages with contractile tails (Taylor *et al*, 2018; Büttner *et al*, 2016; Maxwell *et al*, 2013) (Appendix Table S3).

### Wedge modules attach baseplate arms to the baseplate core

Wedge modules are structural components of phage baseplates, which enable the attachment of peripheral baseplate proteins to the baseplate core (Taylor *et al*, 2016; Watts & Coombs, 1990). Here, we show that the wedge module of 812 is formed by two wedge proteins with different conformations, A and B, and the N-terminal 176 residues of the arm scaffold protein (Fig. 2A, Appendix Figure S4 and %B, C). Wedge protein can be divided into N-terminal α-helices, wing, trifurcation, and dimerization domains (Appendix Figure S6F, G), and the part of the arm scaffold protein belonging to the wedge module can be divided into N-terminal α-helices and a trifurcation domain (Fig. 4A, B, D). Individual parts of the 812 wedge module are structurally conserved and can also be found in the baseplate of phage T4 (Taylor *et al*, 2016). The N-terminal helices and trifurcation domains mediate contacts between the proteins forming the wedge module (Fig. 2F, Appendix Figure S5A-C). The wing domain of wedge protein A binds to the hub and central spike proteins (Fig. 2D). In contrast, the wing domain of wedge protein B binds to the tail sheath initiator protein (Fig. 2F). The six wedge modules form an iris around the baseplate core (Appendix Figure S5A), as described in phage T4 (Taylor *et al*, 2016). Dimerization domains of wedge proteins mediate interactions between neighboring wedge modules (Appendix Figure S58A). Even though the neighboring wedge modules differ in their conformations due to the overall threefold symmetry of the baseplate, the contacts between the dimerization domains are similar (Appendix Figure S5A-C, F).

**Fig. 4.**
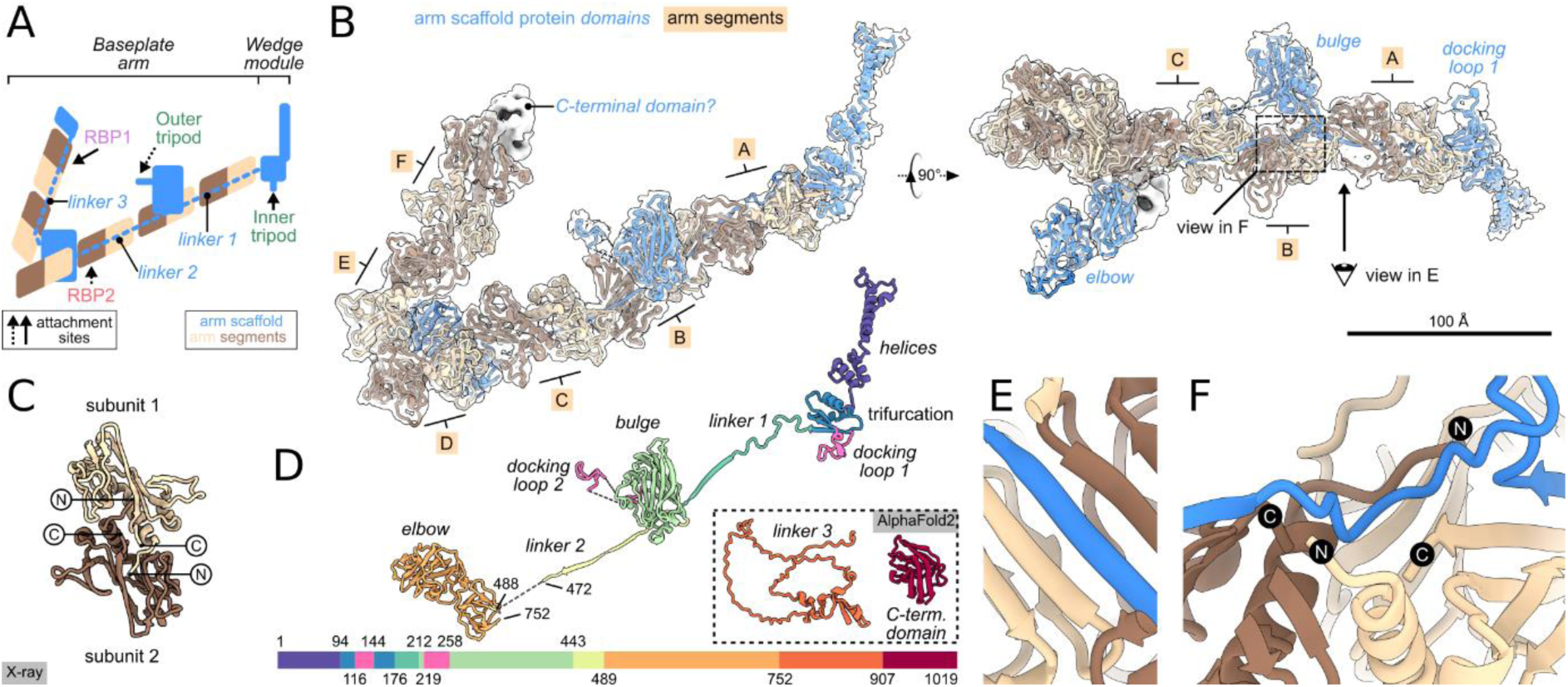
Structure of baseplate arm. **(A)** Schematic representation of the baseplate arm and part of the wedge module. Attachment sites of tripods, RBP1, and RBP2 are indicated. **(B)** Composite cryo-EM map of the lower baseplate arm with structures of arm scaffold and arm segment proteins shown in cartoon. The distal end of the baseplate arm, probably corresponding to the C-terminal domain of the arm scaffold protein, is shown as a white surface. Parts of the baseplate arm with fitted structures are shown as white semi-transparent surfaces Arm segment dimers A to F are indicated. **(C)** The structure of the dimer of arm segment proteins, determined using X-ray crystallography, is shown in a cartoon **(D)** Cartoon representation of the domains of the arm scaffold protein. Domain names and selected chain termini are indicated. Domain structures predicted using AlphaFold2 and not fitted into the cryo-EM maps are shown in dotted rectangle. The linear diagram (bottom) representing arm scaffold protein domains is colored according to the structures. **(E, F)** Cartoon representation of the interaction of the C-terminal β-strands of arm segment proteins with the arm scaffold protein. Views are indicated in the panel B.

### Baseplate arms provide attachment sites for putative receptor-binding proteins

To address the flexibility of the lower and upper baseplate arms from a phage with the extended tail, a series of sub-particle reconstructions were performed, resulting in resolutions ranging from 3.8 to 6.3 Å (Appendix Figure S1E-O and S3, Appendix Table 1). Baseplate arms of 812 are formed by docking loop 1, linker 1, bulge domain including docking loop 2, linker 2, elbow domain, linker 3, and C-terminal domain of the arm scaffold protein that are reinforced by six dimers of the arm segment proteins named A-F (Fig. 4A, B, D, Appendix Figure S11A). We employed X-ray crystallography to determine the structure of the arm segment protein (Fig. 4C, Appendix Table S5) and AlphaFold2 (Jumper *et al*, 2021) to predict the structures of the arm scaffold protein domains (Appendix Figure S11A). The structures were fitted and refined into the sub-particle reconstruction cryo-EM maps of the upper and lower arms. The arm segment protein is composed of an N-terminal α-helix followed by a three-stranded β-sheet and a bundle of loops and short β-strands (Fig. 4C). The overall structure of the arm segment protein is similar to those of gp8 from phage T4 and gp105 from phage A511 (Guerrero-Ferreira *et al*, 2019; Leiman *et al*, 2003). The interactions between the C-terminal β-strands of arm segment proteins from neighboring dimers are mediated by linkers of the arm scaffold protein (Fig. 4E). The linkers of the arm scaffold protein fill the gaps between the dimers of arm segment proteins, creating continuous, seven-stranded β-sheets, as has been speculated previously (Guerrero-Ferreira *et al*, 2019). The cryo-EM map contains resolved density only for the linker 1 and a part of the linker 2 (Fig. 4D). The density likely belonging to the remainder of linker 2, linker 3, and the C-terminal domain could be observed in a difference map after the subtraction of the densities belonging to the arm segment proteins (Fig. 11B-E).

The arm scaffold protein provides attachment sites for two trimers of tripod proteins, one trimer of RBP1, and one trimer of RBP2. The 28 residues-long docking loop 1 of the arm scaffold protein forms the binding site for the inner tripod complex, while the 40-residue-long docking loop 2 enables attachment of the outer tripod complex (Fig. 4A, B, D). The docking loops have triangular structures that fit into crevices formed by the trimers of tripod proteins (Appendix Figure S12A-D). Despite the different length, the docking loops have hydrophobic residues arranged in equivalent positions in the inner and outer tripod crevices (Appendix Figure S12E)

Predicted structure of the elbow domain of the arm scaffold protein with a double β-sandwich fold could be placed at the periphery of baseplate arms, where it forms a protuberance pointing towards the neighboring baseplate arm (Fig. 4B). Close to the elbow domain, a low-resolution density emerging from below the arm segment C likely belongs to a part of the arm scaffold protein that attaches RBP2 (Appendix Figure S11D). Arm segment dimers D, E, and F create an arch at the baseplate periphery, with the linker 3 likely running along the segments (Fig. 4A, Appendix Figure 11B, C). RBP1 is attached to the baseplate arm where a low-resolution density emerges from the arm segment F (Appendix Figure S11E). At the end of the baseplate arm, a density was identified that could accommodate the C-terminal domain of the arm scaffold protein with predicted β-sandwich fold, although it could not be positioned in the reconstruction with confidence (Appendix Figure S11E).

### Baseplate of 812 contains twelve tripod protein complexes

The most prominent components of the 812 baseplate are the twelve trimers of tripod proteins (Fig. 1A-D). We predicted structures of the tripod proteins in 812 with an extended tail using AlphaFold2 (Jumper *et al*, 2021) and refined their structure in the corresponding parts of the reconstructions of the baseplate arms (Fig. 5A-E, Appendix Figure S13). The tripod protein can be divided into spine, flap, fiber, base, fin, leg, anchor, and hook domains (Fig. 5E). The flap domain contains an insertion loop, three of which play a role in the attachment of the tripod complex to the arm scaffold protein (Fig. 5C-E). The base and fin domains of tripod proteins have ten-stranded β-sandwich folds of carbohydrate-binding module families 16 and 61 (Cid *et al*, 2010; Bae *et al*, 2008) (Fig. 5E, Appendix Table S3), indicating that they may bind glycans from the *S. aureus* cell wall. The leg domain includes fifteen β-strands, eight forming a β-sandwich that is structurally similar to lectin-like protein Cwp84 from *C. difficile* (Bradshaw *et al*, 2014) (Fig. 5E, Appendix Table S3), suggesting another possible receptor-binding site. The anchor domain with a β-sandwich fold is followed by the hook domain with sequence similarity to LysM proteins, indicating it may bind peptidoglycan (Buist *et al*, 2008) (Fig. 5E, Appendix Figure S14, Appendix Table S3).

**Fig. 5.**
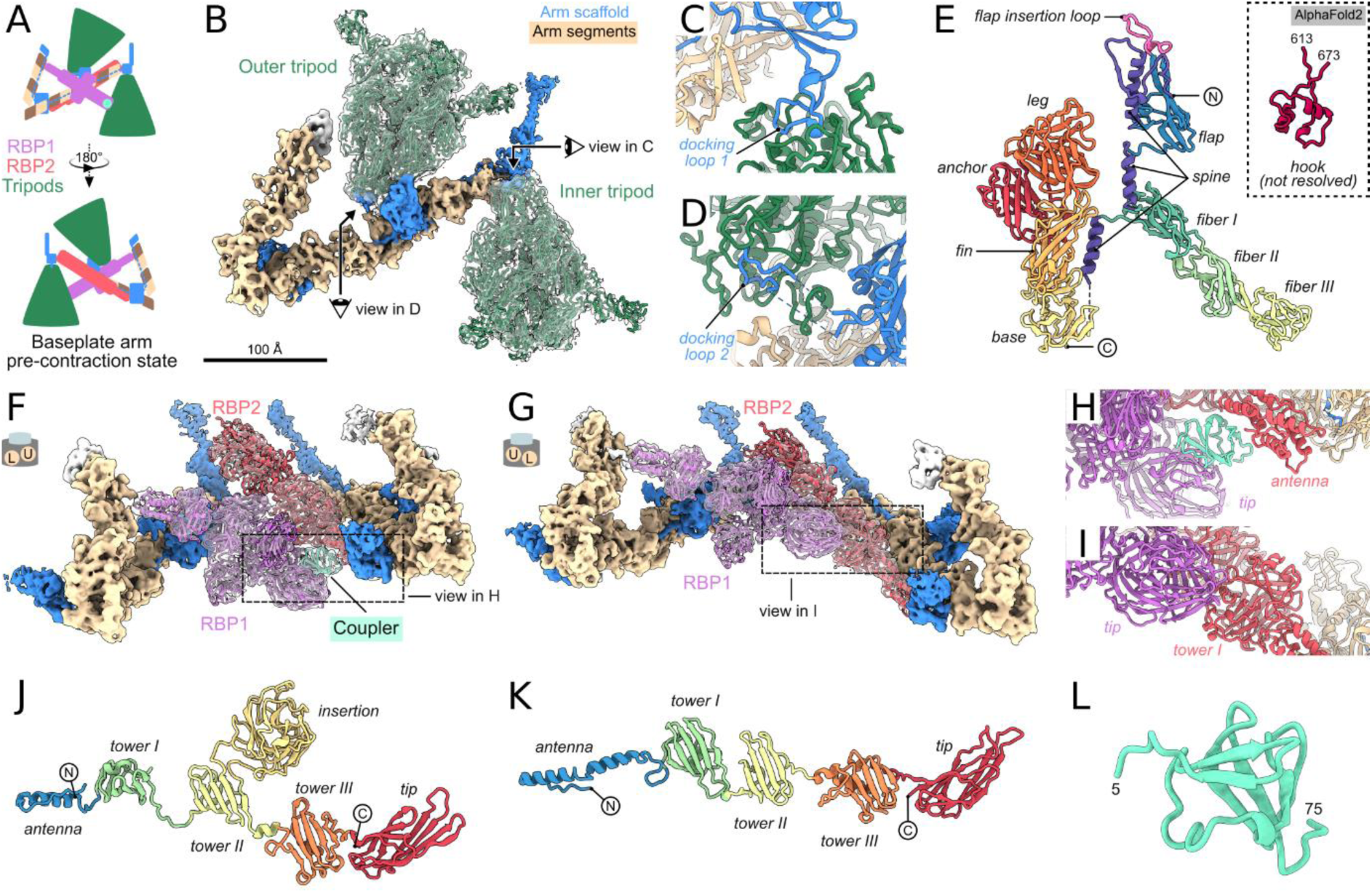
Tripod and receptor-binding proteins of 812 baseplate. **(A)** Schematic representation of the baseplate arm and part of the wedge module of 812 with the extended tail. **(B)** Composite cryo-EM map of the lower baseplate arm. Part of the map corresponding to the inner and outer tripod protein trimers is shown as a white semi-transparent surface. Part of the map corresponding to the arm scaffold and arm segment proteins is shown as an opaque surface colored according to the protein type. Structures of the tripod proteins are shown in cartoon representation. Views of panels C and D are indicated. **(C, D)** Detailed views of attachment sites of inner (C) and outer (D) tripod protein trimers to the docking loops 1 and 2, respectively, of the arm scaffold protein. **(E)** The structure of the tripod protein is shown in cartoon representation and colored by domains. Domain names and chain termini are indicated. **(F, G)** Composite cryo-EM maps of the two neighboring baseplate arms. Parts of maps corresponding to RBP1, RBP2 (F, G) and coupler (F) proteins are shown as a white semi-transparent surface. Parts of maps corresponding to the arm scaffold and arm segment proteins is shown as an opaque surface colored according to the protein type. Structures of RBP1, RBP2 and coupler proteins are shown in cartoon representation. Views of panels H and I are indicated. **(H, I)** Detailed views of protein interactions indicated in panels F (H) and G (I). **(J, K)** Structures of RBP1 (J) and RBP2 (K) monomersare shown in cartoon representation and colored by domains. **(L)** The structure of the coupler protein is shown in cartoon.

### Receptor-binding and coupler proteins

The baseplate of 812 contains six trimers of each of RBP1 and RBP2 with putative receptor-binding activities (Fig. 1A-D). We employed AlphaFold2 (Jumper *et al*, 2021) to predict the structures of RBP1 and RBP2 and refine them in the corresponding parts of the cryo-EM map of the pre-contractionbaseplate (Fig. 5F-K). RBP1 consists of an antenna domain, three tower domains, and a tip domain, all positioned along the symmetry axis of the trimer, and a wing domain protruding laterally from the second tower domain (Fig. 5J). The tower domains are structurally similar to each other and contain characteristic six-stranded β-sheets. The antenna domain mediates the binding of RBP1 to the end of the baseplate arm near the arm segment F and the tip domain of RBP1 points towards the neighboring baseplate arm (Fig. 5F, G). RBP2 has the same domain composition as RBP1, except for the wing domain, which is missing (Fig. 5K). In the baseplate, the antenna domain of RBP2 binds near arm segment C, and the tip domain is in contact with a wedge module (Fig. F, G, Appendix Figure S2). Baseplate of 812 before tail contraction contains three copies of coupler proteins. One coupler protein connects the tip domain of RBP1 from a lower baseplate arm to the antenna domain of RBP2 from the neighboring upper baseplate arm (Fig. 5F). In contrast, RBP1 from the upper baseplate arm interacts directly with RBP2 from the lower baseplate arm (Fig. 5G). Resolved structure of the coupler protein (residues 5-75 out of 124) contains two β-sheets and the closest structural homolog is the tail fiber assembly protein from phage Mu, where it serves as a chaperone for correct folding of the tail fiber (North *et al*, 2019).

### Re-arrangement of RBPs and tripod complexes during tail contraction

Tailed phages use receptor-binding proteins from their baseplates to attach to host cells. For phages with contractile tails, the receptor binding is accompanied by reorganization of the baseplates, which may include conformational changes to individual proteins (Nobrega *et al*, 2018; Taylor *et al*, 2016; Nováček *et al*, 2016). It has been proposed that the reorganization starts at the baseplate periphery, propagates to the inner baseplate, and triggers the contraction of the tail sheath (Leiman & Shneider, 2012; Taylor *et al*, 2016). The overall asymmetric reconstruction of the baseplate of 812 with the contracted tail reached the resolution of 6.6 Å (Appendix Figure S7A, Appendix Table S1). The post-contraction baseplate has the shape of a six-pointed star with a diameter of 713 Å and a thickness of 344 Å (Fig. 6A-C). The conformational change transforms the baseplate to sixfold symmetry, with all six baseplate arms protruding at 83° and positioned at the same distance from the tail axis (Fig. 6A-C, Appendix Figure S7B). Focused reconstructions including tail sheath initiator, wedge module, inner tripod and tail sheath proteins show that the central part of the baseplate is stable, reaching local resolutions up to 3.1 Å (Appendix Figure S7C, D, and S15). The sub-particle reconstruction of the post-contraction baseplate arm contains resolved density of arm segment protein dimers A-C as well as of the docking loop1, linker 1, bulge domain with the docking loop 2, and part of the linker 2 of the arm scaffold protein, inner and outer tripods, and RBP2 (Fig. 6D, Appendix Figure S7E, F, and S15). RBP1, arm segments D-F, and the remainder of the arm scaffold protein are not resolved in the reconstruction of the post-contraction baseplate, likely due to the flexibility of the structure (Fig. 6A-D). Comparing the structures of the baseplates of 812 particles with extended and contracted tails enabled us to describe conformational changes controlling the baseplate rearrangement and tail sheath contraction. The reorganization of the 812 baseplate is probably initiated by the interaction of the RBP1 with teichoic acid as its putative receptor. In the post-contraction baseplate, RBP2 trimers are oriented parallel with the tail axis, and their tip domains face away from the phage head (Fig. 6A-D). The movement of RBP1 and RBP2 provides space for the outer tripod complexes to rotate ∼170° to point their receptor-binding domains away from the phage head (Fig. 6A-D). The inner tripod complexes assume positions with their axes parallel to the tail axis (Fig. 6A-D).

**Fig. 6.**
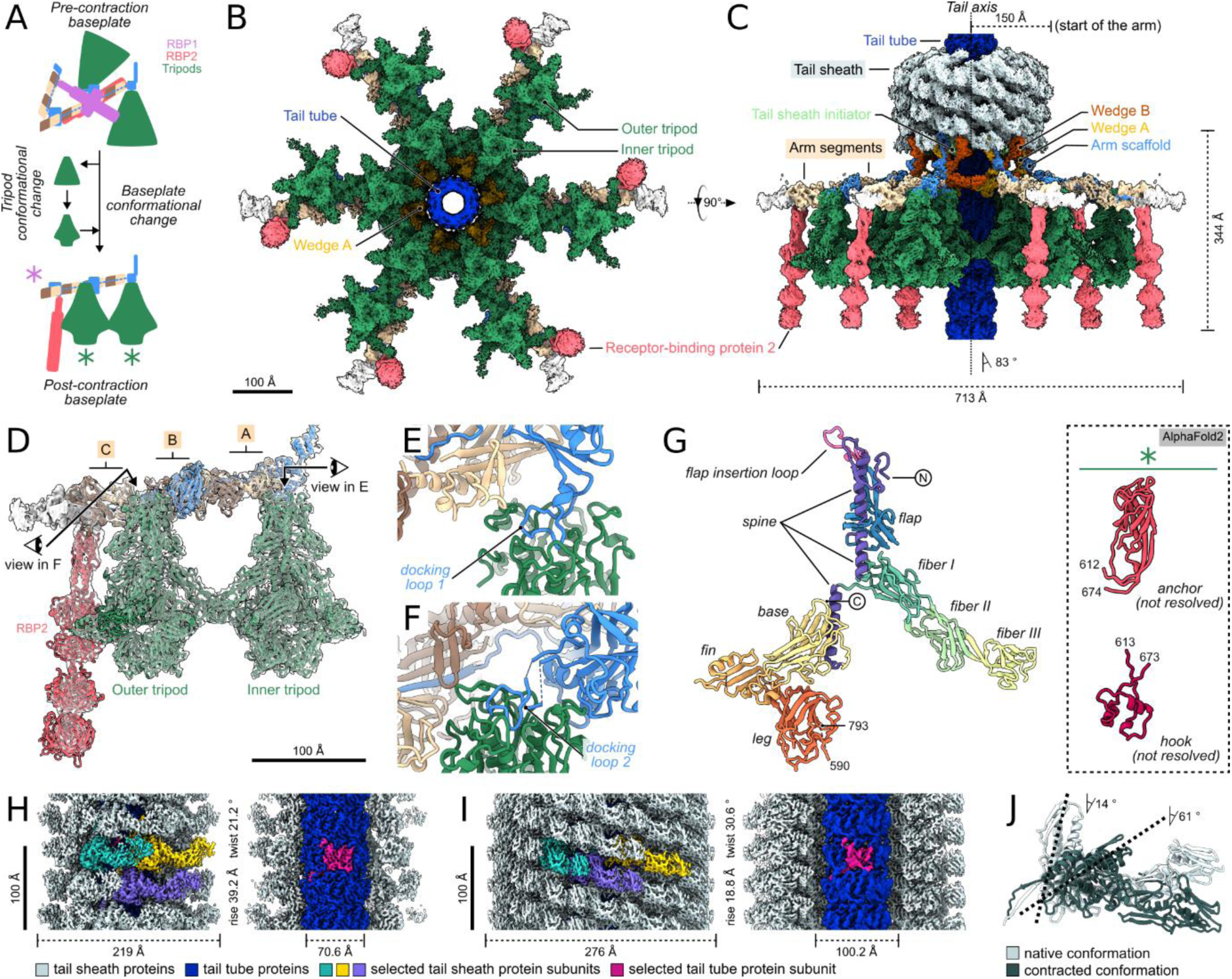
Structures of 812 baseplate, tripods, and tail sheath after the tail sheath contraction. **(A)** Schematic representation of conformational changes of the baseplate arm and tripod complexes. The pink asterisk indicates the putative attachment site of RBP1, which was not resolved in the reconstruction of the contracted baseplate. Green asterisks indicate the putative position of the anchor and hook domains of tripod proteins, which were not resolved in the reconstruction. **(B, C)** Composite cryo-EM maps of the contracted baseplate are viewed along (B) and perpendicular to (C) the tail axis. Proteins are colored according to Fig. 1D, and names of selected components are indicated. In panel C, the height and width of the baseplate, position of the tail axis, angle of baseplate arms relative to the tail axis, and distance of baseplate arms from the tail axis are indicated. **(D)** Composite cryo-EM map of the baseplate arm from contracted baseplate. The distal end of the baseplate arm that could not be interpreted is shown as a white surface. Part of the baseplate arm with fitted structures is shown as white semi-transparent surface. Structures are shown in cartoon representation. Views of panels E and F are indicated. **(E, F)** Detailed views of attachment sites of inner (E) and outer (F) tripod protein trimers to the docking loops 1 and 2, respectively, of the arm scaffold protein. **(G)** Cartoon representation of the tripod protein is colored by domains. The structures of the anchor and hook domains of the tripod protein, which were not resolved in the cryo-EM reconstruction of the contracted baseplate, were predicted using AlphaFold2, and their putative positions are indicated in panel A by green asterisks. Domain names and chain termini are indicated. **(H, I)** Cryo-EM map of the extended tail (H) and composite cryo-EM map of the contracted tail (I). For each panel, complete maps are shown on the left, and maps with the front half of the tail sheath removed shown on the right. Protein components are colored according to the legend below the panel. **(J)** Comparison of the structures of tail sheath proteins in extended and contracted conformations. The structures are shown in cartoon representation, and their domains II were superimposed. Angles of the long axes of domain I relative to the tail axis are indicated for each conformation.

The interactions between the tripod complexes and the docking loops 1 and 2 of the arm scaffold protein are preserved after the baseplate re-organization (Fig 6E, F). As described above, the docking loop 2 of the arm scaffold protein, which binds the outer tripod complex, is twelve residues longer than the docking loop 1 that binds the inner tripod complex (Appendix Figure 12E). The extra residues of the docking loop 2 enable the ∼170° rotation of the outer tripod complexes during the baseplate transformation.

The tripod complexes not only rotate, but also undergo complex conformational changes (Fig. 5E and 6G): (i) The second and third spine helices move towards the first spine helix by 10 and 16 Å, respectively. (ii) The fiber domain moves 13 Å towards the flap domain and rotates 20° counterclockwise around the tripod threefold symmetry axis when looking from the bottom of the tripod. (iii) The base domain shifts 24 Å along the third spine helix while rotating 174° around an axis perpendicular to the tripod symmetry axis. (iv) The fin domain moves 56 Å to protrude laterally from the tripod. (v) The three leg domains detach from the flap domains and move 88 Å to interact with each other under the base domains. (vi) The anchor and hook domains were not resolved in the cryo-EM reconstruction of the baseplate from a phage with a contracted tail (Fig. 6G); however, phage particles attached to the cell wall contain density at the bottom of the tripod complexes, suggesting that the anchor and hook domains stretch along the tripod threefold axis toward the cell wall (Appendix Figure S16). Similar density under the tripod complexes was observed in micrographs of contracted phage A511 attached to *Listeria* cells, and in the baseplate reconstructions of the spontaneously contracted 812 (Nováček *et al*, 2016) and A511 (Guerrero-Ferreira *et al*, 2019). Because of their positioning after the tripod conformational change and their sequence similarity to peptidoglycan-binding proteins of the LysM family (Appendix Figure S14), we speculate that the hook domains of 812 and related phages bind the host cell wall.

The conformational change of tripod proteins is required to form tripod-tripod interactions in phages with contracted tails. In particular, the rotation and translation of the base domain are essential to expose an interface to which a fiber domain of the neighboring tripod complex binds through an interface with a buried surface area of 1,900 Å^2^ (Appendix Figure S17). Each inner tripod complex connects two inner tripods from neighboring baseplate arms and an outer tripod complex attached to the same baseplate arm. Additionally, one fiber domain of each outer tripod complex interacts with tower domains 1 and 2 of RBP2 (Fig. 6B). The contacts mediated by the fiber domains probably stabilize the baseplate structure in phages with contracted tails.

Overall, the recognition and adhesion of 812 to the host cell requires re-orientation of the RBP1, RBP2, and conformational changes of tripod complexes, affecting the positions of the six arms of the post-contraction baseplate. Less extensive conformational changes were described for the receptor-binding proteins of other phages (Li *et al*, 2023; Šiborová *et al*, 2022; Sciara *et al*, 2010).

### Conformational changes to the baseplate core prompt tail sheath contraction

The structure of wedge modules and baseplate core is affected by the rearrangement of receptor-binding proteins and baseplate arms, which in turn trigger the contraction of the tail sheath. The polypeptide chains of arm scaffold proteins link baseplate arms of 812 to the wedge modules (Fig. 4A, B). Therefore, the changes in the position and orientation of baseplate arms cause the iris-like structure formed by the wedge modules to expand in diameter from 104 to 122 Å (Appendix Figure S5A, D). The expansion of the iris is enabled by the movement of the dimerization and trifurcation domains relative to each other, which brings the six wedge modules to an identical conformation (Appendix Figure S5D, E). In contrast, the interactions between the dimerization domains of wedge proteins remain the same as in the baseplate of the extended tail (Appendix Figure S5F). A similar expansion was described for gp6 and gp7 during the transition of the T4 baseplate from its extended to contracted state (Taylor *et al*, 2016). In phage 812, the expansion of the diameter of the tail sheath initiator hexamer, which probably occurs together with that of the wedge iris, is enabled by the partial unfolding of the linker helices between LysM and handshake domains (Appendix Figure S10A, B, D, and E). The unfolding of the linker enables the tail sheath initiator proteins to maintain the same contacts within the hexamer and with wedge modules as in the baseplate of the extended tail (Appendix Figure S10G, H). The handshake domain of the tail sheath initiator protein inclines 17° (Appendix Figure S10B, E) and, together with the wedge iris expansion, triggers the contraction of the tail sheath, as discussed below.

### The tail of 812 in extended conformation can bend

Contractile tails of phages differ in their length and mechanical properties. Tail bending up to 90° has been observed for *Herelleviridae* phages infecting *Staphylococcus* (Nováček *et al*, 2016; Klumpp *et al*, 2010), *Listeria* phage A511 (Guerrero-Ferreira *et al*, 2019), and phage Milano (Sonani *et al*, 2024) from genus *Schmittlotzvirus* of an unclassified family. In contrast, the ∼100-nm-long tails of *Straboviridae* phages, represented by T4, are rigid (Kostyuchenko *et al*, 2005). The flexibility of the longer tails was speculated to be an adaptation to withstand mechanical forces without breaking (Nováček *et al*, 2016; Zinke *et al*, 2022). To explain the ability of the 812 tail to bend without breaking, we determined the structure of the extended 812 tail to a resolution of 4.2 Å (Fig. 6H, Appendix Figure S7G, Appendix Table S1). Both tail tube and tail sheath proteins are arranged as six-start helices with a rise of 39.2 Å and twist of 21.2° (Fig. 6H).

The tail tube protein of 812 has an eight-stranded β-barrel fold with a protruding stacking loop, characteristic of tail tube proteins of long-tailed phages and CISs (Zinke *et al*, 2022) (Appendix Figure S18A). Whereas the outer surface of the 812 tail tube features roughly equal distribution of positively and negatively charged residues, the inner surface of the tail tube is negatively charged, probably to facilitate the passage of the phage DNA during ejection (Appendix Figure S18B, C). The ability of phage Milano to bend was attributed to the missing loop in the β-barrel domain of the tail tube protein (Sonani *et al*, 2024). The phage Milano tail tube straightening, after contraction, was speculated to be caused by the negative electrostatic potential of its outer surface (Sonani *et al*, 2024). However, since the tail tube of 812 contains the loop between the β-strands 6 and 7 (Appendix Figure S18D) and its outer surface charges are balanced (Appendix Figure S18B), its bending and straightening are likely enabled by different mechanisms.

The sheath protein of 812 can be divided into domains I-IV, according to the convention established for the tail sheath protein (gp18) of phage T4 (Leiman & Shneider, 2012) (Appendix Figure S19A). In an extended tail, the domains I and II of the 812 tail sheath proteins interact with tail tube proteins and mediate contacts between tail sheath proteins (Appendix Figure S19B, C). Domains III and IV are positioned successively next to domain II at the tail surface (Appendix Figure S19B). The two-stranded β-sheet of domain I is augmented by accepting the N- and C-termini of tail sheath proteins from a disc positioned closer to the phage head (Appendix Figure S19C). By mediating the handshake interactions, the domain I interconnects tail sheath proteins from successive discs. These interactions are conserved in the tail sheath structures of numerous phages and CISs (Taylor *et al*, 2016; Ge *et al*, 2020; Jiang *et al*, 2019; Xu *et al*, 2022; Li *et al*, 2023; Wang *et al*, 2023b; Yang *et al*, 2023a, 2023b; Yu *et al*, 2024; Sonani *et al*, 2024; Desfosses *et al*, 2019; Weiss *et al*, 2022; Kudryashev *et al*, 2015; Ouyang *et al*, 2022). The domain II of the tail sheath protein is formed by six β-strands surrounded by eight α-helices, whereas domains III and IV each contain a beta-sandwich flanked by an α-helix (Appendix Figure S19A). Domain III contacts domain IV of the tail sheath protein from the disc located closer to the baseplate (Appendix Figure S19D). Additionally, domain IV interacts with domain II of the neighboring tail sheath protein from the same disc (Appendix Figure S19D). The domain IV of the T4 gp18 is not involved in inter-subunit contacts (Aksyuk *et al*, 2009); however, the bulkier domain III of the T4 tail sheath protein (168 residues) interacts with another tail sheath protein from the disc closer to the phage head (Aksyuk *et al*, 2009). We speculate that interactions of domains III and IV of the 812 tail sheath proteins allow a spring-like movement of the subunits relative to each other, which enables 812 tail bending (Appendix Figure S19E).

### Contraction of the tail sheath

In phages with contractile tails and CISs, the rearrangement of the baseplate induced by binding to a target cell wall triggers the contraction of the tail sheath, which transforms from the extended state to a lower energy contracted state (Maghsoodi *et al*, 2019). The released energy of the tail sheath contraction is used to drive the tail tube through the host cell wall (Ge *et al*, 2020; Li *et al*, 2023; Yu *et al*, 2024; Nováček *et al*, 2016; Guerrero-Ferreira *et al*, 2019). The contracted tail sheath of 812, which was reconstructed to a resolution of 3.2 Å, broadens to 276 Å in diameter (Fig. 6I, Appendix Figure S7J, Appendix Table S1). The sheath broadening is coupled to a change in the rise and twist of the six-start helix formed by the tail sheath proteins to 18.8 Å and 30.6°, respectively (Fig. 6I). The successive discs of the contracted tail sheath are brought closer to each other due to the tilting of domain I by 47° compared to its extended conformation (Fig. 6J) while maintaining the inter-subunit handshake interactions among domains I from successive discs (Appendix Figure S19F, G). During contraction, the individual domains of the tail sheath protein of phage 812, unlike those of T4 and some CISs (Jiang *et al*, 2019; Aksyuk *et al*, 2009; Ge *et al*, 2015), move relative to each other, which enables the formation of the more extensive contacts between the subunits (Appendix Figure S19H). The buried surface area of intersubunit contacts per tail sheath monomer increases from ∼5,500 Å in the extended tail sheath to ∼9,500 Å in the contracted tail sheath. We speculate that the enlarged contacts stabilize the straight structure of the contracted tail sheath, which is less flexible than the extended one.

It has been shown for phages T4 and A511 and R-type diffocin that the contraction of the tail sheath is gradual and occurs as a wave along the tail axis (Guerrero-Ferreira *et al*, 2019; Cai *et al*, 2024; Maghsoodi *et al*, 2019; Moody, 1973). In agreement with the previous *in vitro* observations, we captured particles of 812 with a partially contracted tail sheath attached to the *S. aureus* cell wall (Appendix Figure S16). Furthermore, the cryo-EM reconstructions of baseplate-proximal parts of the tail sheath in the extended and contracted state provide structures of discrete intermediates of contraction (Appendix Figure S1D, S7C). The first three baseplate-proximal tail sheath discs of the extended 812 tail sheath differ in conformation from those in the central part of the extended tail sheath (Appendix Figure S20A, B). The deviation occurs gradually, with the first tail sheath disc being the most affected (Appendix Figure S20B). Similar deviations of baseplate-proximal tail sheath proteins were observed in the R2 pyocin and T6SS (Ge *et al*, 2020; Nazarov *et al*, 2018). The first three sheath discs of 812 have their organization shifted toward that of the contracted tail sheath, which may reduce the activation energy required to initiate the tail sheath contraction (Fraser *et al*, 2021).

In the contracted tail sheath of 812, the structures of the first six baseplate-proximal tail sheath discs deviate from those of the central part of the contracted tail sheath (Appendix Figure S20C, D). Similar to the extended tail sheath, the first contracted sheath disc is the most affected, and the sixth one the least affected (Appendix Figure S20D). The first two baseplate-proximal sheath discs have the rise increased by 5.5 Å relative to the tail sheath disc from the center of the contracted tail sheath (Appendix Figure S20D). The absence of a twist difference may be due to the more extensive contacts between consecutive tail sheath discs than in the extended tail, which is probably incompatible with deviations in the angle of the helical twist. Interactions with the sheath initiator and wedge module proteins likely cause the altered structure of the baseplate-proximal tail sheath discs and directly couple the conformational changes of baseplate proteins with the sheath contraction.

### Changes in the baseplate structure required for cell wall penetration

Phages with contractile tails infecting Gram-positive bacteria employ enzymes that degrade teichoic acid and peptidoglycan to facilitate tail tube penetration through the cell wall (van Dalen *et al*, 2020; Fernandes & São-José, 2018). Phage 812 may use the petal domain of the central spike protein to cleave teichoic acid (Fig. 3B, C); however, the central spike protein needs to be released from the tail channel to enable the genome ejection. We speculate that the conformational change to inner tripods, which disrupts their interaction with weld proteins, triggers the release of the weld and central spike proteins from the baseplate. Dissociation of the weld protein is required to enable the cleaver domains of the hub protein to interact with and degrade the cell wall peptidoglycan (Appendix Figure S9). This indicates that the weld and central spike proteins of 812 are released before the cell wall penetration by the tail tube. In contrast, the central spike proteins of phages T4 and E217, which infect Gram-negative bacteria, are released from baseplates after puncturing the host outer membrane (Li *et al*, 2023; Nishima *et al*, 2011; Wenzel *et al*, 2020).

Relative rotation of consecutive tail sheath discs in the extended 812 tail is 21.2°, whereas that of the contracted tail sheath is 30.6°. Since the tail contains 51 tail sheath discs, as the 812 tail sheath contracts, the tip of the tail tube is pushed through the teichoic acid and peptidoglycan layers while rotating about its axis by 475° (see Methods). The rotation of the hub proteins at the tip of the tail tube can enhance their peptidoglycan degradation activity by bringing them into contact with more substrate. Once the peptidoglycan layer is breached, the clamp domains of the hub proteins can interact with the *S. aureus* cytoplasmic membrane. It was shown that after the penetration of the outer membrane and peptidoglycan layer, gp27 from T4 (a homolog of the hub protein from 812) remains associated with the tip of the tail tube and probably interacts with the ejected tape measure protein to form a pore through the inner membrane of *E. coli* (Hu *et al*, 2015). The tail tube of 812 penetrates 10-30 nm into the *S. aureus* cytoplasm, depending on the thickness of the cell wall, which varies between 50-70 nm (Nováček *et al*, 2016). Therefore, unlike in T4, the hub proteins of 812 are probably not required to form a pore in the *S. aureus* membrane, and their role in genome delivery may be limited to facilitation of the penetration of the cell wall and cytoplasmic membrane by the tail tube tip. Whether the hub and other baseplate core proteins remain associated with the tip of the tail tube after the membrane penetration remains unknown.

### Conclusions

In this study, we employed a combination of cryo-EM, X-ray crystallography, NMR, and structure prediction methods to elucidate the intricate mechanisms employed by phage 812 to penetrate the *S. aureus* cell wall in preparation for transferring its genetic material into the host cytoplasm. While most of the previously structurally characterized phages with contractile tails target Gram-negative bacteria, our focus has been on 812 attacking *S. aureus*, a representative of Gram-positive bacteria, which are comparatively less explored. We show that the baseplate of the phage 812 comprises a core, wedge modules, and baseplate arms that carry RBP1, RBP2, and tripod complexes (Fig. 1A-D). Upon binding to the *S. aureus* cell wall, the baseplate symmetry transitions from threefold to sixfold, which is enabled by a series of conformational changes that propagate from the baseplate periphery to the core (Fig. 6A-C). Based on our results, we propose a model of 812 cell adhesion and penetration (Fig. 7). During the attachment, the putative receptor-binding proteins reorient, and tripod complexes undergo conformational changes to enable host cell surface binding (Fig. 7A, B). Subsequently, the central spike proteins degrade cell wall teichoic acids (Fig. 7A, B). In contrast, phages employ central spike proteins with rigid β-prism or β-helix domains to penetrate the outer membrane of Gram-negative bacteria(Browning *et al*, 2012; Kanamaru *et al*, 2002; Shneider *et al*, 2013). The conformational changes to the tripod complexes of 812 trigger the release of the central spike and weld proteins from the baseplate, which enables the hub proteins to cleave peptidoglycan (Fig. 7C, D) and facilitate penetration of the tail tube through the cell membrane (Fig. 7D, E). Changes in the baseplate arms’ positions relayed through wedge modules to the hexamer of tail sheath initiator proteins cause an expansion of its diameter, triggering a tail sheath contraction (Fig. 7C-E). The structures of the baseplate core, tail sheath, and tail tube proteins of 812 are similar to those of phages and CISs targeting Gram-negative bacteria CISs (Park *et al*, 2018; Ge *et al*, 2020; Weiss *et al*, 2022; Xu *et al*, 2022; Desfosses *et al*, 2019; Jiang *et al*, 2019; Sonani *et al*, 2024; Yang *et al*, 2023a; Li *et al*, 2023; Yu *et al*, 2024; Wang *et al*, 2023a; Taylor *et al*, 2016), indicating that the mechanism triggering the tail sheath contraction is conserved among the phages infecting hosts with distinct cell wall properties. The baseplate-proximal tail sheath discs of the extended 812 particle have an altered structure, which may lower the energy barrier required to initiate the tail sheath contraction. The tail sheath shortens from 200 to 96 nm upon contraction (Nováček *et al*, 2016), pushing the tail tube’s tip through the peptidoglycan layer and plasma membrane 10-30 nm into the cytoplasm, depending on the local thickness of the *S. aureus* cell wall. The hub proteins with putative peptidoglycan-degrading activity are positioned at the tip of the 812 tail tube and may facilitate its penetration through the cell wall. In contrast, T4 baseplate lysozymes are released from the baseplate into the periplasmic peptidoglycan layer of Gram-negative *E. coli* (Kanamaru *et al*, 2002; Arisaka *et al*, 2003). The molecular characterization of the 812 baseplate structure and the mechanism of cell wall binding open up the possibility of a rational design of engineered bacteriophages to target specific bacterial strains.

**Fig. 7.**
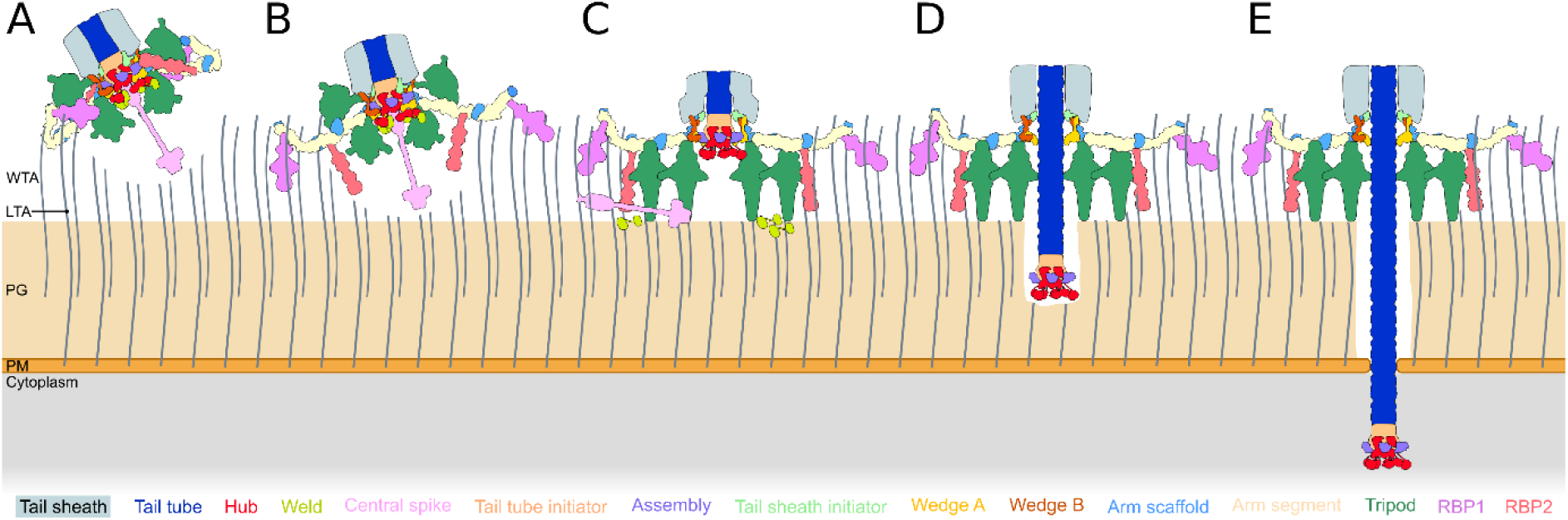
Scheme of 812 cell adhesion and penetration mechanism. **(A)** The baseplate of phage 812 comes in contact with the cell wall of *S. aureus*, trimers of RBP1 bind to teichoic acid, and the petal domains of the central spike proteins cleave teichoic acid below the baseplate. **(B)** Trimers of RBP2 rotate and bind to teichoic acid. **(C)** Tripod complexes undergo a conformational change and anchor the baseplate to the cell wall. Central spike and weld proteins detach from the baseplate. The diameter of the iris formed by wedge modules increases, leading to a conformational change to tail sheath initiator proteins and baseplate-proximal sheath discs. The remaining baseplate core proteins detach from the sheath initiator and wedge module proteins. **(D)** As the contraction of the sheath discs propagates towards the phage head, the tail tube with tail tube initiator, hub, and assembly proteins at its tip is driven through the cell wall. The cleaver domain of the hub protein degrades the peptidoglycan. **(E)** The tail tube penetrates the cytoplasmic membrane, and its tip reaches the cytoplasm. However, it is not known whether the baseplate core proteins remain associated with the tip of the tail tube during the membrane penetration. WTA - wall teichoic acid; LTA - lipoteichoic acid; PG - peptidoglycan; PM - plasma membrane. The scheme is colored according to the list of proteins at the bottom.

## Materials and Methods

### Bacteriophage 812 propagation and purification

For the 812 virion preparation, 150 mL of meat peptone broth (13 g/l nutrient broth, 3 g/l yeast extract, 5 g/l tryptone, pH 7.2) was inoculated with 2.5 ml of the overnight culture and incubated at 37°C with aeration until the culture reached an optical density (OD600) of 0.5-0.6. Next, the culture was supplemented with 2 mM CaCl2 and infected with 15 ml of phage lysate (∼108 PFU/ml). The mixture was incubated at 37°C with shaking at 340 RPM. After 2-3 hours, the first signs of bacterial lysis appeared, and the mixture was placed in the fridge at 4°C overnight. The next day, the lysate was centrifuged for 30 minutes at 5,000 g and 4°C to remove cell debris. The supernatant was collected, filtered through a 0.45 μm filter, and stored at 4°C. Phage particles were pelleted by centrifugation at 54,000g for 2.5 hours at 4°C using a Beckman 50.2 Ti rotor. The supernatant was discarded, and 100 μl of phage buffer (10 mM NaCl, 10 mM CaCl₂, 50 mM Tris-HCl, pH 8) was added to each centrifuge tube. The pellets were left to dissolve overnight at 4°C with shaking. The next day, the resuspended phage sample was loaded onto a cesium chloride step gradient (1.4, 1.45, 1.5, and 1.7 g/ml), prepared in the phage buffer, and centrifuged at 40,000 RPM for 4 hours at 4°C using a Beckman SW41Ti rotor. After centrifugation, the visible bands were collected and dialyzed against phage buffer overnight at 4°C.

### Induction of 812 tail contraction and genome release *in vitro*

To induce contraction of the 812 tail, a previously reported protocol was modified (Fokine *et al*, 2004). The purified phage sample (10^12^ PFU/ml) in phage buffer was supplemented with teichoic acid from *Staphylococcus aureus* (SIGMA) at a concentration of 100 μg/ml, then diluted 10x using a solution of 3M urea in the phage buffer and incubated at 42°C for 45 min. After incubation, the sample was diluted 3x in the phage buffer and treated with Turbonuclease (Abnova) at a final concentration of 3.5 U/ml for 15 min at room temperature. Phage particles were pelleted by centrifugation at 75,000 x *g* at 4°C for 1 h using a 50.2 Ti rotor (Beckman Coulter) and resuspended in the phage buffer.

### Identification of structural proteins of the phage baseplate

The purified 812 phage was resuspended in Laemmli buffer and boiled for 3 min. Subsequently, the sample was treated with Turbonuclease (Abnova) at a final concentration of 2 U/μl for 15 min at room temperature. Proteins from the phage particles were separated using tricine gradient gel electrophoresis. Major protein bands were cut from the gel and used for mass spectrometry analysis. Mass spectrometry data processing and analyses were performed using FlexAnalysis 3.4 and MS BioTools (Bruker Daltonics). Mascot software (Matrix Science, London, UK) was used for sequence searches in exported MS/MS spectra against the National Center for Biotechnology Information database and a local database supplied with the expected protein sequences. The mass tolerance of peptides and MS/MS fragments for MS/MS ion searches was 50 parts per million and 0.5 Da, respectively. The oxidation of methionine and propionyl-amidation of cysteine as optional modifications, and one enzyme mis-cleavage were set for all searches. Peptides with a statistically significant peptide score (P < 0.05) were considered.

### Cryo-EM phage sample preparation, data collection, motion correction, and CTF estimation

Purified 812 (10^11^ PFU/ml) in phage buffer (4 μl), either in the extended or contracted state, was pipetted onto a holey carbon-coated copper grid (R2/1, mesh 200; Quantifoil), blotted and vitrified by plunging into liquid ethane using a Vitrobot Mark IV (FEI). The vitrified sample was transferred to a Titan Krios electron microscope (Thermo Fisher) operated under cryogenic conditions and at an acceleration voltage of 300 kV. The beam was aligned for parallel illumination in NanoProbe mode, and coma-free alignments were performed to remove the residual beam tilt. Micrographs of 812 with extended tail were collected at a magnification of 105,000× using a K3 direct electron detector (Gatan) operating in counting mode, resulting in a calibrated pixel size of 0.8336 Å/pix. Imaging was done under low-dose conditions (total dose 40.8 e-/Å^2^) and with defocus values ranging from −0.5 to −2.0 μm. A two-second exposure was fractioned into 40 frames and saved as a movie. Micrographs of 812 in a contracted state were collected at a magnification of 130,000x using a K2 Summit direct electron detector (Gatan) operating in counting mode, resulting in a calibrated pixel size of 1.057 Å/pix. Imaging was done under low-dose conditions (total dose 42 e-/Å^2^) and with defocus values ranging from −1.0 to −2.4 μm. A seven-second exposure was fractioned into 40 frames and saved as a movie. Automated data acquisitions were performed using the software Serial EM. The movies were corrected for global and local motion during acquisition using MotionCor2 (Zheng *et al*, 2017), and saved as dose-weighted micrographs. Defocus values were estimated from aligned non-dose-weighted micrographs using the program Gctf (Zhang, 2016).

### Reconstruction of the baseplate of phage 812 with extended tail

A total of 444 particles were manually picked from random 1000 micrographs using Relion (Scheres, 2015) and used to train Topaz (Bepler *et al*, 2019). Automated picking with Topaz on 26,731 micrographs yielded 64,874 particle coordinates. Particles were initially extracted into a 1,280 px box and binned 4x (320 px box size, 3.3344 Å/px). Two rounds of 3D multireference classification followed by two rounds of reference-free 2D classification resulted in a selection of 17,393 particles. Particles were 3D refined with no symmetry imposed, and the previously reconstructed native baseplate (EMD-8210), low-pass-filtered to 60 Å, was used as an initial model. Two rounds of subsequent 3D classifications yielded 4,195 particles, which were re-extracted with 2x binning into a 640 px box (1.6672 Å/px). A further round of 3D classification yielded a final set of 3,586 particles. After correction of anisotropic magnification and beamtilt, an asymmetric reconstruction reached a resolution of 8.4 Å after post-processing; imposing C3 symmetry improved the resolution to 5.9 Å after post-processing.

To improve the reconstruction of the core and wedge proteins, 8,470 particles of a 3D refined baseplate were re-extracted with 2× binning into a 640 px box (1.6672 Å/px). After one round of 3D refinement and 3D classification with a mask including the core and wedge modules and no symmetry imposed, particles were assigned to optics groups, and CTF refinements were carried out. A subsequent re-extraction of 8,368 particles to a 640 px box rescaled to a 512 px box (1.042 Å/px) was followed by 3D refinement with C3 symmetry imposed. After one round of CTF refinements and Bayesian polishing, the final map reached the resolution of 3.0 Å after post-processing using a structure-derived mask.

To improve the reconstruction of the baseplate-proximal tail, the 3D refined baseplate map was segmented, and the segment containing the tail sheath initiator protein and the first three disks of tail sheath and tail tube protein was used to calculate the center-of-mass coordinates for targeted re-extraction. Using this offset, 8,919 particles of 3D refined baseplate were re-extracted into a 640 px box and binned 2× to a 320 px box (1.6672 Å/px). Four rounds of asymmetric 3D refinements and classifications reduced the dataset to 4,071 particles. These particles were re-extracted into a 640 px box and rescaled to a 512 px box (1.042 Å/px), then 3D refined with C3 symmetry imposed. After three rounds of CTF refinements and Bayesian polishing, the final map reached the resolution of 3.4 Å after post-processing using a structure-derived mask.

To improve the reconstruction of the lower and upper baseplate arms, the 3D refined baseplate map was segmented and the segments corresponding to the lower and upper arms were used to calculate the center-of-mass coordinates for a modified localized reconstruction protocol (Ilca *et al*, 2015) (github.com/fuzikt/localrec). Using these offsets, 26,748 particles for each arm were re-extracted into a 720 px box and binned 2× to a 360 px box (1.6672 Å/px). One round of 3D refinement and classification produced 22,031 particles for the lower arm, and 19,727 particles for the upper arm; further 3D classifications revealed classes with variable conformations. To address the heterogeneity for each arm, maps of the lower and upper arms were segmented into several subparts, and center-of-mass coordinates were calculated for every subpart. For each arm subpart, one or two rounds of 3D refinement with Blush regularization and 3D classification were performed; when a subpart reconstruction reached ∼4.5 Å resolution, particles were assigned to optics groups and subjected to CTF refinements. Structure-derived masks were used for post-processing to generate the final subpart maps.

In all the above cases, the resolution after post-processing was calculated according to the gold standard FSC = 0.143 criterion. The local resolution for each map was computed using Relion (Kimanius *et al*, 2021).

### Reconstruction of the baseplate of phage 812 with a contracted tail

Out of 15,371 micrographs of phages in the post-contraction state, the first 1,000 micrographs were used to manually pick 7,539 particles using crYOLO (Wagner *et al*, 2019). Particles were then extracted and binned 4x to a 208 px box size (4.228 Å/px) using Relion (Zivanov *et al*, 2020). After a single round of reference-free 2D classification, all aligned particles were re-extracted unbinned to an 832 px box size (1.057 Å/px) with re-centering of the offsets to x,y,z coordinates equal to 0,0,0. Then, particles were imported into Topaz (Bepler *et al*, 2019) with a down-sampling factor of 8. After training the neural network, the positions of particles were predicted using a radius of r = 36 and a picking threshold of t = 0. After the picking coordinates were corrected for the sampling factor, 54,841 particles were extracted using a 960 px box size and binned 4x to a 240 px box size (4.228 Å/px) using Relion (Kimanius *et al*, 2021). After several rounds of reference-free 2D classification, 31,308 particles were subject to asymmetric 3D reconstruction and classification, using the previously reconstructed post-contraction baseplate (EMD-8212) low-pass-filtered to 60 Å as an initial model. The resulting 25,487 particles were re-extracted to a 1280 px box and binned 2x to obtain a 640 px box size (2.114 Å/px). After 3D classification, 21,264 particles were 3D refined, and the final asymmetric map reached the resolution of 6.6 Å after post-processing. Another 3D refinement with C6 symmetry imposed yielded a final map with a resolution of 4.4 Å after post-processing.

To improve the reconstruction of the core, wedge module, and inner tripod proteins, 3D refined post-contraction baseplate particles were re-extracted to a 720 px box (1.057 Å/px), and a 3D refinement was performed using a focused mask encompassing selected regions. The particles were assigned to their respective optics groups, and several rounds of CTF refinements (anisotropic magnification estimation, beam tilt, trefoil, and tetrafoil aberrations estimation, and fitting of the defocus and astigmatism parameters per particle) were performed. Then, two rounds of Bayesian polishing and subsequent 3D refinements and CTF refinements were performed. The final map reached the resolution of 3.1 Å after post-processing. The final post-processing step was also performed using a mask encompassing the first six tail sheath discs, core, wedge module, and inner tripod proteins, and a subunit of the arm segment protein, resulting in a map with a resolution of 4.1 Å after post-processing.

To improve the reconstruction of the post-contraction baseplate arm, the 3D refined post-contraction baseplate map was segmented, and the segment corresponding to the arm was used to calculate the center-of-mass coordinates for a modified localized reconstruction protocol (Ilca *et al*, 2015) (github.com/fuzikt/localrec). Using these offsets, 152,894 particles were re-extracted into a 480 px box and binned 2× to a 240 px box (2.114 Å/px). After subtraction of the surrounding signal, three rounds of asymmetric 3D refinements and classifications were performed to select particles with the most pronounced density belonging to the RBP2 protein. A total of 15,390 particles were re-extracted into a 480 px box unbinned (1.057 Å/px), assigned to optics groups, and subjected to two rounds of 3D and CTF refinements, yielding a final map with a resolution of 4.1 Å after post-processing.

To improve the reconstruction of the arm segment B, bulge domain of the arm scaffold protein, and the top part of the outer tripod proteins, 93,601 particles from early 3D classification of the post-contraction baseplate arm were subjected to two rounds of 3D refinement with Blush regularization and 3D classification using a focused mask encompassing the selected region. The baseplate arm map was segmented, and center-of-mass coordinates were calculated for a subpart including the selected region. Using the coordinate offsets, 56,778 particles were re-extracted into a 256 px box unbinned (1.057 Å/px) and the surrounding signal was subtracted. One round of 3D and CTF refinements yielded a final map with the resolution of 4.4 Å after post-processing.

In all the above cases, the resolution after post-processing was calculated according to the gold standard FSC = 0.143 criterion. The local resolution for each map was computed using Relion (Kimanius *et al*, 2021).

### Reconstruction of the extended tail

The 812 tails were manually boxed using the program e2helixboxer.py from the EMAN2 suite (Tang *et al*, 2007), and the helical segments were extracted in a 360 px box using Relion (Zivanov *et al*, 2020). Tail edges and bent and damaged particles were removed from the dataset by multiple rounds of reference-free 2D classification. A map of the 812 tail in its extended conformation (EMD-4051), low-pass-filtered to 50 Å, was used as an initial model for the 3D refinement of 18,188 particles using a helical refinement protocol with C6 symmetry imposed. The helical refinement and the per-particle CTF refinement were iterated until convergence. The final map reached a resolution of 4.2 Å after post-processing. The resolution after post-processing was calculated according to the gold standard FSC = 0.143 criterion. Local resolution maps were computed using Relion (Zivanov *et al*, 2020).

### Reconstruction of the contracted tail

A total of 106,091 particles were manually picked from 4,833 micrographs using the program e2helixboxer.py from the EMAN2 suite (Tang *et al*, 2007), and the helical segments were extracted using a 400 px box size (1.061 Å/px) in Relion (Zivanov *et al*, 2020). Tail edges and bent and damaged particles were removed from the dataset by multiple rounds of reference-free 2D classification. Then, 85,549 particles were subjected to several rounds of 3D refinement and classification with C6 symmetry imposed. A map of the contracted tail determined previously (EMD-4052) filtered to 50 Å was used as an initial model for the 3D refinement of 69,969 particles using a helical refinement protocol with C6 symmetry imposed. The final map reached an average resolution of 3.0 Å after post-processing. The resolution after post-processing was calculated according to the gold standard FSC = 0.143 criterion. The local resolution map was computed using Relion (Zivanov *et al*, 2020).

### Localized reconstruction of the tail tube from the phage with a contracted tail

Information about particle positions and orientations from the final reconstruction of the post-contraction baseplate was used to extract sub-particle images of the tail tube with a modified localized reconstruction protocol (Ilca *et al*, 2015) (github.com/fuzikt/localrec). The initial set of 25,203 sub-particles centered on the contact region between the tail tube and wedge A proteins was extracted to a 256 px box size (1.057 Å/px) and subjected to several rounds of reference-free 2D classifications in Relion (Zivanov *et al*, 2020). After the initial 3D refinement with imposed C6 symmetry, a single 3D classification was performed. The selected 12,906 particles belonging to the best class were further refined with imposed C6 symmetry and maximum allowed deviations from previous orientations of 10°. The final map reached an average resolution of 3.6 Å after post-processing. The resolution after post-processing was calculated according to the gold standard FSC = 0.143 criterion. The local resolution map was computed using Relion (Zivanov *et al*, 2020).

### Calculation of tail tube rotation during tail sheath contraction

Electron micrographs of the phages with extended tails were used to determine that the tail contains 51 tail sheath discs. The rotation between consecutive discs of tail sheath proteins is 21.1° in the extended tail sheath and 30.6° in the contracted tail. The rotation of the tail tip upon tail contraction was calculated as follows. Taking the first tail sheath disc as a reference point, the rotation of 50 tail sheath discs with a 21.1° twist equals the overall rotation of 1055°. For the contracted tail sheath, the rotation of 50 tail sheath discs with the 30.6° twist equals to the overall rotation of 1530°. The overall rotation of the extended tail sheath was subtracted from the overall rotation of the contracted tail sheath, resulting in a difference of 475°.

### Cryo-EM of 812 attached to *S. aureus* cell wall fragments

*S. aureus* SA 812 was cultured in 100 ml of meat peptone broth (1 g of Lab-Lemco (Difco), 5 g of yeast extract powder L21 (Oxoid), 10 g of peptone L37 (Oxoid), and 5 g of NaCl were dissolved in distilled water to a final volume of 1000 ml, and the pH was adjusted to 7.4 using 10 M NaOH) at 37°C until OD_600_=0.3. Cells were harvested by centrifugation at 3,000 × *g* for 15 min. The resulting pellet was resuspended in 80% isopropanol to a final volume of 10 ml and incubated for 1 h at room temperature. The suspension was centrifuged at 1,000 × *g* at room temperature for 15 min. The resulting supernatant was transferred to another tube, and cell wall fragments were harvested by centrifugation at 20,000 × *g* for 30 min. The resulting pellet was resuspended in 100 µl of the phage buffer and mixed with 812 at MOI 10, relative to the starting bacterial culture. After 5 min of incubation at room temperature, the sample was applied to a holey carbon-coated copper grid for electron microscopy (R2/1, mesh 200; Quantifoil), blotted, and vitrified by plunging into liquid ethane using a Vitrobot Mark IV (FEI), blot force 2, blotting time 2 s. Grids were imaged at a magnification of 17,000 ×, using a Tecnai TF20 operated (FEI) under cryogenic conditions and at an acceleration voltage of 200 kV.

### Production of recombinant cleaver domain of hub protein

The DNA segment encoding the cleaver domain of the hub protein (residues 650-808) was amplified from the genomic DNA of phage 812 by PCR using forward and reverse primers (5’GTAGCCATGGAACAAAGTAGTGGA3’ and 5’TAGGATCCTTTATTTCTTATCGTAAA3’). Using restriction cloning, the amplicon was cloned into the vector pET28a(+) (Novagen/Merck), providing a T7 tag and a His6 tag at the C-terminus of the protein. An *E. coli* expression strain BL21(DE3) was transformed with the construct using a standard heat shock procedure (45 s at 42°C). An overnight culture was used to inoculate 1 l of 2% Luria-Bertani broth (Oxoid) supplemented with kanamycin (50 µg/mL) and cultivated at 37°C until OD_600_=0.5. The protein expression was induced with 0.4 mM IPTG, and was allowed to progress for 4-6 h at 30°C.

For nuclear magnetic resonance spectroscopy (NMR), the cleaver domain was produced in M9 minimal medium supplemented with traces of metals, vitamins, and 2 g/L [^13^C]glucose or 0.5 g/L ^15^NH_4_Cl (Cambridge Isotope Laboratories). *E. coli* BL21 (DE3) was grown to OD_600_ = 0.5, and the gene expression was induced with 0.4 mM IPTG and continued at 30 °C for 12 h. The cultures were harvested, resuspended in the sonication buffer (100 mM Tris-Cl, pH 8, 500 mM NaCl, 4 mM DTT, 10 mM imidazole, 0.1 % TritonX-100), and subjected to sonication (Sartorius Labsonic® M), followed by centrifugation at 14,000 x *g* and 4°C for 45 min. The supernatant was filtered (0.45 µm cut-off, Corning Inc.), loaded into a 5 ml Histrap column (GE Healthcare), pre-equilibrated with the loading buffer, and connected to an AKTA FPLC system (GE Healthcare). After the column was washed with 15 column volumes of the loading buffer (50 mM Tris-Cl, 300 mM NaCl, 20 mM imidazole), gradient elution was performed using the elution buffer (50 mM Tris-Cl, 300 mM NaCl, 500 mM imidazole). Fractions with protein were pooled, concentrated, and loaded onto a HiLoad 16/600 Superdex 75 pg size-exclusion chromatography column (GE Healthcare Life Sciences), followed by an isocratic elution with 100 mM Na-phosphate, pH 6.8, 150 mM NaCl. Purified protein was pooled, and the buffer was exchanged by performing two washes with 20 mM Na-phosphate buffer, pH 6.8, using an Amicon Ultra 15 filter (10 kDa cut-off, Millipore). While exchanging the buffer, the sample was concentrated to 20 mg/ml. For the NMR studies, the protein was concentrated at 35 mg/ml while buffer-exchanged to 25 mM Na-phosphate, pH 6.6, and 100 mM NaCl.

### Zymogram analysis

The cleaver domain of the hub protein after Histrap purification was separated on a conventional 12.5% polyacrylamide gel using SDS-PAGE. Polyacrylamide gels (12,5 %) for zymograms contained 0.5% bacterial cell walls, prepared from *S. aureus* CCM 885 cells, autoclaved at 121 °C for 5 min. Samples of the cleaver domain were mixed with loading buffer (1% SDS, 6% sucrose, 100 mM dithiothreitol, 10 mM Tris, pH 6.8, 0.0625% bromophenol blue) and boiled for 5 min before loading onto SDS-PAGE gels and zymograms. The SDS-gel and zymogram were separated using gel electrophoresis. After the gel electrophoresis, the zymogram was washed for 30 min with distilled water to remove SDS. To renature the cleaver domain, the zymogram was soaked at room temperature in distilled water for 2-4 h, and then for 1 h in the buffer containing 150 mM Na-phosphate, pH 7.0, 0.1% Triton X-100, and 10 mM MgCl_2_. The zymogram was stained for 3 h with 0.1% methylene blue in 0.001% KOH and washed with water. The peptidoglycan-hydrolytic activity of the cleaver domain of the hub protein was detected as a clear zone on a dark blue background.

### NMR structure determination of cleaver domain of hub protein

NMR spectra were recorded at 298 K in an 850 MHz Bruker Avance III spectrometer equipped with a ^1^H/^13^C/^15^N TCI cryogenic probe head with z-axis gradients. The composition of the NMR sample was 1.8 mM uniformly ^13^C-, ^15^N-labeled cleaver domain in a buffer containing 25 mM sodium phosphate, pH 6.6, 100 mM NaCl, 1.5 mM DTT, and 10% D_2_O for the lock. Three sparsely sampled 4D NMR experiments were acquired: 4D HC(CC-TOCSY(CO))NH, 4D ^13^C,^15^N edited HMQC-NOESY-HSQC (HCNH), and 4D ^13^C,^13^C edited HMQC-NOESY-HSQC (HCCH). Sequential and aliphatic side chain assignments were obtained automatically using the 4D-CHAINS algorithm (Evangelidis *et al*, 2018) and checked manually. Backbone dihedral angle restraints were derived from TALOS (Shen *et al*, 2009) using a combination of five kinds (H^N^, H^α^, C^α^, C^β^, N) of chemical shift assignments for each residue in the sequence. NOE cross-peaks from the two 4D NOESY spectra (HCNH and HCCH) were assigned automatically by CYANA 3.0 (Güntert, 2009) in structure calculations with torsion angle dynamics. Unambiguous distance and torsion angle restraints (Appendix Table S3) were used in a water refinement calculation (Linge *et al*, 2003), applying the RECOORD protocol (Nederveen *et al*, 2005). The quality of the NMR-derived structure ensembles was validated using PSVS (Bhattacharya *et al*, 2007).

### Production of recombinant central spike protein

The gene for central spike protein was amplified from the genomic DNA of phage 812 using forward and reverse primers (5’AGATTGGTGGCTCGGATGATTTAAATGTAAAAGGATTAGTTTTA3’ and 5’GAGGAGAGTTAAGACTTATTAAATCTTTTTATATTCATCTAGGTAGTTTGTAAA3’). Compared to the full-length gene, the first 17 nucleotide triplets from the 5’ end of the amplified gene were omitted, as they code for the first 17 residues of the central spike protein, which were predicted to be disordered by the secondary structure prediction using the program Phyre (Kelley *et al*, 2015). The truncated gene was cloned into a vector derived from pET22b, which was modified to provide a His6-SUMO-tag fusion followed by the Ulp1 cleavage site at the N-terminus of the protein. The *E. coli* expression strain BL21(DE3) was transformed with the construct using a standard heat shock procedure (45 s at 42°C). An overnight culture was used to inoculate 2% Luria-Bertani broth (Sigma) supplemented with ampicillin (50 µg/ml). Cells were cultivated at 37°C and 250 rpm until OD_600_=0.5, followed by cultivation at 16°C and 250 rpm for 24 h without the induction of expression, as soluble central spike protein was only produced after non-induced leaky expression. Cells were harvested in a typical yield of 8 g per liter of cell culture, resuspended in 2.5 ml of the loading buffer (50 mM Na/K phosphate, pH 7, 200 mM NaCl) per g of cell pellet, and subjected to chemical disintegration (0.5 % octylthioglucoside, 1 mg/ml lysozyme, 2 U/ml Turbonuclease, 1 mM PMSF, 5 mM TCEP) at 6°C overnight. The lysate was centrifuged at 20,000 x *g* and 8°C for 30 min. The resulting supernatant was filtered (0.45 µm cut-off, Corning Inc.) and loaded into a 5 ml Histrap column (GE Healthcare) pre-equilibrated with the loading buffer and connected to an AKTA FPLC system (GE Healthcare). After the column was washed with 15 column volumes of the loading buffer, gradient elution was performed using the elution buffer (50 mM Na/K phosphate, pH 7.0, 200 mM NaCl, 500 mM imidazole). Eluted fractions containing the central spike protein, as determined by SDS-PAGE analysis using 12.5 % polyacrylamide gels, were pooled and supplemented with 5 mM TCEP before the cleavage of the purification tag overnight at 6°C using the in-house prepared Ulp1 protease. The cleavage solution was desalted using a HiPrep^TM^ 26/10 Desalting column (Cytiva) before reverse affinity chromatography using the 5 ml Histrap column (GE Healthcare). Cleaved protein collected in the flow-through fraction was concentrated (30 kDa cut-off; Amicon® Ultra; Merc Millipore) and loaded into a Superdex 16/600 200 pg column (GE Healthcare) pre-equilibrated with the loading buffer (50 mM Na/K phosphate, pH 7.0, 200 mM NaCl), followed by an isocratic elution. Monodisperse fractions were analyzed by SDS-PAGE and concentrated to a final concentration of 5 mg/ml.

### Cryo-EM reconstruction of recombinant central spike protein

The purified central spike protein sample was diluted in buffer (50 mM Na/K phosphate, pH 7.0, 200 mM NaCl) to a concentration of 0.8 mg/ml and then diluted 10x in water. Four µl of the central spike protein at a concentration of 0.08 mg/ml were applied onto a holey carbon-coated copper grid (R1.2/1.3, mesh 200; Quantifoil) glow-discharged for 60 s from both sides. The excess of the sample was blotted for 1 s with filter paper (Whatman No. 1) using blot force -2 and immediately plunge-frozen in liquid ethane using a Vitrobot Mark IV (FEI). The instrument chamber was maintained at 100 % humidity and at 4°C. Cryo-EM data were collected using a Titan Krios TEM (Thermo Fisher Scientific) operated at 300 kV and equipped with a K2 Summit direct electron detector (Gatan). The beam was aligned for parallel illumination in NanoProbe mode, and the residual beam tilt was removed by coma-free alignments. The software SerialEM was used to acquire the data in the electron counting mode at 130,000x magnification, corresponding to a calibrated pixel size of 1.04 Å/px over a defocus range of -0.3 to -3.0 µm. The exposure of 8 s with a dose rate of 6.6 e^-^/px/s was fractionated into 40 frames (total dose 57 e^-^/Å^2^, 1.425 e^-^/Å^2^ per frame). A total of 3,120 micrographs were collected. For each movie, the drift and gain correction was performed with MotionCor2 (Zheng *et al*, 2017), while the contrast-transfer function was estimated using the program Gctf (Zhang, 2016). crYOLO (Wagner *et al*, 2019) was used to pick particles from micrographs denoised using the program janni (Wagner, 2019). Particles were manually picked from 10 micrographs and used to train the neural network. The resulting 487,160 coordinates were used to extract particles to a 256 px box size and binned 2x to a 128 px box size (2.08 Å/px) using Relion (Zivanov *et al*, 2020). Several rounds of 2D classification sorted out classes of the central spike protein’s C-terminal domains, containing 226,791 particles. These particles were re-extracted and binned to a 128 px box size (2.08 Å/px) using Relion 5.0 beta (github.com/3dem/relion/tree/ver5.0). Three additional rounds of reference-free 2D classification resulted in 135,736 particles, which were used for 3D reconstruction using the initial model generated by the stochastic gradient descent algorithm within Relion. Subsequent 3D classifications did not improve the quality or resolution of the reconstruction. Therefore, all 135,736 particles were re-extracted unbinned to a 256 px box (1.04 Å/px) and assigned to their respective optics groups. Several rounds of CTF refinements, including anisotropic magnification estimation, beam tilt and trefoil aberrations estimation, and fitting of the defocus and astigmatism parameters per micrograph, improved the resolution of the post-processed map to 3.6 Å. Then, two rounds of Bayesian polishing and subsequent 3D refinement with Blush regularization and CTF refinements were performed. The final map reached a resolution of 3.2 Å. The resolution after post-processing was calculated according to the gold standard FSC = 0.143 criterion. Local resolution maps were computed using Relion (Kimanius *et al*, 2021).

### Production of recombinant Arm segment protein

The gene encoding the arm segment protein was cloned into the pET-28b(+) vector (Novagen/Merck), providing a His6 tag at the C-terminus of the protein. An *E. coli* expression strain BL21(DE3) was transformed with the construct using a standard heat shock procedure (45 s at 42°C). An overnight culture was used to inoculate 2% Luria-Bertani broth (Sigma) supplemented with 1 mM MgCl2, 1 % glucose, and ampicillin (50 µg/ml), followed by cultivation at 37°C and 250 rpm until OD_600_=0.5. The expression was induced with 0.4 mM IPTG at 18°C and was allowed to progress for 17 h. Cells were harvested and lysed using the lysis buffer (0.5 % octyl-thioglucoside, 1 mg/ml lysozyme, 2 U/ml Turbonuclease, 1 mM PMSF, 5 mM TCEP) at room temperature for 30 minutes, and subsequently disintegrated with an Emulsiflex C3 (Avestin). After centrifugation at 20,000 x *g* and 8°C for 30 min, the supernatant was filtered (0.45 µm cut-off, Corning Inc.) and loaded into a 5 ml Histrap column (GE Healthcare) pre-equilibrated with the loading buffer (50 mM HEPES, pH 8, 200 mM NaCl, 40 mM imidazole) connected to an AKTA Pure FPLC system (GE Healthcare). The protein was eluted in one step using the elution buffer (50 mM HEPES, pH 8, 200 mM NaCl, 500 mM imidazole). Eluted fractions containing the arm segment protein, detected by SDS-PAGE analysis on 12 % polyacrylamide gels, were loaded into a Superdex 16/600 75 pg column (GE Healthcare) pre-equilibrated with the loading buffer (50 mM HEPES, pH 8, 200 mM NaCl), followed by an isocratic elution. Monodisperse fractions were analyzed by SDS-PAGE and concentrated to a final concentration of 14 mg/ml.

### Crystallization and X-ray structure determination of recombinant arm segment protein

The arm segment protein was crystallized by vapor diffusion using the sitting-drop method at 21°C. The protein sample at a concentration of 14 mg/ml was used to prepare drops with protein to precipitant ratios of 200 nl: 100 nl, 100 nl: 100 nl, and 100 nl: 200 nl in MRC 3-drop 96-well plates using a Dragonfly robot (SPT Labtech) with a range of commercially available screens (Molecular Dimensions, Hampton Research, Qiagen-Nextal). Thick needle-like crystals were observed in the drops containing the solution of 1.6 M LiCl, 0.1 M Tris, pH 7.5, 25 % w/v PEG 4,000. The same crystallization solution was used to prepare hanging drops for vapor diffusion. Crystals were harvested from the hanging drops and flash-vitrified in liquid nitrogen.

Diffraction data were recorded at 100 K on the Proxima-2A beamline at the Soleil synchrotron (Paris, France). Native crystals were soaked in I3C using the I3C phasing kit (Hampton Research), and the dataset was phased using the multiple wavelength anomalous dispersion method. The diffraction data from the crystals soaked with I3C were collected at 12.67 ke^-^V and 5.00 ke^-^V. The structure of the arm segment protein was determined using the program SHELXD (Schneider & Sheldrick, 2002). The native diffraction data recorded from the crystals of the arm segment protein were phased using the program Phaser (McCoy *et al*, 2007), and the structure was determined using the anomalous data as a molecular replacement model. Refinement was performed manually using the program *Coot* (Emsley *et al*, 2010) and automatically in *REFMAC5* (Vagin *et al*, 2004). The structures were validated using *MolProbity* (Williams *et al*, 2018).

### Structure prediction, determination, and refinement using cryo-EM maps

The structures of phage proteins were predicted using the program Alphafold2 (Jumper *et al*, 2021) and fitted into their densities as rigid bodies or by individual domains using the programs Chimera (Pettersen *et al*, 2004) or ChimeraX (Meng *et al*, 2023), and then adjusted interactively using ISOLDE (Croll, 2018). The structures were iteratively improved using the real-space refinement implemented in the *Phenix* package (Liebschner *et al*, 2019), and corrected manually using programs *Coot* (Emsley *et al*, 2010) and ISOLDE (Croll, 2018). The geometry evaluation of models was performed using *MolProbity* (Williams *et al*, 2018).

### Visualization and structure analysis

Chimera (Pettersen *et al*, 2004) and ChimeraX (Meng *et al*, 2023) were used to visualize maps and structures. PISA web server (Krissinel & Henrick, 2007) was used to calculate interface areas. Sequences were analyzed using Clustal Omega (Sievers & Higgins, 2018) available via the EMBL-EBI server (Madeira *et al*, 2022) and visualized with Jalview (Waterhouse *et al*, 2009).

### Construction of the composite maps

Composite maps of pre- and post-contraction baseplates were constructed in ChimeraX using the *volume maximum* command. Given variable resolution and quality of consensus, focused and sub-particle maps, the numeric threshold levels were adjusted individually, and the command option *scaleFactors fromLevels* was used. In case of the pre-contraction baseplate, sub-particle reconstructions of lower (Appendix Figure S1F-K) and upper (Appendix Figure S1M-O) baseplate arms were combined with the overall reconstruction of the lower (Appendix Figure S1E) and upper (Appendix Figure S1L) baseplate arm, respectively. Then, newly constructed maps of the lower and upper arms were combined with reconstructions of the core and wedge module proteins (Appendix Figure S1C) and baseplate-proximal tail (Appendix Figure S1D) to create the final composite map of the pre-contraction baseplate.

In the case of the post-contraction baseplate, sub-particle reconstruction of the arm segment B (Appendix Figure S7F) was combined with the post-processed reconstruction of the baseplate arm (Appendix Figure S7E). To better visualize the density of the RBP2, a refined map (without post-processing) of the baseplate arm was segmented, and the density corresponding to the RBP2 and an unassigned density at the baseplate periphery were extracted. This map was then combined with the baseplate arm map created in the previous step. Finally, the newly constructed map of the baseplate arm was combined with the reconstruction of the core, wedge module, inner tripod, and baseplate-proximal tail sheath proteins (Appendix Figure S7C) and reconstructions of the tail tube (Appendix Figure S7I) stacked onto each other to create the final composite map of the post-contraction baseplate.

## Data and Code Availability

### Data availability

Cryo-EM maps and structure coordinates were deposited into the Electron Microscopy Data Bank (EMDB) and Protein Data Bank (PDB), respectively, with the following accession numbers: Structure of whole baseplate and baseplate-proximal tail of phage with extended tail computed with no symmetry imposed: EMD-55950. Structure of whole baseplate and baseplate-proximal tail of phage with extended tail computed with imposed threefold symmetry: EMD-55951 and PDB-9TIC. Structure of core and wedge module proteins of phage with extended tail computed with imposed threefold symmetry: EMD-55952 and PDB-9TID. Structure of baseplate-proximal tail of phage with extended tail computed with imposed threefold symmetry: EMD-55953 and PDB-9TIE. Structure of lower baseplate arm of phage with extended tail computed with no symmetry imposed: EMD-55954 and PDB-9TIF. Structure of lower baseplate arm (segment A) of phage with extended tail computed with no symmetry imposed: EMD-55955 and PDB-9TIG. Structure of lower baseplate arm (segment B) of phage with extended tail computed with no symmetry imposed: EMD-55956 and PDB-9TIH. Structure of lower baseplate arm (segment C,D) of phage with extended tail computed with no symmetry imposed: EMD-55957 and PDB-9TII. Structure of lower baseplate arm (segment D,E,F) of phage with extended tail computed with no symmetry imposed: EMD-55958 and PDB-9TIJ. Structure of lower baseplate arm (uRBP1-lRBP2) of phage with extended tail computed with no symmetry imposed: EMD-55959 and PDB-9TIK. Structure of lower baseplate arm (lRBP1-uRBP2) of phage with extended tail computed with no symmetry imposed: EMD-55960 and PDB-9TIL. Structure of upper baseplate arm of phage with extended tail computed with no symmetry imposed: EMD-55961 and PDB-9TIM. Structure of upper baseplate arm (segment A) of phage with extended tail computed with no symmetry imposed: EMD-55962 and PDB-9TIN. Structure of upper baseplate arm (segment B) of phage with extended tail computed with no symmetry imposed: EMD-55963 and PDB-9TIO. Structure of upper baseplate arm (segment C,D,E,F) of phage with extended tail computed with no symmetry imposed: EMD-55964 and PDB-9TIP. Composite structure of whole baseplate and baseplate-proximal tail of phage with extended tail: EMD-55977.

Structure of whole baseplate and baseplate-proximal tail of phage with contracted tail computed with no symmetry imposed: EMD-55966. Structure of whole baseplate and baseplate-proximal tail of phage with contracted tail computed with imposed sixfold symmetry: EMD-55967 and PDB-9TIR. Structure of core, wedge module, arm segment, inner tripod, and baseplate-proximal tail sheath proteins of phage with contracted tail computed with imposed sixfold symmetry: EMD-19972 and [PDB-9EUJ](https://doi.org/10.2210/pdb9EUJ/pdb). Structure of core, wedge module, and inner tripod proteins of phage with contracted tail computed with imposed sixfold symmetry: EMD-19973 and [PDB-9EUK](https://www.wwpdb.org/pdb?id=pdb_00009euk). Structure of baseplate arm of phage with contracted tail computed with no symmetry imposed: EMD-55968 and PDB-9TIS. Structure of baseplate arm (segment B) of phage with contracted tail computed with no symmetry imposed: EMD-55969 and PDB-9TIT. Composite structure of whole baseplate, baseplate-proximal tail sheath and tail tube of phage with contracted tail: EMD-55978 and PDB-9TIW.

Structure of extended tail computed with imposed sixfold symmetry: EMD-50093 and [PDB-9F04](https://www.wwpdb.org/pdb?id=pdb_00009f04). Structure of tail tube of phage with contracted tail computed with imposed sixfold symmetry: EMD-50094 and [PDB-9F05](https://www.wwpdb.org/pdb?id=pdb_00009f05). Structure of contracted tail sheath computed with imposed sixfold symmetry: EMD-50095 and [PDB-9F06](https://www.wwpdb.org/pdb?id=pdb_00009f06). Structure of central spike protein (knob and petal domains) computed with imposed threefold symmetry: [EMD-19974](https://www.wwpdb.org/pdb?id=pdb_00009f06) and [PDB-9EUL](https://www.wwpdb.org/pdb?id=pdb_00009eul).

The NMR structure of the cleaver domain of the hub protein was deposited to PDB under the accession number 9EUM. The X-ray structure of the arm segment protein was deposited to PDB under the accession number 9FKO.

### Code availability

All custom scripts used during the single-particle data reconstruction are available at https://github.com/fuzikt/starpy.

## Acknowledgments

We gratefully acknowledge the core facilities Cryo-electron microscopy and tomography, Biomolecular Interactions and Crystallography, Proteomics, and Josef Dadok National NMR Centre of CIISB, Instruct-CZ Centre, supported by MEYS CR (LM2023042) and European Regional Development Fund-Project „UP CIISB“ (No. CZ.02.1.01/0.0/0.0/18_046/0015974). We gratefully acknowledge support from the National Institute of Virology and Bacteriology (Program EXCELES, Project ID No. LX22NPO5103) - Funded by the European Union - Next Generation EU and the project New Technologies for Translational Research in Pharmaceutical Sciences/NETPHARM, project ID OP JAC CZ.02.01.01/00/22_008/0004607, which is co-funded by the European Union. This work received funding from ERC Consolidator Grant No. 101043452 to P.P. Computational resources were provided by the e-INFRA CZ project (ID:90254), suppoted by the Ministry of Education, Youth and Sports of the Czech Republic.

## References

Aksyuk AA, Leiman PG, Kurochkina LP, Shneider MM, Kostyuchenko VA, Mesyanzhinov VV & Rossmann MG (2009) The tail sheath structure of bacteriophage T4: a molecular machine for infecting bacteria. EMBO J 28: 821–829

Arisaka F, Kanamaru S, Leiman P & Rossmann MG (2003) The tail lysozyme complex of bacteriophage T4. The International Journal of Biochemistry & Cell Biology 35: 16–21

Bae B, Ohene-Adjei S, Kocherginskaya S, Mackie RI, Spies MA, Cann IKO & Nair SK (2008) Molecular Basis for the Selectivity and Specificity of Ligand Recognition by the Family 16 Carbohydrate-binding Modules from Thermoanaerobacterium polysaccharolyticum ManA. Journal of Biological Chemistry 283: 12415–12425

Barylski J, Enault F, Dutilh BE, Schuller MB, Edwards RA, Gillis A, Klumpp J, Knezevic P, Krupovic M, Kuhn JH, et al (2020) Analysis of Spounaviruses as a Case Study for the Overdue Reclassification of Tailed Phages. Systematic Biology 69: 110–123

Bateman A & Rawlings ND (2003) The CHAP domain: a large family of amidases including GSP amidase and peptidoglycan hydrolases. Trends in Biochemical Sciences 28: 234–237

Becker SC, Foster-Frey J & Donovan DM (2008) The phage K lytic enzyme LysK and lysostaphin act synergistically to kill MRSA. FEMS Microbiology Letters 287: 185–191

Bepler T, Morin A, Rapp M, Brasch J, Shapiro L, Noble AJ & Berger B (2019) Positive-unlabeled convolutional neural networks for particle picking in cryo-electron micrographs. Nat Methods 16: 1153–1160

Bhardwaj A, Walker-Kopp N, Casjens SR & Cingolani G (2009) An Evolutionarily Conserved Family of Virion Tail Needles Related to Bacteriophage P22 gp26: Correlation between Structural Stability and Length of the α-Helical Trimeric Coiled Coil. Journal of Molecular Biology 391: 227–245

Bhattacharya A, Tejero R & Montelione GT (2007) Evaluating protein structures determined by structural genomics consortia. Proteins 66: 778–795

Böth D, Steiner EM, Izumi A, Schneider G & Schnell R (2014) RipD (Rv1566c) from *Mycobacterium tuberculosis*: adaptation of an NlpC/p60 domain to a non-catalytic peptidoglycan-binding function. Biochemical Journal 457: 33–41

Botka T, Pantůček R, Mašlaňová I, Benešík M, Petráš P, Růžičková V, Havlíčková P, Varga M, Žemličková H, Koláčková I, et al (2019) Lytic and genomic properties of spontaneous host-range Kayvirus mutants prove their suitability for upgrading phage therapeutics against staphylococci. Sci Rep 9: 5475

Bradshaw WJ, Kirby JM, Thiyagarajan N, Chambers CJ, Davies AH, Roberts AK, Shone CC & Acharya KR (2014) The structure of the cysteine protease and lectin-like domains of Cwp84, a surface layer-associated protein from *Clostridium difficile*. Acta Crystallogr D Biol Crystallogr 70: 1983–1993

Browning C, Shneider MM, Bowman VD, Schwarzer D & Leiman PG (2012) Phage Pierces the Host Cell Membrane with the Iron-Loaded Spike. Structure 20: 326–339

Buist G, Steen A, Kok J & Kuipers OP (2008) LysM, a widely distributed protein motif for binding to (peptido)glycans. Mol Microbiol 68: 838–847

Büttner CR, Wu Y, Maxwell KL & Davidson AR (2016) Baseplate assembly of phage Mu: Defining the conserved core components of contractile-tailed phages and related bacterial systems. Proc Natl Acad Sci USA 113: 10174–10179

Cai X, He Y, Yu I, Imani A, Scholl D, Miller JF & Zhou ZH (2024) Atomic structures of a bacteriocin targeting Gram-positive bacteria. Nat Commun 15: 7057

Cid M, Pedersen HL, Kaneko S, Coutinho PM, Henrissat B, Willats WGT & Boraston AB (2010) Recognition of the Helical Structure of β-1,4-Galactan by a New Family of Carbohydrate-binding Modules. Journal of Biological Chemistry 285: 35999–36009

Croll TI (2018) *ISOLDE*: a physically realistic environment for model building into low-resolution electron-density maps. Acta Crystallogr D Struct Biol 74: 519–530

van Dalen R, Peschel A & van Sorge NM (2020) Wall Teichoic Acid in Staphylococcus aureus Host Interaction. Trends in Microbiology 28: 985–998

Desfosses A, Venugopal H, Joshi T, Felix J, Jessop M, Jeong H, Hyun J, Heymann JB, Hurst MRH, Gutsche I, et al (2019) Atomic structures of an entire contractile injection system in both the extended and contracted states. Nat Microbiol 4: 1885–1894

Eck MJ & Sprang SR (1989) The Structure of Tumor Necrosis Factor-α at 2.6 Å Resolution. Journal of Biological Chemistry 264: 17595–17605

Emsley P, Lohkamp B, Scott WG & Cowtan K (2010) Features and development of *Coot*. Acta Crystallogr D Biol Crystallogr 66: 486–501

Evangelidis T, Nerli S, Nováček J, Brereton AE, Karplus PA, Dotas RR, Venditti V, Sgourakis NG & Tripsianes K (2018) Automated NMR resonance assignments and structure determination using a minimal set of 4D spectra. Nat Commun 9: 384

Farenc C, Spinelli S, Vinogradov E, Tremblay D, Blangy S, Sadovskaya I, Moineau S & Cambillau C (2014) Molecular Insights on the Recognition of a Lactococcus lactis Cell Wall Pellicle by the Phage 1358 Receptor Binding Protein. J Virol 88: 7005–7015

Fernandes S & São-José C (2018) Enzymes and Mechanisms Employed by Tailed Bacteriophages to Breach the Bacterial Cell Barriers. Viruses 10: 396

Fokine A, Chipman PR, Leiman PG, Mesyanzhinov VV, Rao VB & Rossmann MG (2004) Molecular architecture of the prolate head of bacteriophage T4. Proc Natl Acad Sci USA 101: 6003–6008

Fraser A, Prokhorov NS, Jiao F, Pettitt BM, Scheuring S & Leiman PG (2021) Quantitative description of a contractile macromolecular machine. Sci Adv 7: eabf9601

Ge P, Scholl D, Leiman PG, Yu X, Miller JF & Zhou ZH (2015) Atomic structures of a bactericidal contractile nanotube in its pre- and postcontraction states. Nat Struct Mol Biol 22: 377–382

Ge P, Scholl D, Prokhorov NS, Avaylon J, Shneider MM, Browning C, Buth SA, Plattner M, Chakraborty U, Ding K, et al (2020) Action of a minimal contractile bactericidal nanomachine. Nature 580: 658–662

Gillespie JJ, Phan IQH, Scheib H, Subramanian S, Edwards TE, Lehman SS, Piitulainen H, Sayeedur Rahman M, Rennoll-Bankert KE, Staker BL, et al (2015) Structural Insight into How Bacteria Prevent Interference between Multiple Divergent Type IV Secretion Systems. mBio 6: e01867–15

Grunenwald CM, Bennett MR & Skaar EP (2018) Nonconventional Therapeutics against Staphylococcus aureus. Microbiol Spectr 6

Guerrero-Ferreira RC, Hupfeld M, Nazarov S, Taylor NM, Shneider MM, Obbineni JM, Loessner MJ, Ishikawa T, Klumpp J & Leiman PG (2019) Structure and transformation of bacteriophage A511 baseplate and tail upon infection of *Listeria* cells. EMBO J 38

Güntert P (2009) Automated structure determination from NMR spectra. Eur Biophys J 38: 129–143

Habann M, Leiman PG, Vandersteegen K, Van den Bossche A, Lavigne R, Shneider MM, Bielmann R, Eugster MR, Loessner MJ & Klumpp J (2014) Listeria phage A511, a model for the contractile tail machineries of SPO1-related bacteriophages. Molecular Microbiology 92: 84–99

Hu B, Margolin W, Molineux IJ & Liu J (2015) Structural remodeling of bacteriophage T4 and host membranes during infection initiation. Proceedings of the National Academy of Sciences 112: E4919–E4928

Ilca SL, Kotecha A, Sun X, Poranen MM, Stuart DI & Huiskonen JT (2015) Localized reconstruction of subunits from electron cryomicroscopy images of macromolecular complexes. Nat Commun 6: 8843

Jiang F, Li N, Wang X, Cheng J, Huang Y, Yang Y, Yang J, Cai B, Wang Y-P, Jin Q, et al (2019) Cryo-EM Structure and Assembly of an Extracellular Contractile Injection System. Cell 177: 370–383.e15

Jumper J, Evans R, Pritzel A, Green T, Figurnov M, Ronneberger O, Tunyasuvunakool K, Bates R, Žídek A, Potapenko A, et al (2021) Highly accurate protein structure prediction with AlphaFold. Nature 596: 583–589

Kanamaru S, Leiman PG, Kostyuchenko VA, Chipman PR, Mesyanzhinov VV, Arisaka F & Rossmann MG (2002) Structure of the cell-puncturing device of bacteriophage T4. Nature 415: 553–557

Kelley LA, Mezulis S, Yates CM, Wass MN & Sternberg MJE (2015) The Phyre2 web portal for protein modeling, prediction and analysis. Nat Protoc 10: 845–858

Kimanius D, Dong L, Sharov G, Nakane T & Scheres SHW (2021) New tools for automated cryo-EM single-particle analysis in RELION-4.0. Biochemical Journal 478: 4169–4185

Kizziah JL, Manning KA, Dearborn AD & Dokland T (2020) Structure of the host cell recognition and penetration machinery of a Staphylococcus aureus bacteriophage. PLoS Pathog 16: e1008314

Klumpp J, Lavigne R, Loessner MJ & Ackermann H-W (2010) The SPO1-related bacteriophages. Arch Virol 155: 1547–1561

Kostyuchenko VA, Chipman PR, Leiman PG, Arisaka F, Mesyanzhinov VV & Rossmann MG (2005) The tail structure of bacteriophage T4 and its mechanism of contraction. Nat Struct Mol Biol 12: 810–813

Krissinel E & Henrick K (2007) Inference of Macromolecular Assemblies from Crystalline State. Journal of Molecular Biology 372: 774–797

Krusche J, Beck C, Lehmann E, Gerlach D, Daiber E, Mayer C, Müller J, Onallah H, Würstle S, Wolz C, et al (2025) Characterization and host range prediction of Staphylococcus aureus phages through receptor-binding protein analysis. Cell Reports 44: 115369

Kudryashev M, Wang RY-R, Brackmann M, Scherer S, Maier T, Baker D, DiMaio F, Stahlberg H, Egelman EH & Basler M (2015) Structure of the Type VI Secretion System Contractile Sheath. Cell 160: 952–962

Kwiecinski JM & Horswill AR (2020) Staphylococcus aureus bloodstream infections: pathogenesis and regulatory mechanisms. Curr Opin Microbiol 53: 51–60

Lee JH, Yang S-T, Rho S-H, Im YJ, Kim SY, Kim YR, Kim M-K, Kang GB, Kim JI, Rhee JH, et al (2006) Crystal Structure and Functional Studies Reveal that PAS Factor from Vibrio vulnificus is a Novel Member of the Saposin-fold Family. Journal of Molecular Biology 355: 491–500

Legrand P, Collins B, Blangy S, Murphy J, Spinelli S, Gutierrez C, Richet N, Kellenberger C, Desmyter A, Mahony J, et al (2016) The Atomic Structure of the Phage Tuc2009 Baseplate Tripod Suggests that Host Recognition Involves Two Different Carbohydrate Binding Modules. mBio 7

Leiman PG, Basler M, Ramagopal UA, Bonanno JB, Sauder JM, Pukatzki S, Burley SK, Almo SC & Mekalanos JJ (2009) Type VI secretion apparatus and phage tail-associated protein complexes share a common evolutionary origin. Proc Natl Acad Sci USA 106: 4154–4159

Leiman PG & Shneider MM (2012) Contractile Tail Machines of Bacteriophages. In Viral Molecular Machines, Rossmann MG & Rao VB (eds) pp 93–114. Boston, MA: Springer US

Leiman PG, Shneider MM, Kostyuchenko VA, Chipman PR, Mesyanzhinov VV & Rossmann MG (2003) Structure and Location of Gene Product 8 in the Bacteriophage T4 Baseplate. Journal of Molecular Biology 328: 821–833

Li F, Hou C-FD, Lokareddy RK, Yang R, Forti F, Briani F & Cingolani G (2023) High-resolution cryo-EM structure of the Pseudomonas bacteriophage E217. Nat Commun 14: 4052

Liebschner D, Afonine PV, Baker ML, Bunkóczi G, Chen VB, Croll TI, Hintze B, Hung L-W, Jain S, McCoy AJ, et al (2019) Macromolecular structure determination using X-rays, neutrons and electrons: recent developments in *Phenix*. Acta Crystallogr D Struct Biol 75: 861–877

Linge JP, Williams MA, Spronk CAEM, Bonvin AMJJ & Nilges M (2003) Refinement of protein structures in explicit solvent. Proteins 50: 496–506

Łobocka M, Hejnowicz MS, Dąbrowski K, Gozdek A, Kosakowski J, Witkowska M, Ulatowska MI, Weber-Dąbrowska B, Kwiatek M, Parasion S, et al (2012) Genomics of Staphylococcal Twort-like Phages - Potential Therapeutics of the Post-Antibiotic Era. In Advances in Virus Research pp 143–216. Elsevier

Madeira F, Pearce M, Tivey ARN, Basutkar P, Lee J, Edbali O, Madhusoodanan N, Kolesnikov A & Lopez R (2022) Search and sequence analysis tools services from EMBL-EBI in 2022. Nucleic Acids Res 50: W276–W279

Maghsoodi A, Chatterjee A, Andricioaei I & Perkins NC (2019) How the phage T4 injection machinery works including energetics, forces, and dynamic pathway. Proc Natl Acad Sci USA 116: 25097–25105

Marín-Arraiza L, Roa-Eguiara A, Pape T, Sofos N, Hendriks IA, Lund Nielsen M, Steiner-Rebrova EM & Taylor NMI (2025) Structural characterization of an extracellular contractile injection system from Photorhabdus luminescens in extended and contracted states. Nat Commun 16: 9327

Maxwell KL, Fatehi Hassanabad M, Chang T, Pirani N, Bona D, Edwards AM & Davidson AR (2013) Structural and Functional Studies of gpX of Escherichia coli Phage P2 Reveal a Widespread Role for LysM Domains in the Baseplates of Contractile-Tailed Phages. J Bacteriol 195: 5461–5468

McCabe O, Spinelli S, Farenc C, Labbé M, Tremblay D, Blangy S, Oscarson S, Moineau S & Cambillau C (2015) The targeted recognition of *L actococcus lactis* phages to their polysaccharide receptors: Receptor binding to lactococcal phage 1358. Molecular Microbiology 96: 875–886

McCoy AJ, Grosse-Kunstleve RW, Adams PD, Winn MD, Storoni LC & Read RJ (2007) *Phaser* crystallographic software. Journal of Applied Crystallography 40: 658–674

Meng EC, Goddard TD, Pettersen EF, Couch GS, Pearson ZJ, Morris JH & Ferrin TE (2023) UCSF CHIMERAX: Tools for structure building and analysis. Protein Science 32: e4792

Moody MF (1973) Sheath of bacteriophage T4. Journal of Molecular Biology 80: 613–635

Myers CL, Li FKK, Koo B-M, El-Halfawy OM, French S, Gross CA, Strynadka NCJ & Brown ED (2016) Identification of Two Phosphate Starvation-induced Wall Teichoic Acid Hydrolases Provides First Insights into the Degradative Pathway of a Key Bacterial Cell Wall Component. Journal of Biological Chemistry 291: 26066–26082

Nazarov S, Schneider JP, Brackmann M, Goldie KN, Stahlberg H & Basler M (2018) Cryo-EM reconstruction of Type VI secretion system baseplate and sheath distal end. The EMBO Journal 37: e97103

Nederveen AJ, Doreleijers JF, Vranken W, Miller Z, Spronk CAEM, Nabuurs SB, Güntert P, Livny M, Markley JL, Nilges M, et al (2005) RECOORD: A recalculated coordinate database of 500+ proteins from the PDB using restraints from the BioMagResBank. Proteins 59: 662–672

Nishima W, Kanamaru S, Arisaka F & Kitao A (2011) Screw Motion Regulates Multiple Functions of T4 Phage Protein Gene Product 5 during Cell Puncturing. J Am Chem Soc 133: 13571–13576

Nobrega FL, Vlot M, De Jonge PA, Dreesens LL, Beaumont HJE, Lavigne R, Dutilh BE & Brouns SJJ (2018) Targeting mechanisms of tailed bacteriophages. Nat Rev Microbiol 16: 760–773

North OI, Sakai K, Yamashita E, Nakagawa A, Iwazaki T, Büttner CR, Takeda S & Davidson AR (2019) Phage tail fibre assembly proteins employ a modular structure to drive the correct folding of diverse fibres. Nat Microbiol 4: 1645–1653

Nováček J, Šiborová M, Benešík M, Pantůček R, Doškař J & Plevka P (2016) Structure and genome release of Twort-like Myoviridae phage with a double-layered baseplate. Proc Natl Acad Sci USA 113: 9351–9356

Ouyang R, Costa AR, Cassidy CK, Otwinowska A, Williams VCJ, Latka A, Stansfeld PJ, Drulis-Kawa Z, Briers Y, Pelt DM, et al (2022) High-resolution reconstruction of a Jumbo-bacteriophage infecting capsulated bacteria using hyperbranched tail fibers. Nat Commun 13: 7241

Pantůček R, Rosypalová A, Doškař J, Kailerová J, Růžičková V, Borecká P, Snopková Š, Horváth R, Götz F & Rosypal S (1998) The Polyvalent Staphylococcal Phage φ812:Its Host-Range Mutants and Related Phages. Virology 246: 241–252

Park Y-J, Lacourse KD, Cambillau C, DiMaio F, Mougous JD & Veesler D (2018) Structure of the type VI secretion system TssK–TssF–TssG baseplate subcomplex revealed by cryo-electron microscopy. Nat Commun 9: 5385

Paul VD, Rajagopalan SS, Sundarrajan S, George SE, Asrani JY, Pillai R, Chikkamadaiah R, Durgaiah M, Sriram B & Padmanabhan S (2011) A novel bacteriophage Tail-Associated Muralytic Enzyme (TAME) from Phage K and its development into a potent antistaphylococcal protein. BMC Microbiology 11: 226

Petrovic Fabijan A, Lin RCY, Ho J, Maddocks S, Ben Zakour NL, Iredell JR, Westmead Bacteriophage Therapy Team, Khalid A, Venturini C, Chard R, et al (2020) Safety of bacteriophage therapy in severe Staphylococcus aureus infection. Nat Microbiol 5: 465–472

Pettersen EF, Goddard TD, Huang CC, Couch GS, Greenblatt DM, Meng EC & Ferrin TE (2004) UCSF Chimera—A visualization system for exploratory research and analysis. J Comput Chem 25: 1605–1612

Sanz-Gaitero M, Keary R, Garcia-Doval C, Coffey A & van Raaij MJ (2014) Crystal structure of the lytic CHAPK domain of the endolysin LysK from Staphylococcus aureus bacteriophage K. Virology Journal 11: 133

Schaechter M (2009) Encyclopedia of microbiology 3rd ed. Amsterdam Boston: Elsevier/Academic Press

Scheres SHW (2015) Semi-automated selection of cryo-EM particles in RELION-1.3. Journal of Structural Biology 189: 114–122

Schneider TR & Sheldrick GM (2002) Substructure solution with *SHELXD*. Acta Crystallogr D Biol Crystallogr 58: 1772–1779

Sciara G, Bebeacua C, Bron P, Tremblay D, Ortiz-Lombardia M, Lichière J, van Heel M, Campanacci V, Moineau S & Cambillau C (2010) Structure of lactococcal phage p2 baseplate and its mechanism of activation. Proc Natl Acad Sci USA 107: 6852–6857

Shen Y, Delaglio F, Cornilescu G & Bax A (2009) TALOS+: a hybrid method for predicting protein backbone torsion angles from NMR chemical shifts. J Biomol NMR 44: 213–223

Shi L, Liu J-F, An X-M & Liang D-C (2008) Crystal structure of glycerophosphodiester phosphodiesterase (GDPD) from *Thermoanaerobacter tengcongensis*, a metal ion-dependent enzyme: Insight into the catalytic mechanism: Crystal Structure of GDPD. Proteins 72: 280–288

Shneider MM, Buth SA, Ho BT, Basler M, Mekalanos JJ & Leiman PG (2013) PAAR-repeat proteins sharpen and diversify the type VI secretion system spike. Nature 500: 350–353

Šiborová M, Füzik T, Procházková M, Nováček J, Benešík M, Nilsson AS & Plevka P (2022) Tail proteins of phage SU10 reorganize into the nozzle for genome delivery. Nat Commun 13: 5622

Sievers F & Higgins DG (2018) Clustal Omega for making accurate alignments of many protein sequences. Protein Science 27: 135–145

Sonani RR, Palmer LK, Esteves NC, Horton AA, Sebastian AL, Kelly RJ, Wang F, Kreutzberger MAB, Russell WK, Leiman PG, et al (2024) An extensive disulfide bond network prevents tail contraction in Agrobacterium tumefaciens phage Milano. Nat Commun 15: 756

Šulák O, Cioci G, Delia M, Lahmann M, Varrot A, Imberty A & Wimmerová M (2010) A TNF-like Trimeric Lectin Domain from *Burkholderia cenocepacia* with Specificity for Fucosylated Human Histo-Blood Group Antigens. Structure 18: 59–72

Takeuchi I, Osada K, Azam AH, Asakawa H, Miyanaga K & Tanji Y (2016) The Presence of Two Receptor-Binding Proteins Contributes to the Wide Host Range of Staphylococcal Twort-Like Phages. Appl Environ Microbiol 82: 5763–5774

Tang G, Peng L, Baldwin PR, Mann DS, Jiang W, Rees I & Ludtke SJ (2007) EMAN2: an extensible image processing suite for electron microscopy. J Struct Biol 157: 38–46

Taylor NMI, Prokhorov NS, Guerrero-Ferreira RC, Shneider MM, Browning C, Goldie KN, Stahlberg H & Leiman PG (2016) Structure of the T4 baseplate and its function in triggering sheath contraction. Nature 533: 346–352

Taylor NMI, van Raaij MJ & Leiman PG (2018) Contractile injection systems of bacteriophages and related systems. Molecular Microbiology 108: 6–15

Uyttebroek S, Chen B, Onsea J, Ruythooren F, Debaveye Y, Devolder D, Spriet I, Depypere M, Wagemans J, Lavigne R, et al (2022) Safety and efficacy of phage therapy in difficult-to-treat infections: a systematic review. The Lancet Infectious Diseases 22: e208–e220

Vagin AA, Steiner RA, Lebedev AA, Potterton L, McNicholas S, Long F & Murshudov GN (2004) *REFMAC*5 dictionary: organization of prior chemical knowledge and guidelines for its use. Acta Crystallogr D Biol Crystallogr 60: 2184–2195

Vandersteegen K, Mattheus W, Ceyssens P-J, Bilocq F, De Vos D, Pirnay J-P, Noben J-P, Merabishvili M, Lipinska U, Hermans K, et al (2011) Microbiological and Molecular Assessment of Bacteriophage ISP for the Control of Staphylococcus aureus. PLoS ONE 6: e24418

Veesler D & Cambillau C (2011) A Common Evolutionary Origin for Tailed-Bacteriophage Functional Modules and Bacterial Machineries. Microbiol Mol Biol Rev 75: 423–433

Venturini C, Petrovic Fabijan A, Fajardo Lubian A, Barbirz S & Iredell J (2022) Biological foundations of successful bacteriophage therapy. EMBO Mol Med 14

Wagner T (2019) MPI-Dortmund/sphire-janni: JANNI 0.0.5.

Wagner T, Merino F, Stabrin M, Moriya T, Antoni C, Apelbaum A, Hagel P, Sitsel O, Raisch T, Prumbaum D, et al (2019) SPHIRE-crYOLO is a fast and accurate fully automated particle picker for cryo-EM. Commun Biol 2: 1–13

Wang Z, Fokine A, Guo X, Jiang W, Rossmann MG, Kuhn RJ, Luo Z-H & Klose T (2023a) Structure of Vibrio Phage XM1, a Simple Contractile DNA Injection Machine. Viruses 15: 1673

Wang Z, Fokine A, Guo X, Jiang W, Rossmann MG, Kuhn RJ, Luo Z-H & Klose T (2023b) Structure of Vibrio Phage XM1, a Simple Contractile DNA Injection Machine. Viruses 15: 1673

Waterhouse AM, Procter JB, Martin DMA, Clamp M & Barton GJ (2009) Jalview Version 2—a multiple sequence alignment editor and analysis workbench. Bioinformatics 25: 1189–1191

Watts NR & Coombs DH (1990) Structure of the bacteriophage T4 baseplate as determined by chemical cross-linking. J Virol 64: 143–154

Weiss GL, Eisenstein F, Kieninger A-K, Xu J, Minas HA, Gerber M, Feldmüller M, Maldener I, Forchhammer K & Pilhofer M (2022) Structure of a thylakoid-anchored contractile injection system in multicellular cyanobacteria. Nat Microbiol 7: 386–396

Wenzel S, Shneider MM, Leiman PG, Kuhn A & Kiefer D (2020) The Central Spike Complex of Bacteriophage T4 Contacts PpiD in the Periplasm of Escherichia coli. Viruses 12: 1135

Williams CJ, Headd JJ, Moriarty NW, Prisant MG, Videau LL, Deis LN, Verma V, Keedy DA, Hintze BJ, Chen VB, et al (2018) MolProbity: More and better reference data for improved all-atom structure validation. Protein Science 27: 293–315

Willis C, Wang CK, Osman A, Simon A, Pickering D, Mulvenna J, Riboldi-Tunicliffe A, Jones MK, Loukas A & Hofmann A (2011) Insights into the Membrane Interactions of the Saposin-Like Proteins Na-SLP-1 and Ac-SLP-1 from Human and Dog Hookworm. PLOS ONE 6: e25369

Wu X, Zhao Y, Sun L, Jiang M, Wang Q, Wang Q, Yang W & Wu Y (2019) Crystal structure of CagV, the *Helicobacter pylori* homologue of the T4 SS protein VirB8. FEBS J 286: 4294–4309

Xia G, Corrigan RM, Winstel V, Goerke C, Gründling A & Peschel A (2011) Wall Teichoic Acid-Dependent Adsorption of Staphylococcal Siphovirus and Myovirus. J Bacteriol 193: 4006–4009

Xu J, Ericson CF, Lien Y-W, Rutaganira FUN, Eisenstein F, Feldmüller M, King N & Pilhofer M (2022) Identification and structure of an extracellular contractile injection system from the marine bacterium Algoriphagus machipongonensis. Nat Microbiol 7: 397–410

Yang F, Jiang Y-L, Zhang J-T, Zhu J, Du K, Yu R-C, Wei Z-L, Kong W-W, Cui N, Li W-F, et al (2023a) Fine structure and assembly pattern of a minimal myophage Pam3. Proc Natl Acad Sci USA 120: e2213727120

Yang F, Wang L, Zhou J, Xiao H & Liu H (2023b) In Situ Structures of the Ultra-Long Extended and Contracted Tail of Myoviridae Phage P1. Viruses 15: 1267

Yu R-C, Yang F, Zhang H-Y, Hou P, Du K, Zhu J, Cui N, Xu X, Chen Y, Li Q, et al (2024) Structure of the intact tail machine of Anabaena myophage A-1(L). Nat Commun 15: 2654

Zhang K (2016) Gctf: Real-time CTF determination and correction. Journal of Structural Biology 193: 1–12

Zheng SQ, Palovcak E, Armache J-P, Verba KA, Cheng Y & Agard DA (2017) MotionCor2: anisotropic correction of beam-induced motion for improved cryo-electron microscopy. Nat Methods 14: 331–332

Zinke M, Schröder GF & Lange A (2022) Major tail proteins of bacteriophages of the order Caudovirales. Journal of Biological Chemistry 298: 101472

Zivanov J, Nakane T & Scheres SHW (2020) Estimation of high-order aberrations and anisotropic magnification from cryo-EM data sets in RELION-3.1. IUCrJ 7: 253–267

