## Supplementary Information for "Conformational changes of baseplate regulating tail contraction of *Staphylococcus* phage 812"

Short title: Tail contraction of *S. aureus* phage 812

Ján Bíňovský[1,2]*, Marta Šiborová[1]*, Maryna Zlatohurska[1], Jiří Nováček[1], Pavol Bárdy[3], Roman Baška[1], Karel Škubník[1], Tibor Botka[3], Martin Benešík[3], Roman Pantůček[3], Konstantinos Tripsianes[1], Pavel Plevka[1#]

[1] *Central European Institute of Technology, Kamenice 753/5, 625 00 Brno, Czech Republic*
[2] *National Centre for Biomolecular Research, Faculty of Science, Masaryk University, 625 00, Brno, Czech Republic*
[3] *Faculty of Science, Masaryk University, Kamenice 753/5, 625 00 Brno, Czech Republic*

** These authors contributed equally to this work*

#### Table of Contents

**Appendix Figures** **2**
**Appendix Tables** **25**
**References for Appendix Table S3** **36**

### Appendix Figures

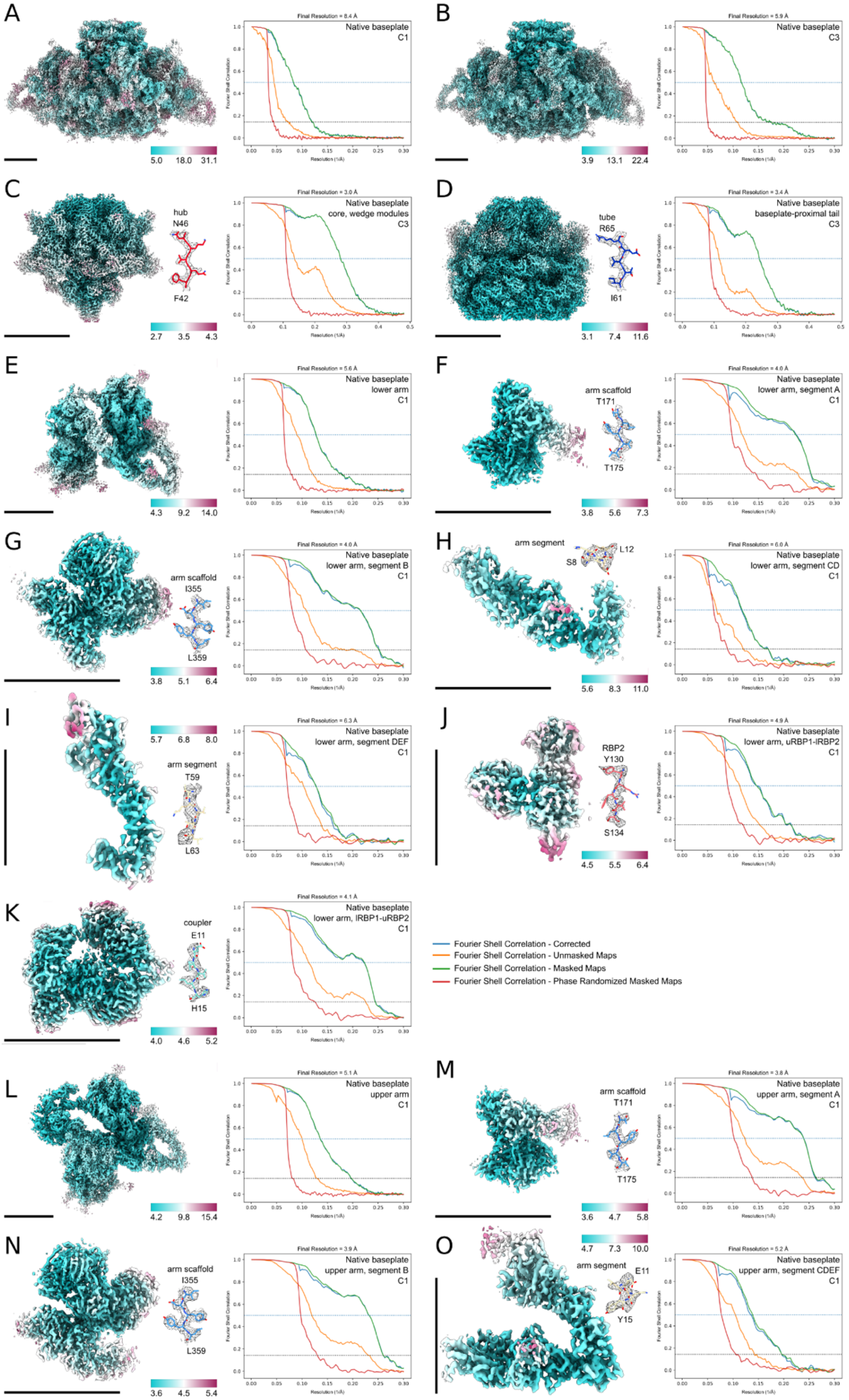

**Appendix Figure S1. Analyses of cryo-EM reconstructions of the baseplate from the phage with an extended tail. (A-O)** Each panel shows a cryo-EM reconstruction colored according to the local resolution in Å, a representative fit of an atomic model shown as sticks to the corresponding cryo-EM density map shown as gray mesh, and a comparison of Fourier shell correlation (FSC) curves color-coded according to the legend. The FSC threshold levels are indicated by a blue dotted line (threshold 0.5) and a black dotted line (threshold 0.143). Panels show reconstructions of the whole baseplate in C1 (A) and C3 (B) symmetry, core and wedge modules (C), baseplate-proximal tail (D), lower baseplate arm (E) and parts of the lower arm (F-K), upper baseplate arm (L), and parts of the upper arm (M-O). uRBP1-lRBP2 = RBP1 from upper arm and RBP2 from lower arm; lRBP1-uRBP2 = RBP1 from lower arm and RBP2 from upper arm.

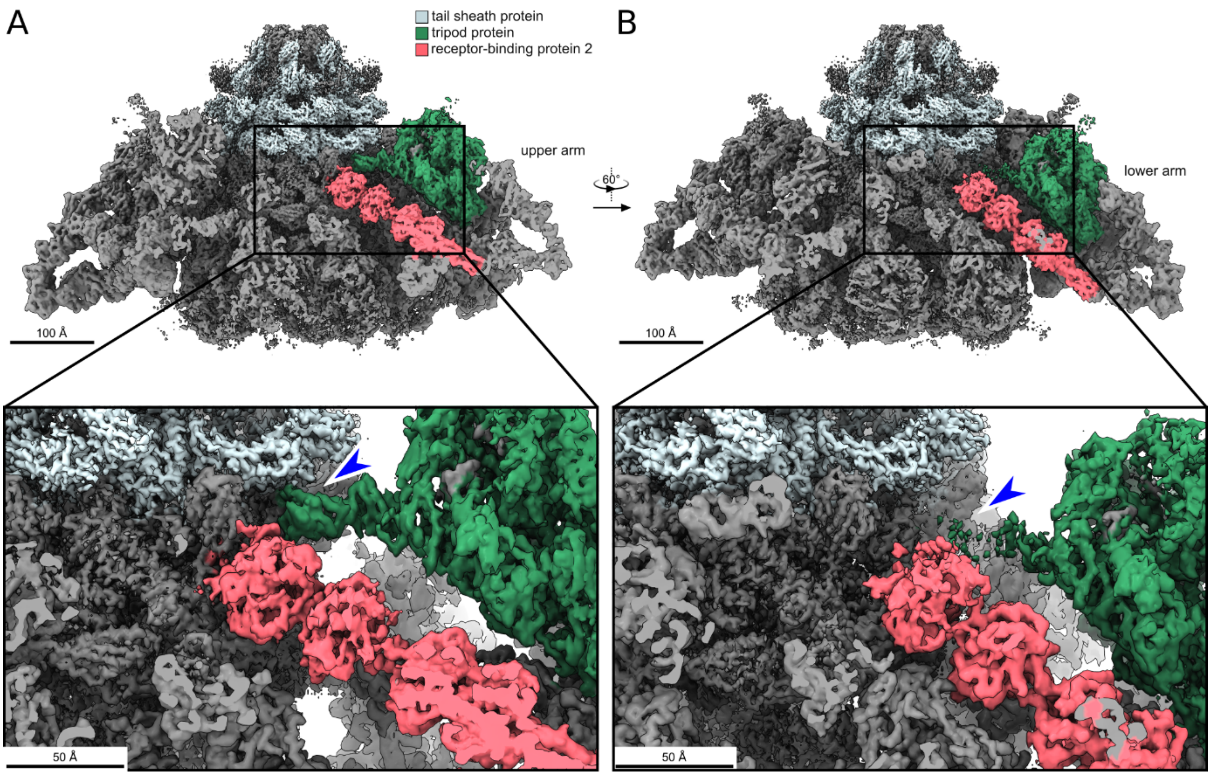

**Appendix Figure S2. Comparison of interactions of tripod proteins and RBP2 attached to upper and lower baseplate arms with baseplate core and tail sheath. (A, B)** Composite cryo-EM map of the baseplate of phage 812 with extended tail. All proteins are colored gray except the tail sheath and selected trimer of tripod proteins and trimer of receptor-binding protein 2. The front part of the baseplate density has been removed to display the interactions of the tripods and receptor-binding proteins from the upper (A) and lower (B) baseplate arms with the baseplate core and tail sheath. Blue arrowheads point to the sites where the interactions of the upper (A) and lower (B) baseplate arms with the tail sheath differ.

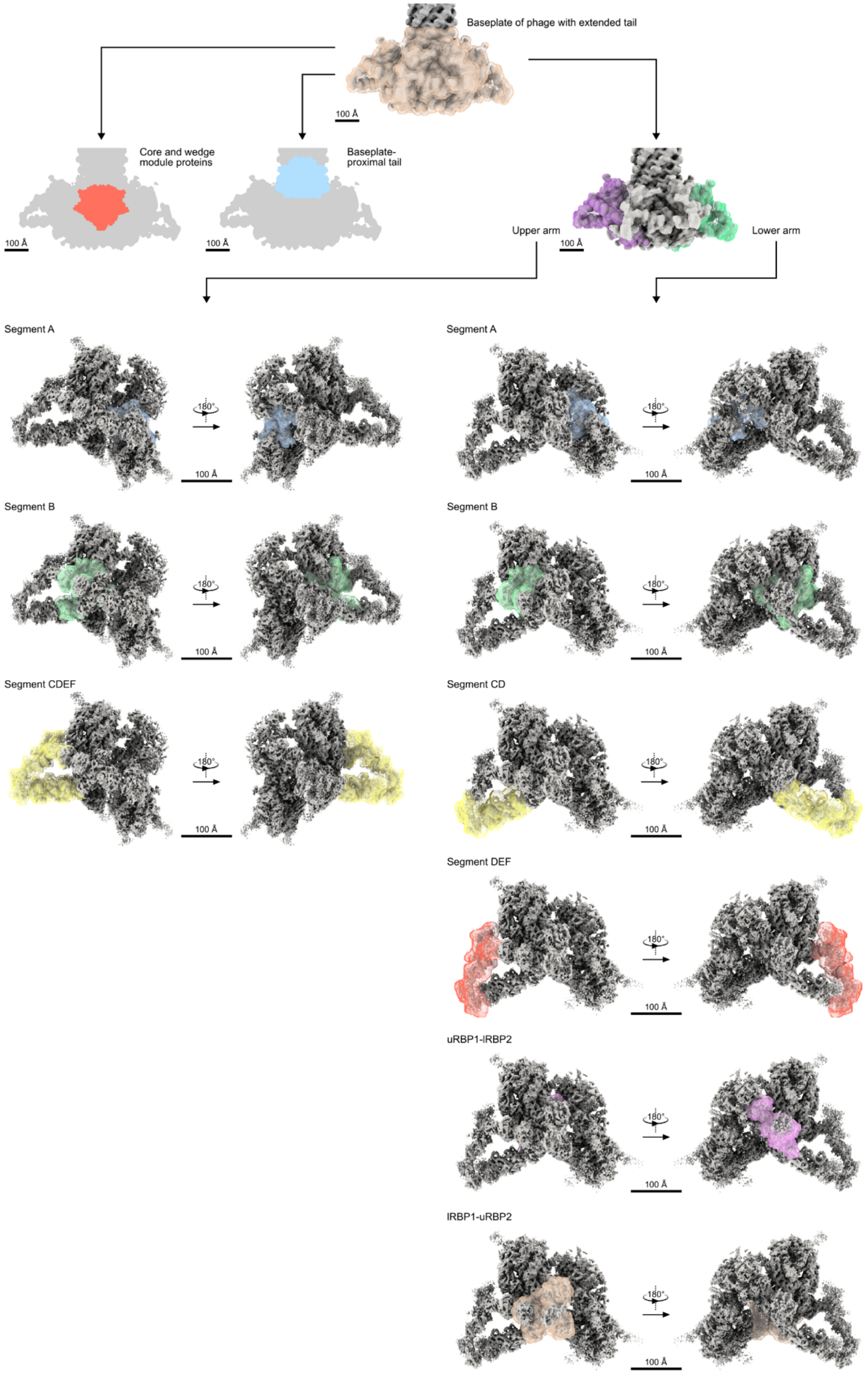

**Appendix Figure S3. Simplified scheme of cryo-EM reconstructions including the baseplate of the phage with an extended tail.** Cryo-EM maps are colored gray, and masks encompassing indicated reconstructions are shown in multiple colors. Maps and masks indicating the reconstruction of the core and wedge module proteins, and reconstruction of the baseplate-proximal tail are shown as flat projections. uRBP1-lRBP2 = RBP1 from upper arm and RBP2 from lower arm; lRBP1-uRBP2 = RBP1 from lower arm and RBP2 from upper arm.

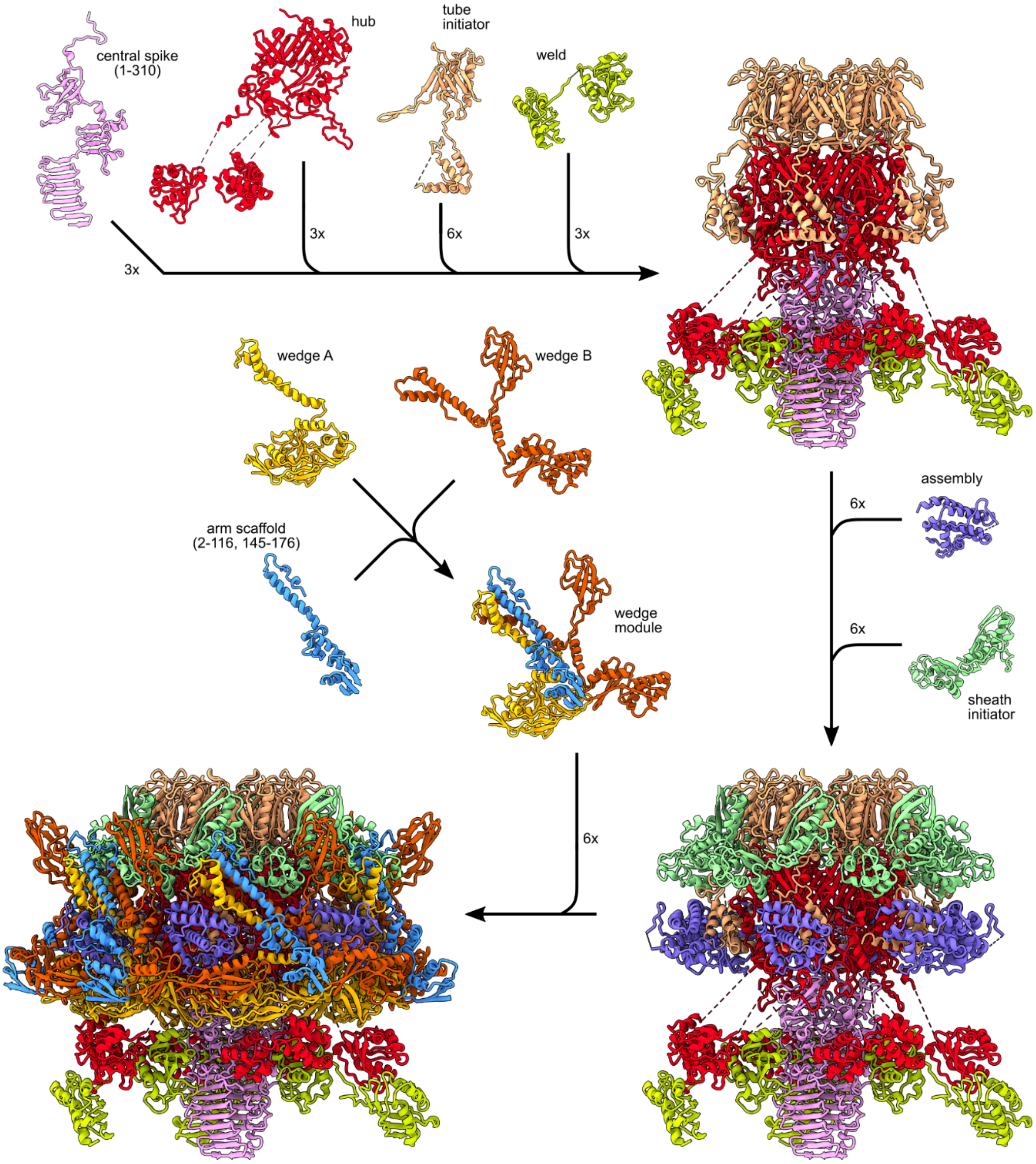

**Appendix Figure S4. Components of phage 812 baseplate core and wedge modules.** The baseplate core and wedge module proteins are shown in cartoon representation. Protein names and the numbers of subunits of each protein forming the baseplate core and wedge modules are indicated.

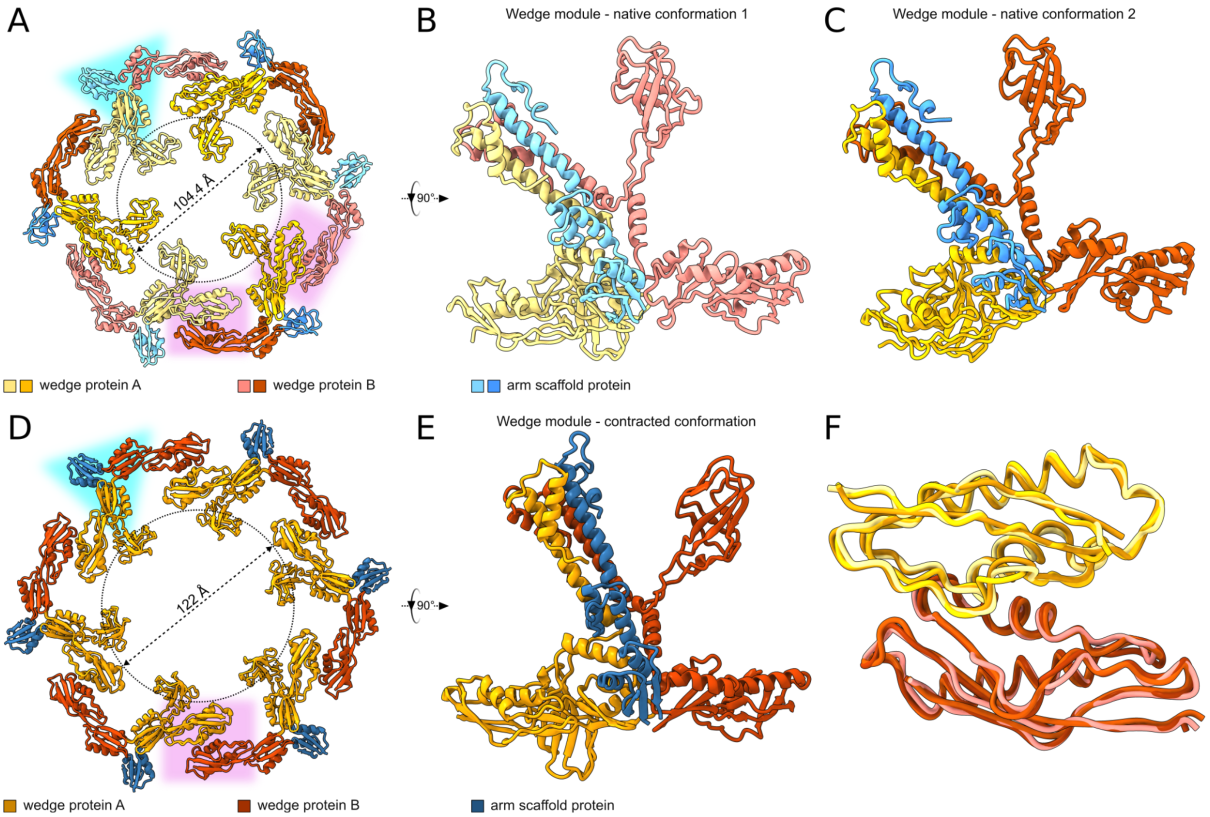

**Appendix Figure S5. Expansion of iris formed by wedge modules upon phage 812 tail contraction. (A-C)** Cartoon representation of wedge modules from the baseplate of phage 812 with extended tail colored according to the legend at the bottom. **(A)** Six wedge modules forming the iris are viewed along the tail axis towards the phage head. Three wedge modules that enable the attachment of lower arms are shown in lighter colors, and the three wedge modules that enable the attachment of upper arms are shown in darker colors. The hetero-trimer interface formed by trifurcation domains is highlighted in cyan. Two conformationally different dimer interactions of dimerization domains are highlighted with magenta backgrounds. **(B, C)** A single wedge module in conformation 1 (B) and 2 (C). **(D-F)** Wedge modules from the contracted baseplate of phage 812 are shown in cartoon representation and colored according to the legend at the bottom. **(D)** Six wedge modules forming the iris structure are viewed along the tail axis towards the phage head. The hetero-trimer interface formed by trifurcation domains is highlighted in cyan. The interfaces of dimerization domains are highlighted in magenta. A dotted circle indicates the inner diameter of the wedge iris. **(E)** A single wedge module from the contracted baseplate. **(F)** Superimposition of three pairs of dimerization domains, which are highlighted in magenta in panels A and D.

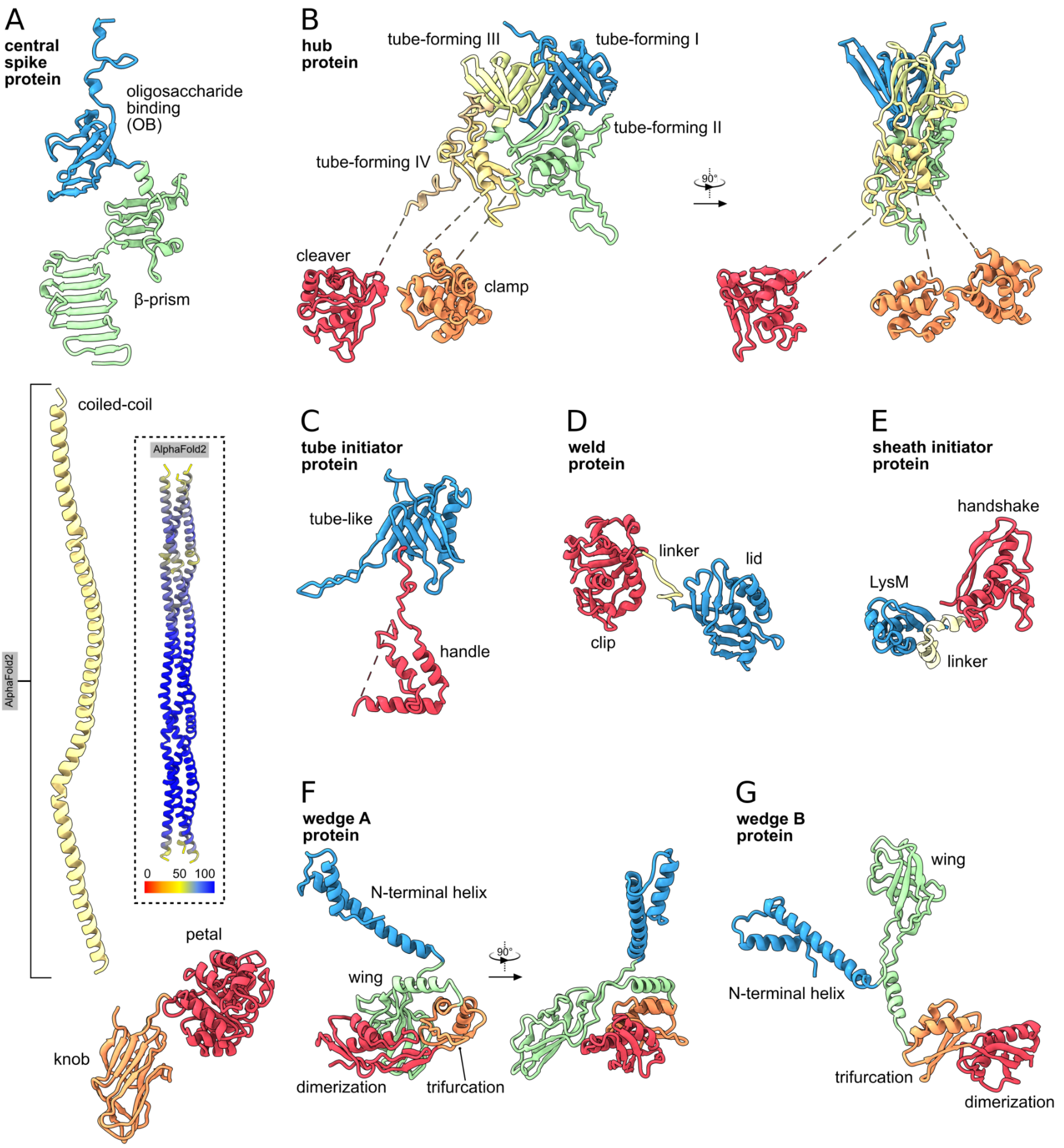

**Appendix Figure S6. Domain organization of proteins forming baseplate core and wedge modules of phage 812. (A-G)** The proteins are shown in cartoon representation and colored according to domains and functional elements. The names of proteins are in bold. The coiled-coil domains of the central spike proteins (panel A, dashed rectangle) are colored according to the AlphaFold2 pLDDT score.

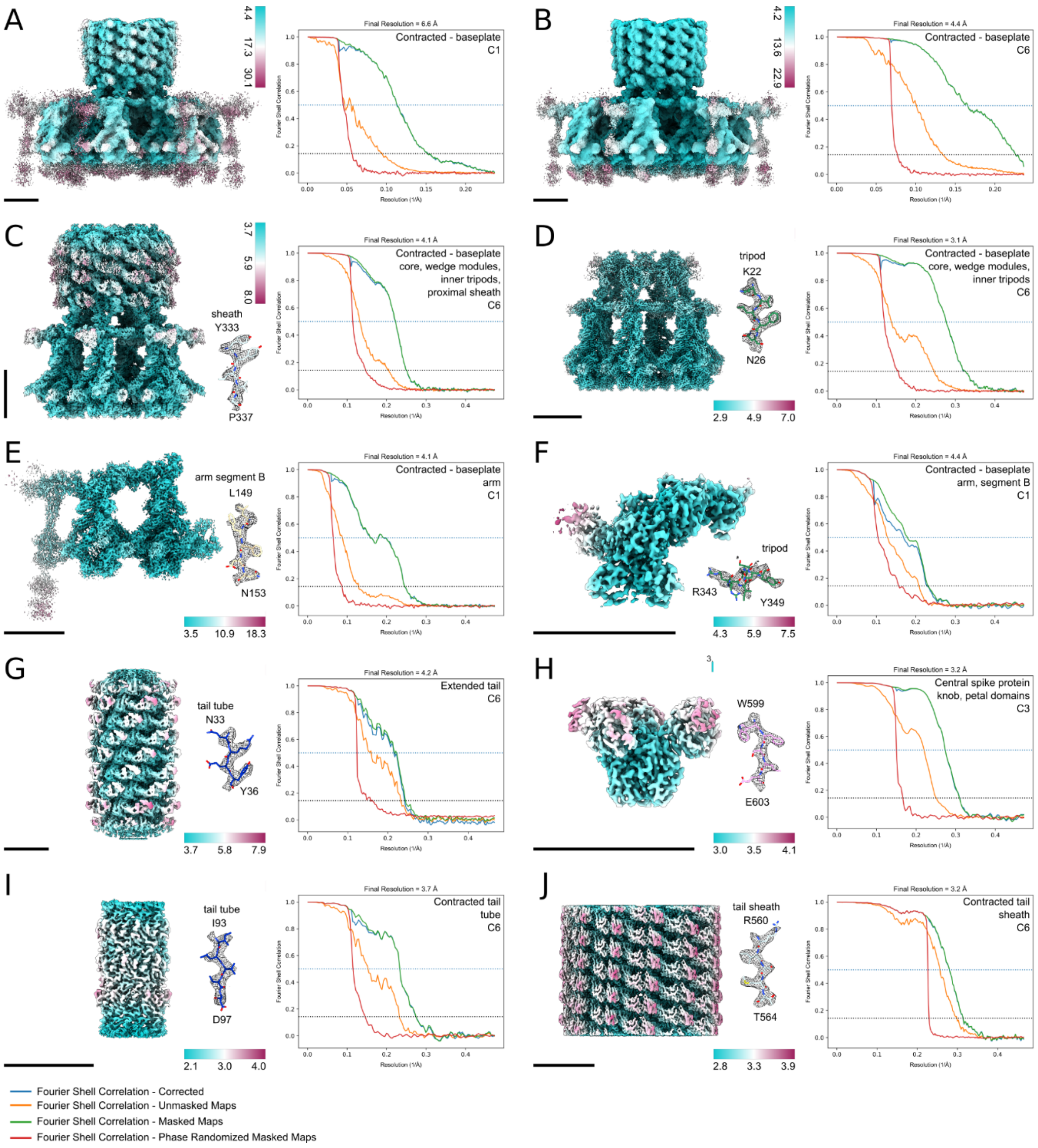

**Appendix Figure S7. Analyses of cryo-EM reconstructions of the baseplate from the phage with a contracted tail, reconstructions of the extended and contracted tail, and reconstruction of the central spike protein. (A-J)** Each panel shows a cryo-EM reconstruction colored according to the local resolution in Å, a representative fit of an atomic model shown as sticks to the corresponding cryo-EM density map shown as gray mesh, and a comparison of Fourier shell correlation (FSC) curves color-coded according to the legend. The FSC threshold levels are indicated by a blue dotted line (threshold 0.5) and a black dotted line (threshold 0.143). Panels show reconstructions of the whole baseplate in C1 (A) and C6 (B) symmetry, core and wedge modules, inner tripods and baseplate-proximal tail sheath (C), core and wedge modules and inner tripods (D), baseplate arm (E), part of the baseplate arm (F), extended tail (G), knob and petal domains of the central spike protein (H), tail tube from the contracted tail (I) and tail sheath from the contracted tail (J).

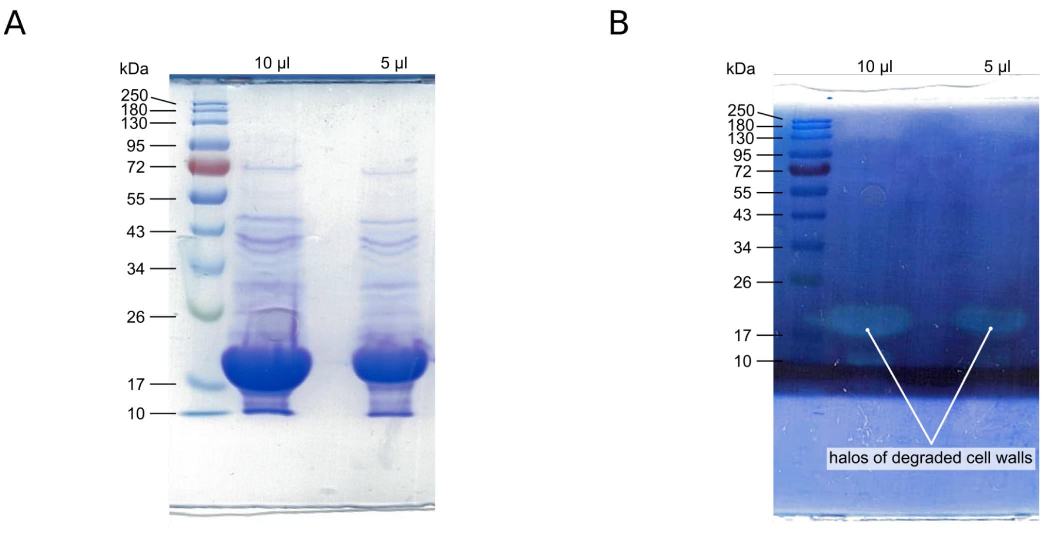

**Appendix Figure S8. SDS-PAGE and zymogram analysis of cleaver domain of hub protein.** Scan of an SDS-PAGE gel **(A)** and zymogram **(B)** of recombinantly expressed cleaver domain of the hub protein after Histrap purification. As indicated, ten and five µl of the purified protein sample concentrated to 10 mg/µl were loaded. Halos on the zymogram resulting from cell wall degradation activity of the cleaver domain of the hub protein are indicated.

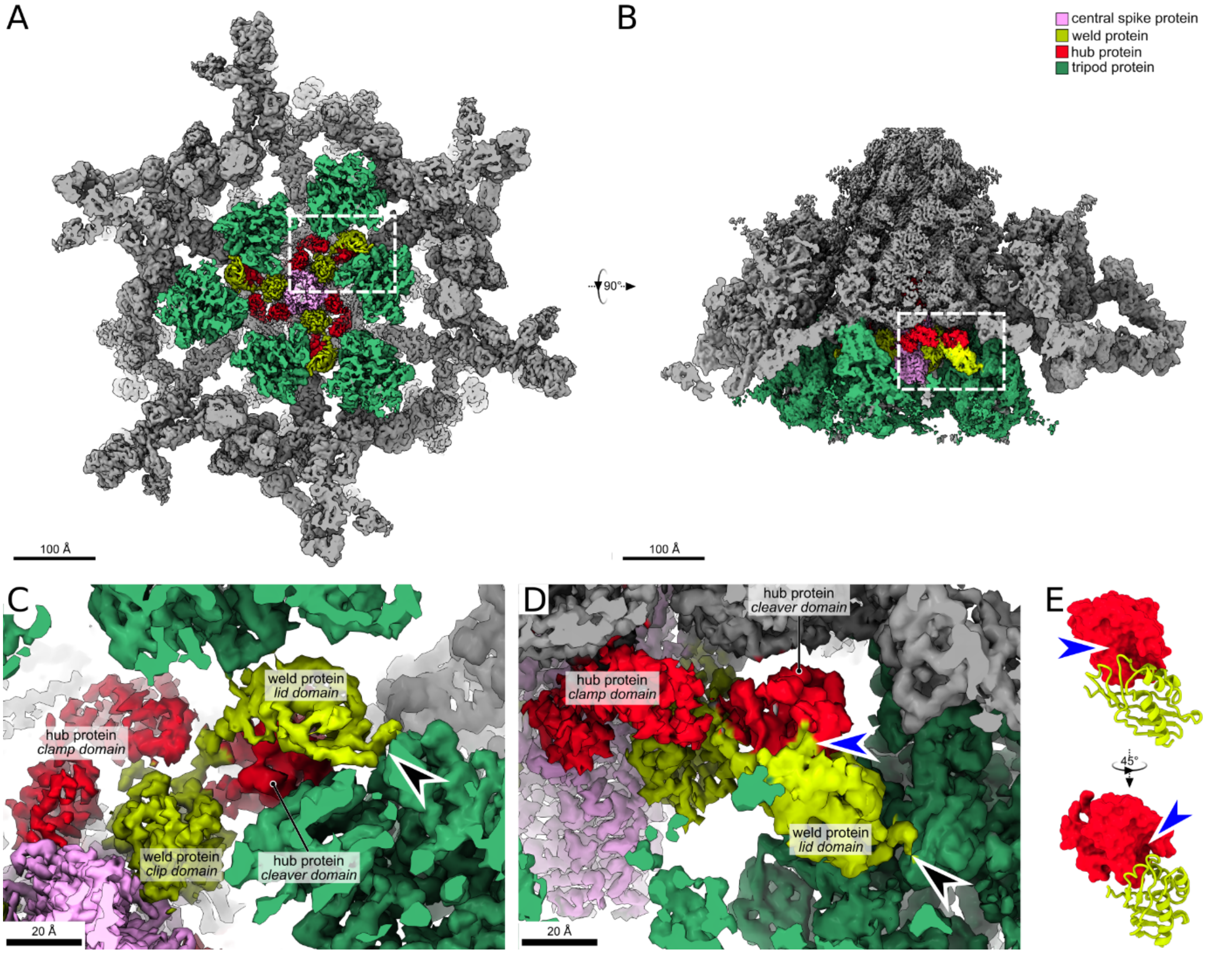

**Appendix Figure S9. Interactions of weld proteins with baseplate. (A-D)** Composite cryo-EM map of the phage 812 baseplate is viewed along the tail axis towards the phage head (A, C) and perpendicular to the tail axis (B, D). Baseplate proteins are shown in gray except for the hub protein, the weld protein, the central spike protein, and the inner tripod proteins, which are colored according to the legend in (B). The front density in panels A and B has been clipped away to show the interactions of the weld protein with the rest of the baseplate. Rectangles in A and B indicate close views, as shown in (C) and (D), respectively. Black arrowheads in C and D point to the interaction between the lid domain of the weld protein and the inner tripod protein. The blue arrowhead in D points to the site where the lid domain of the weld protein covers the active site of the cleaver domain of the hub protein. **(E)** Interaction between the cleaver domain of the hub protein shown as molecular surface and lid domain of the weld protein shown as cartoon. The blue arrowhead points to the active site of the cleaver domain.

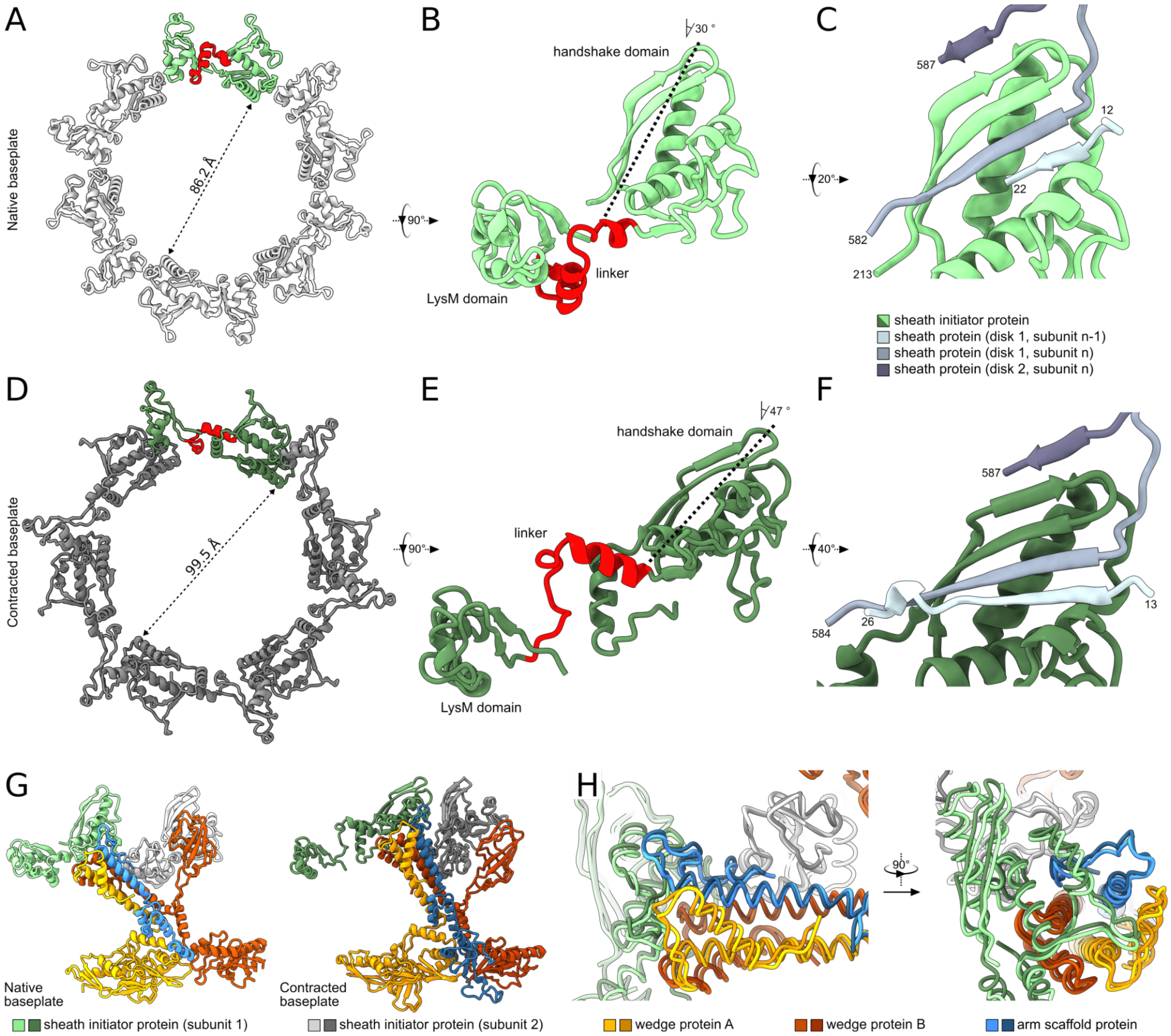

**Appendix Figure S10. Conformational changes and β-sheet augmentation of handshake domain of tail sheath initiator protein. (A-C)** Cartoon representation of the tail sheath initiator protein from the baseplate of phage 812 with extended tail. **(A)** Five subunits from the hexamer of the tail sheath initiator proteins are shown in light gray, and one subunit in light green, with the linker between the LysM and handshake domains in red. **(B)** The structure of the tail sheath initiator protein with the linker between the LysM and handshake domains is shown in red. The angle between the tail axis and the axis of the handshake domain is indicated. **(C)** Augmentation of the β-sheet of handshake domain of the tail sheath initiator protein in the baseplate of phage 812 with extended tail with β-strands formed by an N-terminus and C-termini from three tail sheath proteins colored in three shades of gray. **(D-F)** Structure of tail sheath initiator proteins from the baseplate of phage 812 with a contracted tail. **(D)** The hexamer of the tail sheath initiator proteins with five subunits is shown in dark gray, and one subunit is shown in dark green, with the linker between the LysM and handshake domains in red. **(E)** Structure of the tail sheath initiator protein. **(F)** Augmentation of the β-sheet of handshake domain of tail sheath initiator protein in phage 812 with a contracted tail with β-strands formed by an N-terminus and C-termini from three tail sheath proteins colored in three shades of gray. **(G)** Interactions of tail sheath initiator proteins and a wedge module in baseplates from phage particles with extended and contracted tails. **(H)** Differences between the interactions of tail sheath initiator proteins and wedge modules in baseplates from phage particles with extended and contracted tails.

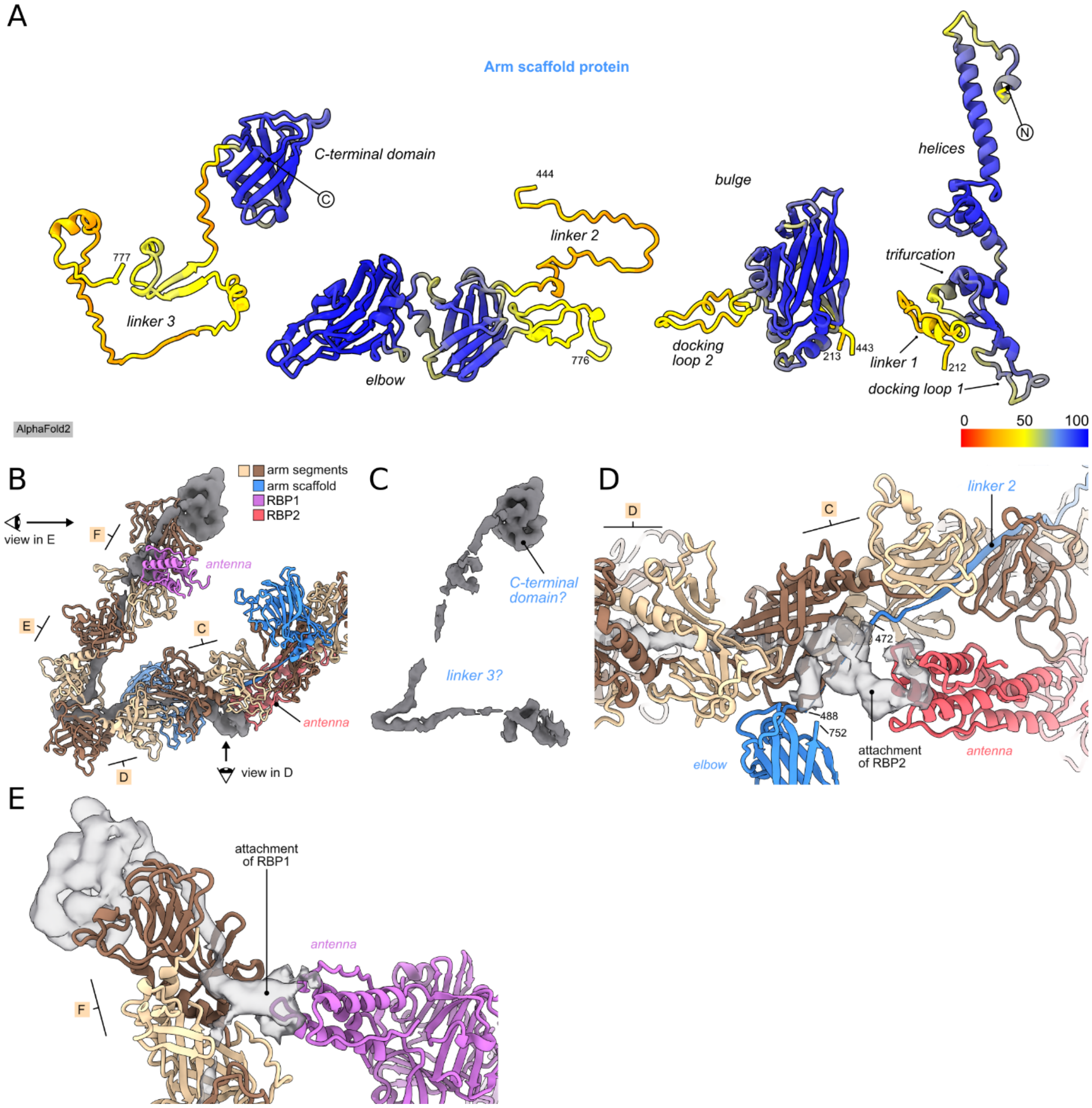

**Appendix Figure S11. Arm scaffold protein. (A)** AlphaFold2-predicted structure of the arm scaffold protein is shown in cartoon representation and colored according to pLDDT score. The structure was divided into several parts for better visualization. Domain names and chain termini are indicated. **(B)** Peripheral part of the lower baseplate arm. Protein structures are shown in cartoon and colored according to the protein types indicated in the top right corner of the panel B. Cryo-EM density, for which no protein structure was assigned, is shown as opaque dark gray surface. Arm segments C-F and antennae domains of RBP1 and RBP2 are indicated. Views of panels D and E are indicated. **(C)** Cryo-EM density from the panel B shown alone. Putative assignment of the density to parts of the arm scaffold protein is indicated. **(D, E)** Close views as indicated in the panel B.

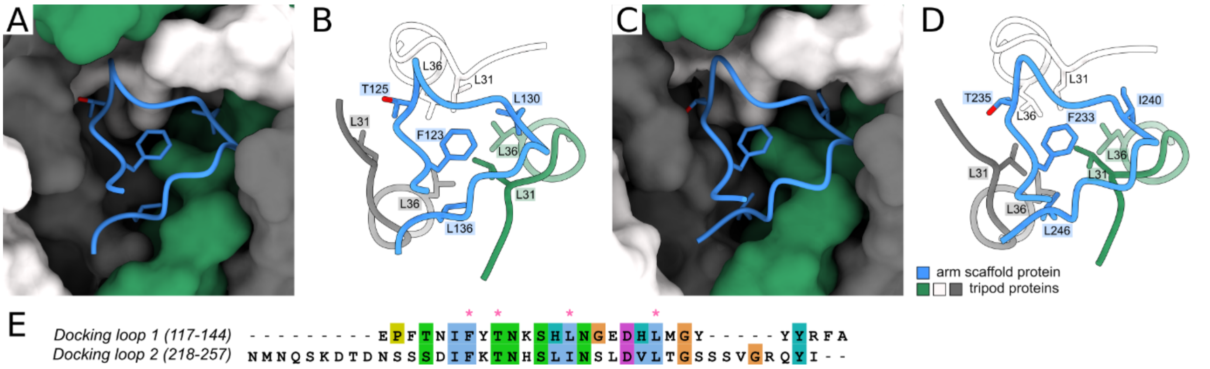

**Appendix Figure S12. Mechanism of binding of tripod trimers to arm scaffold protein. (A, B)** Interface between trimer of the inner tripod proteins and the docking loop 1 from the arm scaffold protein from baseplate of phage 812 with an extended tail. Tripod proteins are shown in molecular surface (A) and cartoon (B), arm scaffold protein is shown in cartoon. Side chains of residues contributing to the hydrophobic interaction are shown as sticks. Structures are colored according to the legend in the panel D. **(C, D)** Interface between trimer of the outer tripod proteins and the docking loop 2 from the arm scaffold protein from baseplate of phage 812 with an extended tail. Style and colors in panels C and D correspond to those in panels A and B, respectively. **(E)** Sequence alignment of docking loops 1 and 2 of the arm scaffold protein. Pink asterisks indicate residues of docking loop 1 participating in the interaction with the tripod proteins, as shown in panel B and D.

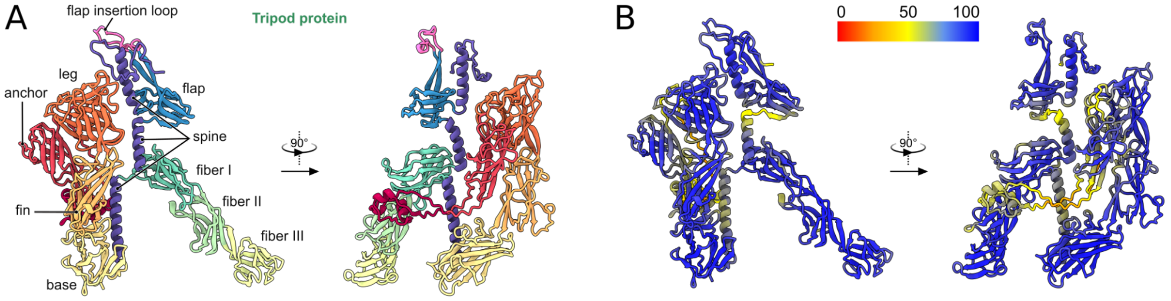

**Appendix Figure S13. Prediction of the tripod protein structure. (A, B)** AlphaFold2-predicted structure of the tripod protein is shown in a cartoon representation and colored according to domains (A) and the pLDDT score (B). In panel A, domain names are indicated.

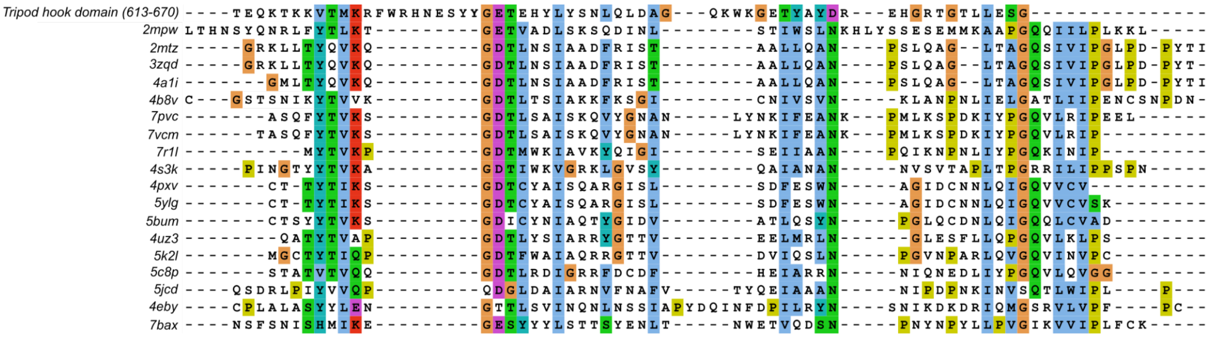

**Appendix Figure S14. The hook domain of the tripod protein shares conserved residues with proteins from the LysM family.** Primary sequence alignment of residues 613-670 forming the hook domain, with LysM protein family members (PDB codes are listed).

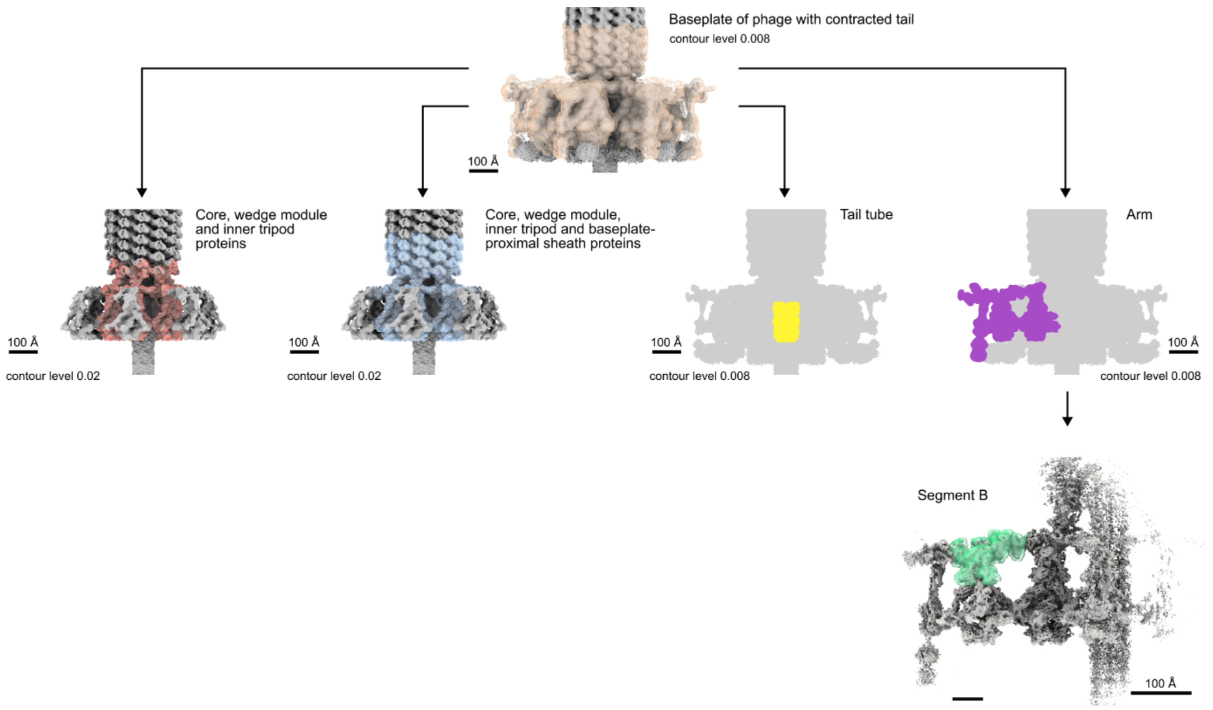

**Appendix Figure S15. Simplified scheme of cryo-EM reconstructions including the baseplate of the phage with contracted tail.** Cryo-EM maps are colored gray, and masks encompassing indicated reconstructions are shown in multiple colors. Maps and masks indicating the reconstruction of the tail tube and the reconstruction of the baseplate arm are shown as flat projections.

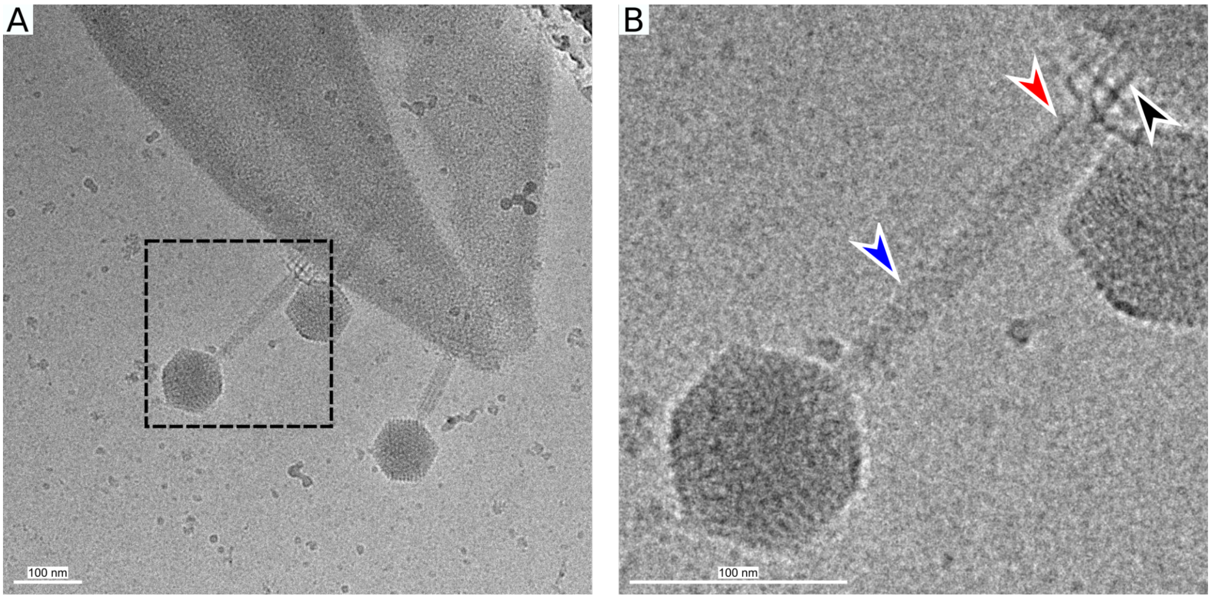

**Appendix Figure S16. Particle of phage 812 captured in an intermediate state of tail contraction. (A)** Cryo-electron micrograph of phage 812 attached to a cell wall of *S. aureus*. The black rectangle indicates the detail in B. **(B)** phage 812 particle with a semi-contracted tail sheath. The blue arrowhead points to the sheath in the extended conformation, whereas the red arrowhead points to the sheath in the contracted conformation. The black arrowhead points to the densities at the bottom of the contracted baseplate, presumably belonging to the anchor and hook domains of tripod proteins.

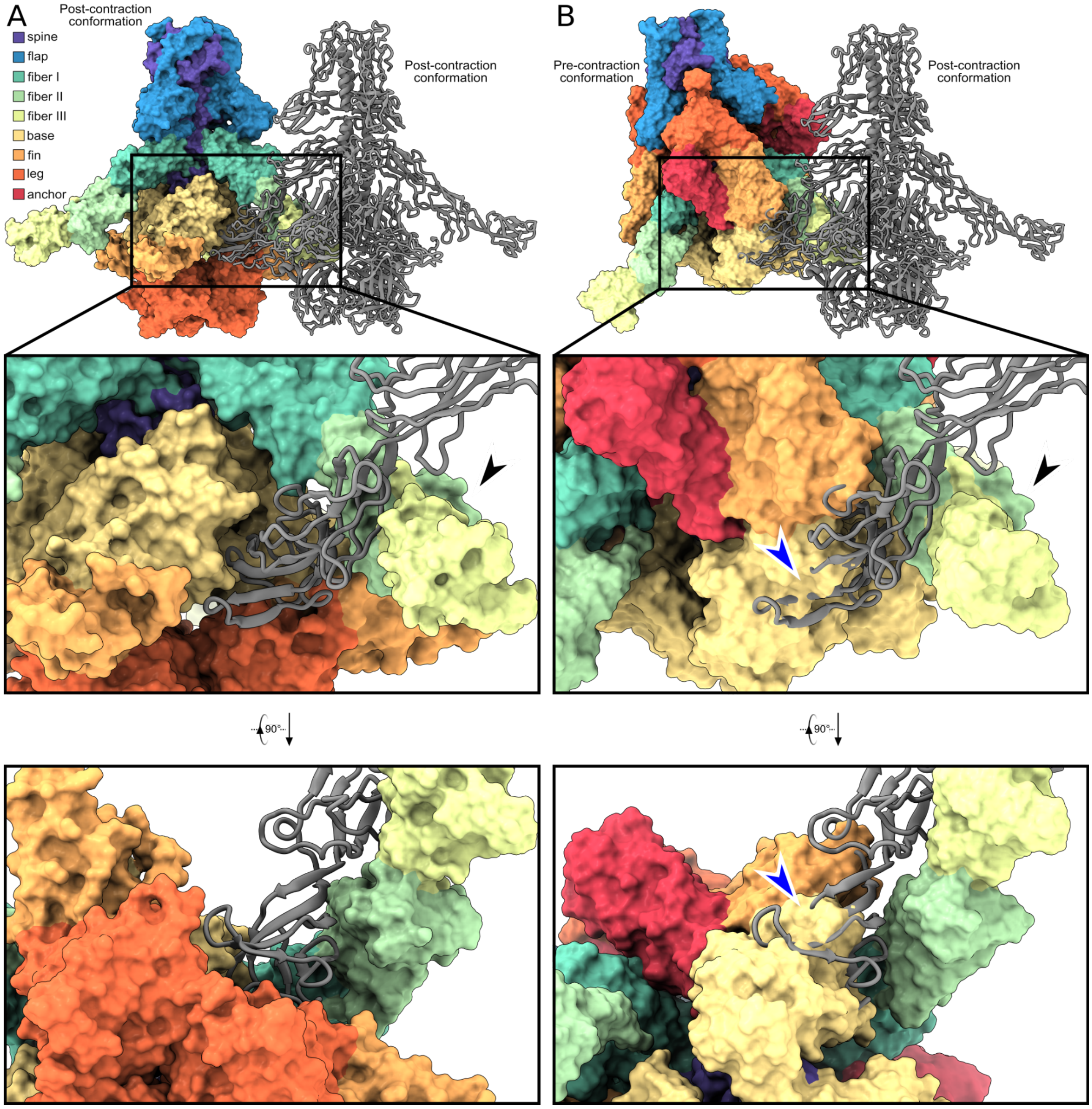

**Appendix Figure S17. Interactions between tripod complexes in the baseplate of phage 812 particle with contracted tail. (A)** Interactions between two neighboring inner tripod complexes. One tripod complex is shown as a molecular surface colored according to domain, and the other is shown in gray as a cartoon representation. The black rectangle indicates the magnified view shown in the insets below. The upper inset shows the interactions between fiber domains viewed perpendicular to the tail axis, and the lower inset shows the view along the tail axis toward the phage head. The black arrowhead points to the fiber domains of a tripod protein subunit used to superimpose the tripod complex in the pre-contraction state, as shown in B. **(B)** The inner tripod complex in the pre-contraction conformation is shown as a molecular surface colored according to domain, and the tripod complex in the post-contraction state is shown in gray cartoon representation. The pre-contraction tripod complex was superimposed onto the post-contraction tripod complex using the fiber domains indicated by the black arrowhead. The black rectangle indicates the close view shown in the insets below. The upper inset shows the view perpendicular to the tail axis, and the lower inset shows the view along the tail axis and towards the phage head. Blue arrowheads point to the site where the fiber III domains of the post-contraction tripod complex clash with the base domain of the pre-contraction tripod complex.

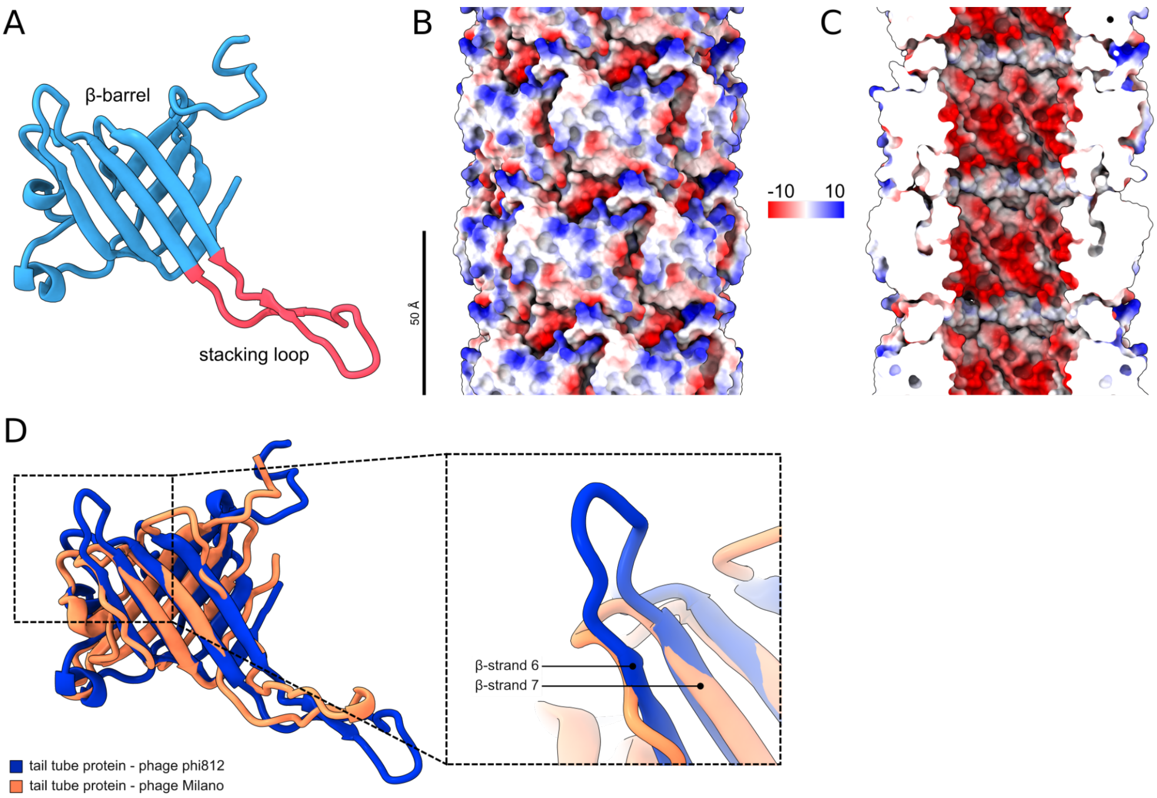

**Appendix Figure S18. Tail tube proteins. (A)** Cartoon representation of the phage 812 tail tube protein colored according to domains. **(B, C)** Molecular surface representation of the phage 812 tail tube viewed as a whole (B) and clipped (C), colored according to Coulombic electrostatic potential in the range from -10 to 10 kcal/(mol·*e*-) at 298 K. **(D)** Cartoon representation of the tail tube proteins from phage phage 812 and phage Milano superposed by the β-strands 6 and 7. The inset shows a close view of the loops between β-strands 6 and 7.

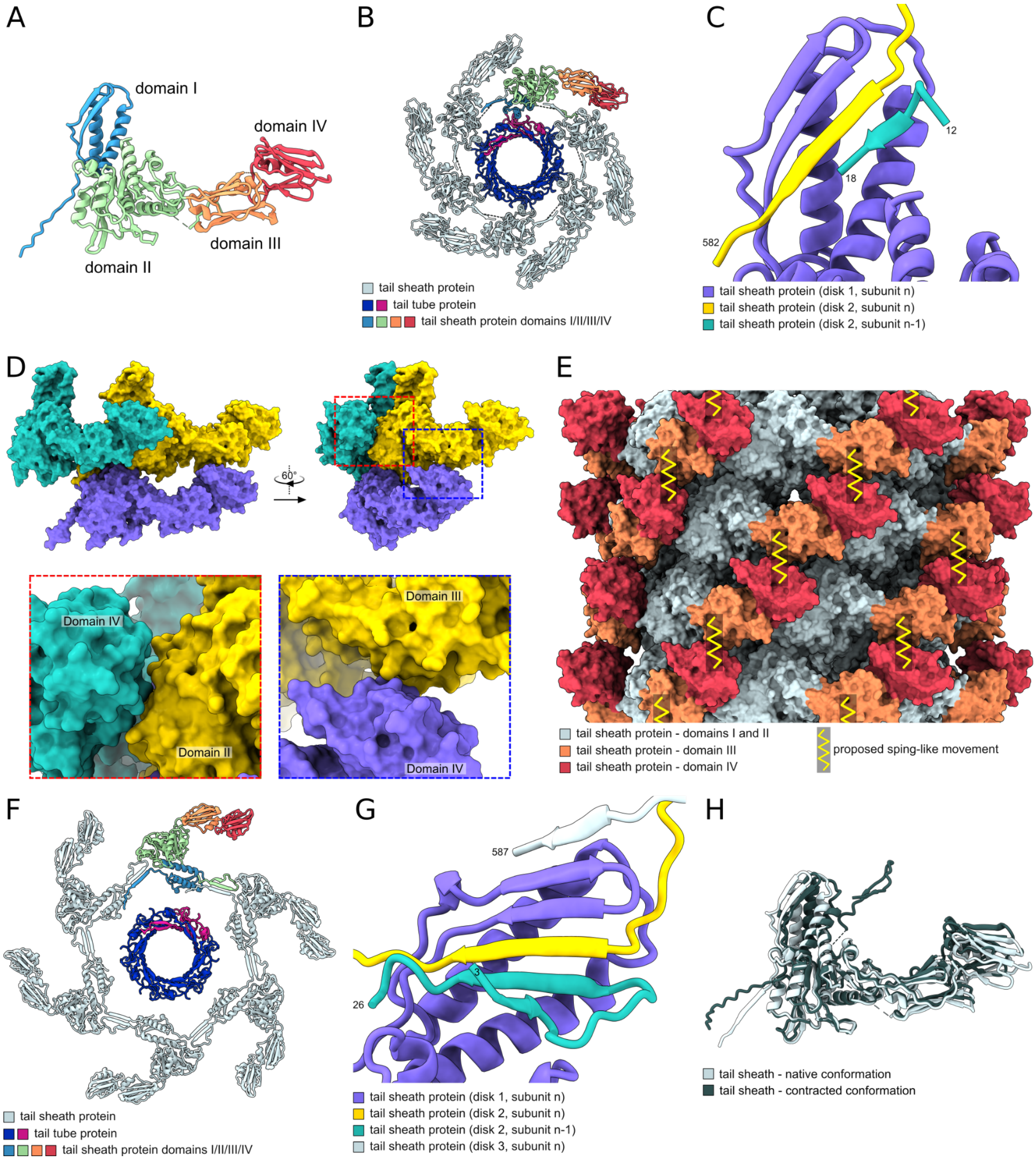

**Appendix Figure S19. Tail sheath proteins. (A)** Cartoon representation of the phage 812 tail sheath protein colored according to domains. **(B)** Cartoon representation of a disc of tail tube and tail sheath proteins from the extended phage 812 tail viewed along the tail axis towards the phage head. Protein subunits are colored according to the legend. **(C)** β-sheet augmentation of the handshake domain of the tail sheath protein in extended conformation with the N-terminus and two C-termini from other tail sheath proteins. The numbers of terminal residues of the displayed structures are indicated. **(D)** Molecular surface representations of three tail sheath proteins in their extended conformation. Red and blue dashed rectangles indicate views of interactions among the subunits shown as insets. **(E)** Molecular surface representation of four tail sheath disks colored according to the legend. The proposed vertical spring-like movement of domains III and IV of the tail sheath protein is indicated. **(F)** Cartoon representation of a disc of the tail tube and tail sheath proteins from the contracted tail viewed along the tail axis towards the phage head. The relative rotation of the tail tube disc against the sheath disc is arbitrary. The proteins are colored according to the legend. **(G)** β-sheet augmentation of the handshake domain of the tail sheath protein from the contracted tail. Terminal residues are indicated. **(H)** Comparison of the tail sheath proteins from extended (light gray) and contracted (black) tails. The subunits were superimposed by aligning their domains II.

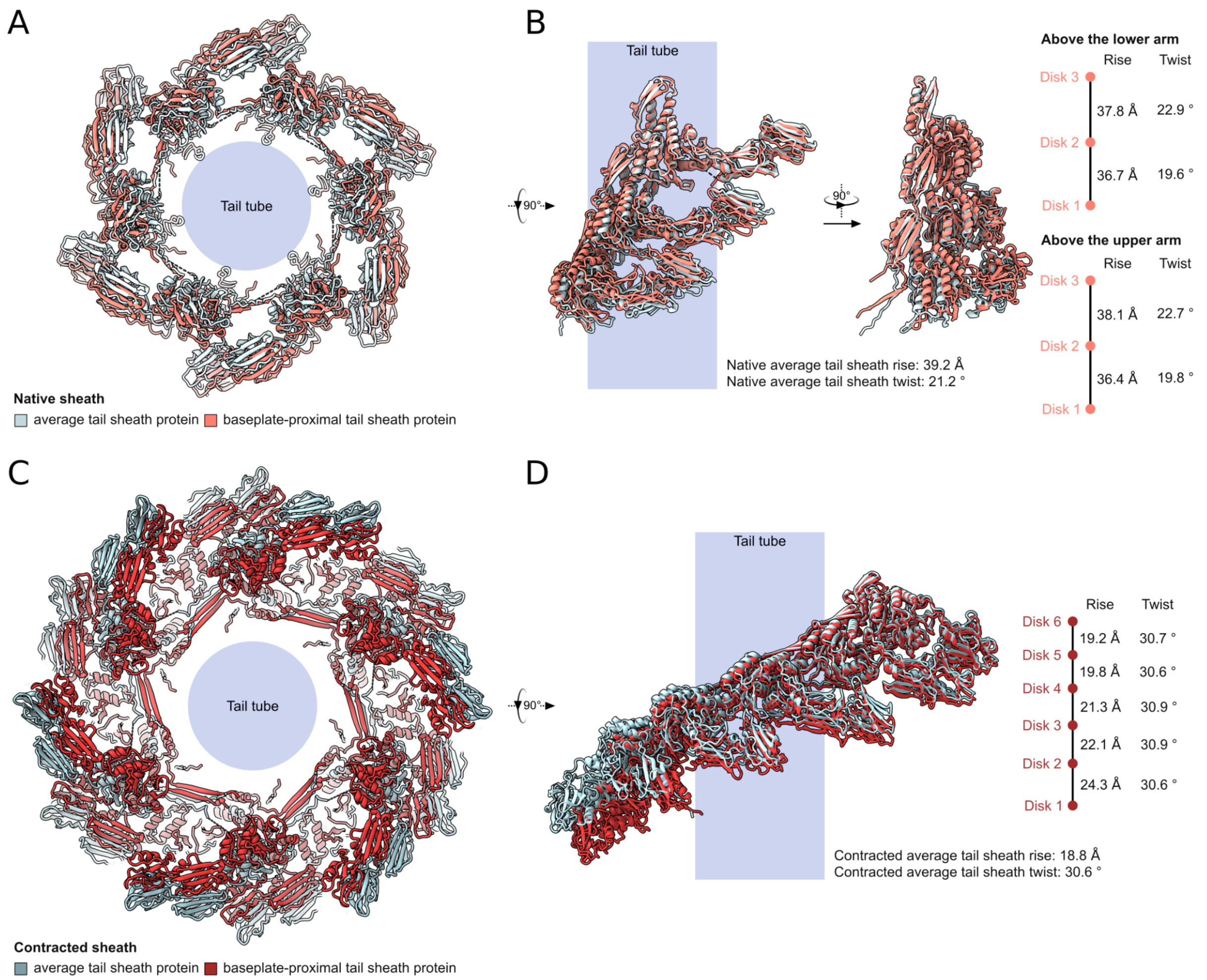

**Appendix Figure S20. Baseplate-proximal discs of tail sheath proteins of phage 812 in extended and contracted states deviate from respective tail sheath structures at the center of the tail. (A)** Comparison of the cartoon representations of three baseplate-proximal tail sheath discs to three tail sheath discs from the central part of the tail. The structures are viewed along the tail axis towards the phage head. The position of the tail tube is indicated. Rise and twist differences were determined by superimposing domains I of tail sheath proteins from the uppermost discs. **(B)** Comparison of distances between tail sheath proteins from three baseplate-proximal tail sheath discs to three tail sheath discs from the central part of the tail. The twist and pitch parameters are indicated. **(C)** Comparison of the cartoon representations of six baseplate-proximal contracted tail sheath discs to six tail sheath discs from the central part of the tail. The structures are viewed along the tail axis towards the phage head. The position of the tail tube is indicated. **(D)** The scene from panel C is viewed perpendicularly to the tail axis. Only one tail sheath protein from six of each average and baseplate-proximal tail sheath disc is shown. The rise and twist difference between tail sheath subunits from successive discs is indicated on the right.

### Appendix Tables

**Appendix Table S1. Cryo-EM data and structure quality indicators – part 1**.

| | **Baseplate of extended tail (core, wedge modules, arms, proximal tail)** | **Baseplate of extended tail (core, wedge modules, arms, proximal tail)** | **Baseplate of extended tail (core, wedge modules, arms, proximal tail) - composite** | **Baseplate of extended tail (core, wedge modules)** | **Baseplate of extended tail (proximal tail)** |
|---|---|---|---|---|---|
| **Sample** | Particles with extended tails | Particles with extended tails | Particles with extended tails | Particles with extended tails | Particles with extended tails |
| **Data collection and processing** | | | | | |
| Detector | K3 | K3 | K3 | K3 | K3 |
| Magnification | 105,000 | 105,000 | 105,000 | 105,000 | 105,000 |
| Voltage [kV] | 300 | 300 | 300 | 300 | 300 |
| Exposure [$e^-$/Å$^2$] | 40.8 | 40.8 | 40.8 | 40.8 | 40.8 |
| Pixel size [Å] | 0.8336 | 0.8336 | 0.8336 | 0.8336 | 0.8336 |
| Symmetry | C1 | C3 | C3 | C3 | C3 |
| Final number of particles | 3,586 | 3,586 | N/A | 8,368 | 4,071 |
| Map resolution [Å] | 8.4 | 5.9 | N/A | 3.0 | 3.4 |
| FSC threshold | 0.143 | 0.143 | N/A | 0.143 | 0.143 |
| **Accession codes** | | | | | |
| EMDB | EMD-55950 | EMD-55951 | EMD-55977 | EMD-55952 | EMD-55953 |
| PDB | N/A | 9TIC | N/A | 9TID | 9TIE |
| **Model composition** | | | | | |
| Atoms (except hydrogens) | N/A | 256,952 | N/A | 32,215 | 30,872 |
| Residues | N/A | 32,543 | N/A | 4,021 | 3,956 |
| B-factors | N/A | 287.6 | N/A | 35.8 | 40.72 |
| FSC (model) at 0.5 (masked) | N/A | 7.9 | N/A | 3.2 | 3.7 |
| **RMSD** | | | | | |
| Bond length [Å] | N/A | 0.012 | N/A | 0.001 | 0.001 |
| Bond angles [°] | N/A | 1.82 | N/A | 0.36 | 0.47 |
| **Validation** | | | | | |
| MolProbity score [percentile] | N/A | 1.96 | N/A | 1.20 | 1.20 |
| ClashScore [percentile] | N/A | 1.97 | N/A | 4.20 | 4.14 |
| Poor rotamers [%] | N/A | 4.25 | N/A | 0.33 | 0.77 |
| C-β outliers [%] | N/A | 0.51 | N/A | N/A | N/A |
| **Ramachandran plot** | | | | | |
| Outliers [%] | N/A | 1.4 | N/A | 0.0 | 0.03 |
| Favored [%] | N/A | 91.1 | N/A | 98.5 | 98.2 |

N/A = not applicable

**Appendix Table S1. Cryo-EM data and structure quality indicators – part 2**.

| | **Lower arm of baseplate of extended tail** | **Lower arm of baseplate of extended tail (segment A)** | **Lower arm of baseplate of extended tail (segment B)** | **Lower arm of baseplate of extended tail (segment C)** | **Lower arm of baseplate of extended tail (segments DEF)** | **Lower arm of baseplate of extended tail (uRBP1-lRBP2)** | **Lower arm of baseplate of extended tail (lRBP1-uRBP2)** |
|---|---|---|---|---|---|---|---|
| **Sample** | Particles with extended tails | Particles with extended tails | Particles with extended tails | Particles with extended tails | Particles with extended tails | Particles with extended tails | Particles with extended tails |
| **Data collection and processing** | | | | | | | |
| Detector | K3 | K3 | K3 | K3 | K3 | K3 | K3 |
| Magnification | 105,000 | 105,000 | 105,000 | 105,000 | 105,000 | 105,000 | 105,000 |
| Voltage [kV] | 300 | 300 | 300 | 300 | 300 | 300 | 300 |
| Exposure [$e^-$/Å$^2$] | 40.8 | 40.8 | 40.8 | 40.8 | 40.8 | 40.8 | 40.8 |
| Pixel size [Å] | 0.8336 | 0.8336 | 0.8336 | 0.8336 | 0.8336 | 0.8336 | 0.8336 |
| Symmetry | C1 | C1 | C1 | C1 | C1 | C1 | C1 |
| Final number of particles | 22,031 | 19,338 | 11,128 | 6,229 | 22,031 | 22,031 | 22,031 |
| Map resolution [Å] | 5.6 | 4.0 | 4.0 | 6.0 | 6.3 | 4.9 | 4.1 |
| FSC threshold | 0.143 | 0.143 | 0.143 | 0.143 | 0.143 | 0.143 | 0.143 |
| **Accession codes** | | | | | | | |
| EMDB | EMD-55954 | EMD-55955 | EMD-55956 | EMD-55957 | EMD-55958 | EMD-55959 | EMD-55960 |
| PDB | 9TIF | 9TIG | 9TIH | 9TII | 9TIJ | 9TIK | 9TIL |
| **Model composition** | | | | | | | |
| Atoms (except hydrogens) | 100,209 | 7,377 | 14,699 | 11,548 | 7,642 | 9,786 | 15,643 |
| Residues | 12,698 | 934 | 1,839 | 1,464 | 971 | 1,260 | 1,959 |
| B-factor | 191.1 | 27.7 | 29.2 | 123.3 | 114.6 | 145.7 | 10.6 |
| FSC (model) at 0.5 (masked) | 6.0 | 4.1 | 4.1 | 7.4 | 6.8 | 5.5 | 4.2 |
| **RMSD** | | | | | | | |
| Bond length [Å] | 0.013 | 0.002 | 0.002 | 0.013 | 0.013 | 0.012 | 0.002 |
| Bond angles [°] | 2.06 | 0.47 | 0.49 | 2.26 | 2.22 | 1.89 | 0.41 |
| **Validation** | | | | | | | |
| MolProbity score [percentile] | 2.15 | 1.24 | 1.23 | 2.53 | 2.42 | 1.24 | 1.25 |
| ClashScore [percentile] | 2.29 | 4.73 | 3.56 | 5.35 | 3.35 | 1.19 | 4.08 |
| Poor rotamers [%] | 5.88 | 0.24 | 0.18 | 7.16 | 8.67 | 1.44 | 0.11 |
| C-β outliers [%] | 0.62 | N/A | N/A | 1.80 | 2.06 | 0.52 | N/A |
| **Ramachandran plot** | | | | | | | |
| Outliers [%] | 1.5 | 0.0 | 0.0 | 2.4 | 1.3 | 0.2 | 0.0 |
| Favored [%] | 89.8 | 98.0 | 97.6 | 88.2 | 89.9 | 95.9 | 97.7 |

N/A = not applicable

**Appendix Table S1. Cryo-EM data and structure quality indicators – part 3.**

| | **Upper arm of baseplate of extended tail** | **Upper arm of baseplate of extended tail (segment A)** | **Upper arm of baseplate of extended tail (segment B)** | **Upper arm of baseplate of extended tail (segment CDEF)** |
|---|---|---|---|---|
| **Sample** | Particles with extended tails | Particles with extended tails | Particles with extended tails | Particles with extended tails |
| **Data collection and processing** | | | | |
| Detector | K3 | K3 | K3 | K3 |
| Magnification | 105,000 | 105,000 | 105,000 | 105,000 |
| Voltage [kV] | 300 | 300 | 300 | 300 |
| Exposure [$e^-$/Å$^2$] | 40.8 | 40.8 | 40.8 | 40.8 |
| Pixel size [Å] | 0.8336 | 0.8336 | 0.8336 | 0.8336 |
| Symmetry | C1 | C1 | C1 | C1 |
| Final number of particles | 19,727 | 19,727 | 19,727 | 14,954 |
| Map resolution [Å] | 5.1 | 3.8 | 3.9 | 5.2 |
| FSC threshold | 0.143 | 0.143 | 0.143 | 0.143 |
| **Accession codes** | | | | |
| EMDB | EMD-55961 | EMD-55962 | EMD-55963 | EMD-55964 |
| PDB | 9TIM | 9TIN | 9TIO | 9TIP |
| **Model composition** | | | | |
| Atoms (except hydrogens) | 97,116 | 7,390 | 15,828 | 15,113 |
| Residues | 12,310 | 936 | 1,981 | 1,915 |
| B-factor | 132.6 | 31.1 | 28.5 | 115.4 |
| FSC (model) at 0.5 (masked) | 5.2 | 4.0 | 4.0 | 5.6 |
| **RMSD** | | | | |
| Bond length [Å] | 0.013 | 0.002 | 0.002 | 0.013 |
| Bond angles [°] | 1.93 | 0.48 | 0.48 | 1.98 |
| **Validation** | | | | |
| MolProbity score [percentile] | 1.79 | 1.23 | 1.13 | 1.85 |
| ClashScore [percentile] | 1.16 | 4.58 | 3.36 | 1.43 |
| Poor rotamers [%] | 4.27 | 0.24 | 0.22 | 4.89 |
| C-β outliers [%] | 0.48 | N/A | N/A | 0.55 |
| **Ramachandran plot** | | | | |
| Outliers [%] | 1.0 | 0.0 | 0.0 | 1.1 |
| Favored [%] | 92.0 | 98.4 | 98.3 | 93.0 |

N/A = not applicable

**Appendix Table S1. Cryo-EM data and structure quality indicators – part 4.**

| | Baseplate of contracted tail (core, wedge modules, arms, proximal tail) | Baseplate of contracted tail (core, wedge modules, arms, proximal tail) | Baseplate of contracted tail (core, wedge modules, arms, proximal tail, tube) - composite | Baseplate of contracted tail (core, wedge modules, arm segment, inner tripods, proximal tail) | Baseplate of contracted tail (core, wedge modules, inner tripods) | Arm of baseplate of contracted tail | Arm of baseplate of contracted tail (segment B) |
|---|---|---|---|---|---|---|---|
| **Sample** | Particles with contracted tails | Particles with contracted tails | Particles with contracted tails | Particles with contracted tails | Particles with contracted tails | Particles with contracted tails | Particles with contracted tails |
| **Data collection and processing** | | | | | | | |
| Detector | K2 Summit | K2 Summit | K2 Summit | K2 Summit | K2 Summit | K2 Summit | K2 Summit |
| Magnification | 130,000 | 130,000 | 130,000 | 130,000 | 130,000 | 130,000 | 130,000 |
| Voltage [kV] | 300 | 300 | 300 | 300 | 300 | 300 | 300 |
| Exposure [$e^-$/Å$^2$] | 42 | 42 | 42 | 42 | 42 | 42 | 42 |
| Pixel size [Å] | 1.057 | 1.057 | 1.057 | 1.057 | 1.057 | 1.057 | 1.057 |
| Symmetry | C1 | C6 | C6 | C6 | C6 | C1 | C1 |
| Final number of particles | 25,483 | 25,483 | N/A | 25,203 | 25,203 | 15,390 | 56,778 |
| Map resolution [Å] | 6.6 | 4.4 | N/A | 4.0 | 3.1 | 4.1 | 4.4 |
| FSC threshold | 0.143 | 0.143 | N/A | 0.143 | 0.143 | 0.143 | 0.143 |
| **Accession codes** | | | | | | | |
| EMDB | EMD-55966 | EMD-55967 | EMD-55978 | EMD-19972 | EMD-19973 | EMD-55968 | EMD-55969 |
| PDB | N/A | 9TIR | 9TIW | 9EUJ | 9EUK | 9TIS | 9TIT |
| **Model composition** | | | | | | | |
| Atoms (except hydrogens) | N/A | 98,679 | 115,329 | 56,898 | 29,740 | 64,703 | 11,197 |
| Residues | N/A | 12,541 | 14,656 | 7,232 | 3,756 | 8,214 | 1,407 |
| B-factor | N/A | 300.0 | 183.9 | 71.5 | 49.6 | 27.7 | 69.8 |
| FSC (model) at 0.5 (masked) | N/A | 6.2 | 6.7 | 4.1 | 3.3 | 4.2 | 4.5 |
| **RMSD** | | | | | | | |
| Bond length [Å] | N/A | 0.011 | 0.010 | 0.006 | 0.005 | 0.013 | 0.002 |
| Bond angles [°] | N/A | 1.65 | 1.53 | 0.88 | 0.78 | 1.92 | 0.49 |
| **Validation** | | | | | | | |
| MolProbity score [percentile] | N/A | 1.47 | 1.42 | 1.16 | 0.94 | 1.64 | 1.21 |
| ClashScore [percentile] | N/A | 1.49 | 1.72 | 1.23 | 0.90 | 1.71 | 4.29 |
| Poor rotamers [%] | N/A | 1.81 | 1.57 | 0.43 | 0.12 | 2.44 | 0.24 |
| C-β outliers [%] | N/A | 0.23 | 0.21 | N/A | N/A | 0.33 | N/A |
| **Ramachandran plot** | | | | | | | |
| Outliers [%] | N/A | 0.5 | 0.4 | 0.1 | 0 | 0.7 | 0.0 |
| Favored [%] | N/A | 94.3 | 95.0 | 95.4 | 97.1 | 93.5 | 98.1 |

N/A = not applicable

**Appendix Table S1. Cryo-EM data and structure quality indicators – part 5.**

| | **Extended tail** | **Tail tube from phage 812 with contracted tail** | **Tail sheath from phage 812 with contracted tail** | **Central spike protein (knob and petal domains)** |
|---|---|---|---|---|
| **Sample** | Particles with extended tails | Particles with contracted tails | Particles with contracted tails | Recombinant central spike protein |
| **Data collection and processing** | | | | |
| Detector | Falcon 2 | K2 Summit | Falcon 2 | K2 Summit |
| Magnification | 75,000 | 130,000 | 75,000 | 130,000 |
| Voltage [kV] | 300 | 300 | 300 | 300 |
| Exposure [$e^-$/Å$^2$] | 48 | 42 | 48 | 42 |
| Pixel size [Å] | 1.061 | 1.057 | 1.061 | 1.04 |
| Symmetry | C6 | C6 | C6 | C3 |
| Final number of particles | 18,188 | 12,906 | 69,969 | 135,736 |
| Map resolution [Å] | 4.2 | 3.7 | 3.2 | 3.2 |
| FSC threshold | 0.143 | 0.143 | 0.143 | 0.143 |
| **Accession codes** | | | | |
| EMDB | EMD-50093 | EMD-50094 | EMD-50095 | EMD-19974 |
| PDB | 9F04 | 9F05 | 9F06 | 9EUL |
| **Model composition** | | | | |
| Atoms (except hydrogens) | 20,968 | 3,330 | 34,568 | 9,132 |
| Residues | 2,680 | 423 | 4,424 | 1,146 |
| B-factor | 89.6 | 55.0 | 98.5 | 69.6 |
| FSC (model) at 0.5 (masked) | 4.2 | 3.9 | 3.3 | 3.3 |
| **RMSD** | | | | |
| Bond length [Å] | 0.005 | 0.001 | 0.002 | 0.002 |
| Bond angles [°] | 0.73 | 0.40 | 0.44 | 0.44 |
| **Validation** | | | | |
| MolProbity score [percentile] | 0.94 | 0.96 | 0.99 | 1.08 |
| ClashScore [percentile] | 0.96 | 1.96 | 2.19 | 2.71 |
| Poor rotamers [%] | 0.18 | 0 | 0 | 0 |
| C-β outliers [%] | 0 | N/A | N/A | N/A |
| **Ramachandran plot** | | | | |
| Outliers [%] | 0 | 0 | 0 | 0 |
| Favored [%] | 97.1 | 99.0 | 98.4 | 97.9 |

N/A = not applicable

**Appendix Table S2. List of proteins identified in phage 812 particles by mass spectrometry.**

| ORF | Protein | MW [kDa] | NSAF [%] | Coverage [%] | Number of unique peptides | Present in phages with contracted tail |
|---|---|---|---|---|---|---|
| 109 | Tape measure | 144 | 30 | 66 | 85 | no |
| 119 | Tripod | 129 | 82 | 87 | 99 | yes |
| 117 | Arm scaffold | 116 | 40 | 81 | 73 | yes |
| 112 | Central spike | 96 | 24 | 80 | 56 | yes |
| 103 | Tail sheath | 65 | 77 | 88 | 58 | yes |
| 91 | Portal | 64 | 46 | 75 | 50 | yes |
| 94 | Major capsid | 51 | 93 | 90 | 40 | yes |
| 123 | Receptor-Binding 2 | 50 | 36 | 93 | 36 | yes |
| 116 | Wedge | 39 | 43 | 67 | 25 | yes |
| 97 | Stopper | 34 | 57 | 77 | 28 | yes |
| 96 | Adaptor | 34 | 6 | 95 | 19 | yes |
| 99 | Terminator | 32 | 42 | 60 | 15 | yes |
| 113 | Tail tube initiator | 29 | 55 | 87 | 24 | yes |
| 115 | Tail sheath initiator | 27 | 54 | 80 | 13 | yes |
| 154 | Minor Capsid | 23 | 66 | 88 | 15 | no |
| 89 | Terminase | 30 | 47 | 64 | 20 | yes |
| 156 | Minor capsid | 18 | 65 | 96 | 17 | no |
| 114 | Assembly | 18 | 31 | 94 | 12 | yes |
| 118 | Arm segment | 19 | 63 | 92 | 19 | yes |
| 102 | Tail tube | 13 | 95 | 72 | 12 | yes |
| 157 | Minor capsid | 8 | 49 | 81 | 5 | no |
| 56 | Terminator cement | 10 | 35 | 77 | 9 | yes |
| 77 | Vertex cement | 8 | 81 | 94 | 11 | yes |

NSAF – normalized spectral abundance factor.

**Appendix Table S3. List of structural proteins from phage 812 tail and baseplate.**

| ORF | protein | residues | domains (residue range) | representative homologs |
|---|---|---|---|---|
| 101 | Tail sheath | 587 | I (491-587) | Phage T4, gp18 (Taylor *et al*, 2016); phage E217, gp31 (Li *et al*, 2023); *V. cholerae* Type VI Secretion System, VipA/B (Kudryashev *et al*, 2015) |
| | | | II (34-96,313-490) | |
| | | | III (97-151,251-312) | Prophage *L. innocua*, Lin1278, PDB: 3LML |
| | | | IV (152-250) | |
| 102 | Tail tube | 142 | β-barrel (1-34,59-142) | Phage T4, gp19 (Taylor *et al*, 2016); *P. aeruginosa* R2 pyocin, PA0626 (Ge *et al*, 2020), *C. difficile* diffocine, CD1364 (Cai *et al*, 2024) |
| | | | stacking loop (35-58) | |
| 110 | Hub | 810 | tube-forming I (1-125) | Phage T4, gp27 (Kanamaru *et al*, 2002); *E. coli* Type VI Secretion System, VgrG (Leiman *et al*, 2009); *P. aeruginosa* R2 pyocin, PA0628 (Ge *et al*, 2020); phage 80alpha, ORF59 (Kizziah *et al*, 2020) |
| | | | tube-forming II (126-266) | |
| | | | tube-forming III (267-285,546-641) | |
| | | | tube-forming IV (286-334,527-545) | |
| | | | clamp (358-511) | *V. vulnificus*, PAS factor (Lee *et al*, 2006) |
| | | | cleaver (642-808) | Phage K, ORF56 (tail-associated lysin) (Paul *et al*, 2011); phage K, LysK (endolysin) (Becker *et al*, 2008) |
| 111 | Weld | 295 | lid (1-128) | *R. typhi* Type IV Secretion System, VirB8 (Gillespie *et al*, 2015); *H. pylori* Type IV Secretion System, CagV (Wu *et al*, 2019) |
| | | | linker (129-142) | |
| | | | clip (143-295) | *B. cereus*, YkfC (Xu *et al*, 2010); *M. tuberculosis*, RipD (Böth *et al*, 2014) |
| 112 | Central spike | 848 | oligosaccharide-binding (OB) (1-110) | Phage T4, gp5 (Kanamaru *et al*, 2002); phage phi92, gp138 (spike )(Browning *et al*, 2012) |
| | | | β-prism (111-310) | |
| | | | coiled-coil (311-466) | Phage P22, gp26 (tail needle) (Bhardwaj *et al*, 2009) |
| | | | knob (467-610) | Phage Sf6, tail needle (Bhardwaj *et al*, 2011); phage 1358, receptor-binding protein (Farenc *et al*, 2014); *B. cenocepacia,* BC2L-C (lectin) (Šulák *et al*, 2010) |
| | | | petal (611-848) | *B. subtilis*, GlpQ (phosphodiesterase) (Shi *et al*, 2008) |
| 113 | Tail tube initiator | 263 | tube-like (1-156) | Phage T4, gp54/gp48 (Taylor *et al*, 2016); *P. aeruginosa* R2 pyocin, PA0626 (ripcord) (Ge *et al*, 2020); *C. difficile* diffocine, CD1367 (Cai *et al*, 2024) |
| | | | handle (157-263) | *P. aeruginosa* R2 pyocin, PA0626 (ripcord )(Ge *et al*, 2020); phage E217, gp37/gp38 (Li *et al*, 2023); phage Pam3, gp17 (Yang *et al*, 2023) |

| | | | | |
|---|---|---|---|---|
| 114 | Assembly | 173 | - | *P. aeruginosa* R2 pyocin, PA0626 (ripcord) (Ge *et al*, 2020); phage Milano, BCP (gp25) (Sonani *et al*, 2024) |
| 115 | Tail sheath initiator | 234 | LysM (1-66) | Phage T4, gp53 (Taylor *et al*, 2016); *P. aeruginosa* R2 pyocin, PA0627 (glue) (Ge *et al*, 2020); phage Milano, BCP (gp25) (Sonani *et al*, 2024) |
| | | | linker (67-92) | - |
| | | | handshake (93-234) | Phage T4, gp25 (Taylor *et al*, 2016); *P. aeruginosa* R2 pyocin, PA0617 (sheath initiator) (Ge *et al*, 2020); phage E217, gp34 (Li *et al*, 2023); phage Pam3, gp17 (Yang *et al*, 2023) |
| 116 | Wedge A, B | 348 | helices (1-67) | Phage T4, gp6 (Taylor *et al*, 2016); *E. coli* Type VI Secretion System, TssFa/b (Park *et al*, 2018); *C. difficile* diffocine, CD1364 (Cai *et al*, 2024) |
| | | | wing (68-192) | *C. difficile* diffocine, CD1364 (Cai *et al*, 2024); phage E217, gp44 (Li *et al*, 2023); phage Milano, BCP (gp25) (Sonani *et al*, 2024) ; phage A1(L), gp31 (Yu *et al*, 2024) |
| | | | trifurcation unit (193-256) | Phage T4, gp6 (Taylor *et al*, 2016); *E. coli* Type VI Secretion System, TssFa/b (Park *et al*, 2018); *C. difficile* diffocine, CD1364 (Cai *et al*, 2024) |
| | | | dimerization (257-348) | |
| 117 | Arm scaffold | 1019 | helices (1-94) | Phage T4, gp7 (Taylor *et al*, 2016); *E. coli* Type VI Secretion System, TssG (Park *et al*, 2018); *P. aeruginosa* R2 pyocin, PA0619 (Tri2) (Ge *et al*, 2020); phage E217, gp45 (Li *et al*, 2023); phage Pam3, gp23 (Yang *et al*, 2023) |
| | | | trifurcation (95-116,145-176) | |
| | | | docking loop 1 (117-144) | |
| | | | linker 1 (177-212) | - |
| | | | bulge (213-219,259-443) | *A. thermocellus*, Xyn10B (carbohydrate-binding module 22) (Najmudin *et al*, 2010); *P. barcinonensis*, Xyn10C (carbohydrate-binding module 22) (Sainz-Polo *et al*, 2015) |
| | | | docking loop 2 (220-258) | |
| | | | linker 2 (444-489) | - |
| | | | elbow (490-776) | Phage Tuc2009, BppA (putative receptor-binding protein) (Legrand *et al*, 2016); *P. barcinonensis*, Xyn10C (family 22 carbohydrate-binding module) (Sainz-Polo *et al*, 2015) |
| | | | linker 3 (777-907) | - |
| | | | C-terminal (908-1019) | *B. ovatus*, putative alpha-L-fucosidase, (4ZRX); *C. polysaccharolyticus*, family 16 carbohydrate-binding module (Bae *et al*, 2008) |
| 118 | Arm segment | 173 | - | Phage T4, gp8 (Taylor *et al*, 2016); A511, gp105 (Guerrero-Ferreira *et al*, 2019) |
| 119 | Tripod | 1152 | spine helices (1-54,172-189,468-495) | - |
| | | | flap (55-115,139-171) | Phage 1358, receptor-binding protein (shoulder) (Farenc *et al*, 2014); *E. coli* Type VI Secretion System, TssK (Park *et al*, 2018) |
| | | | flap insertion loop (116-138) | |
| | | | fiber I (190-242,404-467) | |

| | | | | |
|---|---|---|---|---|
| | | | fiber II (243-276,359-403) | *R. capsulatus* Gene Transfer Agent, adaptor (Bárdy *et al*, 2020); phage YSD1, major tail protein (domain 2) (Hardy *et al*, 2020); phage CBA120, central spike (domain XD2/XD3) (Chao *et al*, 2022) |
| | | | fiber III (277-358) | |
| | | | base (496-538,1045-1152) | *C. polysaccharolyticus*, family 16 carbohydrate-binding module (Bae *et al*, 2008) ; *T. maritima*, family 61 carbohydrate-binding module (Cid *et al*, 2010) |
| | | | fin (539-565,944-1044) | |
| | | | leg (566-590,793-943) | *C. difficile*, Cwp84 (lectin-like) (Bradshaw *et al*, 2014) |
| | | | anchor (591-612,674-792) | SSV19, B120 (adaptor) (Han *et al*, 2022); *B. ovatus*, BoSGBP$_{MLG}$-B (Surface glycan-binding protein) (Tamura *et al*, 2019) |
| | | | hook (613-673) | - |
| 121 | Receptor-binding protein 1 | 640 | antenna (1-37) | - |
| | | | tower I (38-121) | Phage T4, gp34 (long tail fiber) (Granell *et al*, 2017); *P. aeruginosa* R1 pyocin, PA0620 (fiber) (Salazar *et al*, 2019) |
| | | | tower II (122-184,397-410) | |
| | | | wing (185-396) | Phage CBA120, tail spike protein 4 (Plattner *et al*, 2019); *T. polysaccharolyticum*, Family 16 Carbohydrate-binding Module (Bae *et al*, 2008) |
| | | | tower III (411-514) | Phage T4, gp34 (long tail fiber) (Granell *et al*, 2017); *P. aeruginosa* R1 pyocin, PA0620 (fiber) (Salazar *et al*, 2019) |
| | | | tip (515-640) | Phage T7, gp17 (fiber) (Garcia-Doval & van Raaij, 2012) |
| 122 | Coupler | 124 | - | Phage Mu, gpU (tail fiber assembly protein) (North *et al*, 2019) |
| 123 | Receptor-binding protein 2 | 458 | antenna (1-60) | - |
| | | | tower I (61-149) | Phage T4, gp34 (long tail fiber) (Granell *et al*, 2017); *P. aeruginosa* R1 pyocin, PA0620 (fiber) (Salazar *et al*, 2019) |
| | | | tower II (150-228) | |
| | | | tower III (229-327) | |
| | | | tip (328-458) | T7, gp17 (fiber) (Garcia-Doval & van Raaij, 2012) |

**Appendix Table S4. NMR data and structure quality indicators for cleaver domain of hub protein.**

| Structural statistics of 20 models | Cleaver domain of the hub protein |
|---|---|
| **NMR distance and dihedral restraints** | |
| Distance restraints | |
| Total NOE | 1632 |
| Intra-residue ($i$=$j$) | 460 |
| Sequential ($\|i – j\|$ = 1) | 439 |
| Medium-range (1 < $\|i – j\|$ < 5) | 223 |
| Long-range ($\|i – j\|$ > = 5) | 510 |
| Total dihedral angle restraints | 195 |
| $\varphi$ | 95 |
| $\psi$ | 100 |
| Violations (mean ± s.d.) | |
| Distance restraints (Å) | 0.020 ± 0.002 |
| Dihedral angle restraints (º) | 0.81 ± 0.09 |
| Max. dihedral angle violation (º) | 5.7 |
| Max. distance constraint violation (Å) | 0.95 |
| Deviations from idealized geometry | |
| Bond lengths (Å) | 0.011± 0.000 |
| Bond angles (º) | 1.12± 0.032 |
| Impropers (º) | 1.88 ± 0.071 |
| Average pairwise r.m.s. deviation* (Å) | |
| Backbone | 0.99 ± 0.26 |
| Heavy | 1.36 ± 0.23 |
| Ramachandran plot statistics* (%) | |
| Residues in most favoured regions | 96.9 |
| Residues in additionally allowed regions | 2.7 |
| Residues in generously allowed regions | 0.1 |
| Residues in disallowed regions | 0.3 |

* Ordered residues: 25-42, 51-100, 103-129, 135-159

**Appendix Table S5. X-ray data collection and refinement statistics.**

| **Data collection** | Arm segment protein |
| --- | --- |
| Space group | P2(1) |
| Cell axes a, b, c [Å] | 92.93, 75.14, 94.42 |
| Cell angles α, β, γ [°] | 90.00, 109.87, 90.00 |
| Beamline | Soleil, Proxima 2A |
| Resolution range [Å]* | 46.49-1.80 (1.86-1.80) |
| Reflections* | 109 411 (8 517) |
| Completeness [%]* | 96 (76) |
| Mean <I/s(I)>* | 3.04 (1.20) |
| $R_{merge}$ [%]* | 0.146 (0.520) |
| Wilson B [Å$^2$] | 38.12 |
| **Refinement** | |
| Reflections $R_{free}$* | 5471 (426) |
| *R*-factor* | 0.245 (0.374) |
| *R*-free* | 0.275 (0.399) |
| Overall B-factor [Å$^2$] | 43.5 |
| Ramachandran stats [%] | 98.2 |
| rmsd bonds (Å)/angles [°] | 0.011; 1.15 |
| PDB code | 9FKO |

* Numbers in brackets are for the high-resolution bin.

### References for Appendix Table S3

Bae B, Ohene-Adjei S, Kocherginskaya S, Mackie RI, Spies MA, Cann IKO & Nair SK (2008) Molecular Basis for the Selectivity and Specificity of Ligand Recognition by the Family 16 Carbohydrate-binding Modules from Thermoanaerobacterium polysaccharolyticum ManA. *J Biol Chem* 283: 12415–12425

Bárdy P, Füzik T, Hrebík D, Pantůček R, Thomas Beatty J & Plevka P (2020) Structure and mechanism of DNA delivery of a gene transfer agent. *Nat Commun* 11: 3034

Becker SC, Foster-Frey J & Donovan DM (2008) The phage K lytic enzyme LysK and lysostaphin act synergistically to kill MRSA. *FEMS Microbiol Lett* 287: 185–191

Bhardwaj A, Molineux IJ, Casjens SR & Cingolani G (2011) Atomic Structure of Bacteriophage Sf6 Tail Needle Knob. *J Biol Chem* 286: 30867–30877

Bhardwaj A, Walker-Kopp N, Casjens SR & Cingolani G (2009) An Evolutionarily Conserved Family of Virion Tail Needles Related to Bacteriophage P22 gp26: Correlation between Structural Stability and Length of the α-Helical Trimeric Coiled Coil. *J Mol Biol* 391: 227–245

Böth D, Steiner EM, Izumi A, Schneider G & Schnell R (2014) RipD (Rv1566c) from *Mycobacterium tuberculosis* : adaptation of an NlpC/p60 domain to a non-catalytic peptidoglycan-binding function. *Biochem J* 457: 33–41

Bradshaw WJ, Kirby JM, Thiyagarajan N, Chambers CJ, Davies AH, Roberts AK, Shone CC & Acharya KR (2014) The structure of the cysteine protease and lectin-like domains of Cwp84, a surface layer-associated protein from *Clostridium difficile*. *Acta Crystallogr D Biol Crystallogr* 70: 1983–1993

Browning C, Shneider MM, Bowman VD, Schwarzer D & Leiman PG (2012) Phage Pierces the Host Cell Membrane with the Iron-Loaded Spike. *Structure* 20: 326–339

Cai X, He Y, Yu I, Imani A, Scholl D, Miller JF & Zhou ZH (2024) Atomic structures of a bacteriocin targeting Gram-positive bacteria. *Nat Commun* 15: 7057

Chao KL, Shang X, Greenfield J, Linden SB, Alreja AB, Nelson DC & Herzberg O (2022) Structure of Escherichia coli O157:H7 bacteriophage CBA120 tailspike protein 4 baseplate anchor and tailspike assembly domains (TSP4-N). *Sci Rep* 12: 2061

Cid M, Pedersen HL, Kaneko S, Coutinho PM, Henrissat B, Willats WGT & Boraston AB (2010) Recognition of the Helical Structure of β-1,4-Galactan by a New Family of Carbohydrate-binding Modules. *J Biol Chem* 285: 35999–36009

Farenc C, Spinelli S, Vinogradov E, Tremblay D, Blangy S, Sadovskaya I, Moineau S & Cambillau C (2014) Molecular Insights on the Recognition of a Lactococcus lactis Cell Wall Pellicle by the Phage 1358 Receptor Binding Protein. *J Virol* 88: 7005–7015

Garcia-Doval C & van Raaij MJ (2012) Structure of the receptor-binding carboxy-terminal domain of bacteriophage T7 tail fibers. *Proc Natl Acad Sci* 109: 9390–9395

Ge P, Scholl D, Prokhorov NS, Avaylon J, Shneider MM, Browning C, Buth SA, Plattner M, Chakraborty U, Ding K, *et al* (2020) Action of a minimal contractile bactericidal nanomachine. *Nature* 580: 658–662

Gillespie JJ, Phan IQH, Scheib H, Subramanian S, Edwards TE, Lehman SS, Piitulainen H, Sayeedur Rahman M, Rennoll-Bankert KE, Staker BL, *et al* (2015) Structural Insight into How Bacteria Prevent Interference between Multiple Divergent Type IV Secretion Systems. *mBio* 6: e01867-15

Granell M, Namura M, Alvira S, Kanamaru S & van Raaij M (2017) Crystal Structure of the Carboxy-Terminal Region of the Bacteriophage T4 Proximal Long Tail Fiber Protein Gp34. *Viruses* 9: 168

Guerrero-Ferreira RC, Hupfeld M, Nazarov S, Taylor NM, Shneider MM, Obbineni JM, Loessner MJ, Ishikawa T, Klumpp J & Leiman PG (2019) Structure and transformation of bacteriophage A511 baseplate and tail upon infection of *Listeria* cells. *EMBO J* 38

Han Z, Yuan W, Xiao H, Wang L, Zhang J, Peng Y, Cheng L, Liu H & Huang L (2022) Structural insights into a spindle-shaped archaeal virus with a sevenfold symmetrical tail. *Proc Natl Acad Sci* 119: e2119439119

Hardy JM, Dunstan RA, Grinter R, Belousoff MJ, Wang J, Pickard D, Venugopal H, Dougan G, Lithgow T & Coulibaly F (2020) The architecture and stabilisation of flagellotropic tailed bacteriophages. *Nat Commun* 11: 3748

Kanamaru S, Leiman PG, Kostyuchenko VA, Chipman PR, Mesyanzhinov VV, Arisaka F & Rossmann MG (2002) Structure of the cell-puncturing device of bacteriophage T4. *Nature* 415: 553–557

Kizziah JL, Manning KA, Dearborn AD & Dokland T (2020) Structure of the host cell recognition and penetration machinery of a Staphylococcus aureus bacteriophage. *PLOS Pathog* 16: e1008314

Kudryashev M, Wang RY-R, Brackmann M, Scherer S, Maier T, Baker D, DiMaio F, Stahlberg H, Egelman EH & Basler M (2015) Structure of the Type VI Secretion System Contractile Sheath. *Cell* 160: 952–962

Lee JH, Yang S-T, Rho S-H, Im YJ, Kim SY, Kim YR, Kim M-K, Kang GB, Kim JI, Rhee JH, *et al* (2006) Crystal Structure and Functional Studies Reveal that PAS Factor from Vibrio vulnificus is a Novel Member of the Saposin-fold Family. *J Mol Biol* 355: 491–500

Legrand P, Collins B, Blangy S, Murphy J, Spinelli S, Gutierrez C, Richet N, Kellenberger C, Desmyter A, Mahony J, *et al* (2016) The Atomic Structure of the Phage Tuc2009 Baseplate Tripod Suggests that Host Recognition Involves Two Different Carbohydrate Binding Modules. *mBio* 7

Leiman PG, Basler M, Ramagopal UA, Bonanno JB, Sauder JM, Pukatzki S, Burley SK, Almo SC & Mekalanos JJ (2009) Type VI secretion apparatus and phage tail-associated protein complexes share a common evolutionary origin. *Proc Natl Acad Sci* 106: 4154–4159

Li F, Hou C-FD, Lokareddy RK, Yang R, Forti F, Briani F & Cingolani G (2023) High-resolution cryo-EM structure of the Pseudomonas bacteriophage E217. *Nat Commun* 14: 4052

Najmudin S, Pinheiro BA, Prates JAM, Gilbert HJ, Romão MJ & Fontes CMGA (2010) Putting an N-terminal end to the Clostridium thermocellum xylanase Xyn10B story: Crystal structure of the CBM22-1–GH10 modules complexed with xylohexaose. *J Struct Biol* 172: 353–362

North OI, Sakai K, Yamashita E, Nakagawa A, Iwazaki T, Büttner CR, Takeda S & Davidson AR (2019) Phage tail fibre assembly proteins employ a modular structure to drive the correct folding of diverse fibres. *Nat Microbiol* 4: 1645–1653

Park Y-J, Lacourse KD, Cambillau C, DiMaio F, Mougous JD & Veesler D (2018) Structure of the type VI secretion system TssK–TssF–TssG baseplate subcomplex revealed by cryo-electron microscopy. *Nat Commun* 9: 5385

Paul VD, Rajagopalan SS, Sundarrajan S, George SE, Asrani JY, Pillai R, Chikkamadaiah R, Durgaiah M, Sriram B & Padmanabhan S (2011) A novel bacteriophage Tail-Associated Muralytic Enzyme (TAME) from Phage K and its development into a potent antistaphylococcal protein. *BMC Microbiol* 11: 226

Plattner M, Shneider MM, Arbatsky NP, Shashkov AS, Chizhov AO, Nazarov S, Prokhorov NS, Taylor NMI, Buth SA, Gambino M, *et al* (2019) Structure and Function of the Branched Receptor-Binding Complex of Bacteriophage CBA120. *J Mol Biol* 431: 3718–3739

Sainz-Polo MA, González B, Menéndez M, Pastor FIJ & Sanz-Aparicio J (2015) Exploring Multimodularity in Plant Cell Wall Deconstruction. *J Biol Chem* 290: 17116–17130

Salazar AJ, Sherekar M, Tsai J & Sacchettini JC (2019) R pyocin tail fiber structure reveals a receptor-binding domain with a lectin fold. *PLOS ONE* 14: e0211432

Shi L, Liu J-F, An X-M & Liang D-C (2008) Crystal structure of glycerophosphodiester phosphodiesterase (GDPD) from *Thermoanaerobacter tengcongensis* , a metal ion-dependent enzyme: Insight into the catalytic mechanism: Crystal Structure of GDPD. *Proteins Struct Funct Bioinforma* 72: 280–288

Sonani RR, Palmer LK, Esteves NC, Horton AA, Sebastian AL, Kelly RJ, Wang F, Kreutzberger MAB, Russell WK, Leiman PG, *et al* (2024) An extensive disulfide bond network prevents tail contraction in Agrobacterium tumefaciens phage Milano. *Nat Commun* 15: 756

Šulák O, Cioci G, Delia M, Lahmann M, Varrot A, Imberty A & Wimmerová M (2010) A TNF-like Trimeric Lectin Domain from *Burkholderia cenocepacia* with Specificity for Fucosylated Human Histo-Blood Group Antigens. *Structure* 18: 59–72

Tamura K, Foley MH, Gardill BR, Dejean G, Schnizlein M, Bahr CME, Louise Creagh A, van Petegem F, Koropatkin NM & Brumer H (2019) Surface glycan-binding proteins are essential for cereal beta-glucan utilization by the human gut symbiont Bacteroides ovatus. *Cell Mol Life Sci* 76: 4319–4340

Taylor NMI, Prokhorov NS, Guerrero-Ferreira RC, Shneider MM, Browning C, Goldie KN, Stahlberg H & Leiman PG (2016) Structure of the T4 baseplate and its function in triggering sheath contraction. *Nature* 533: 346–352

Wu X, Zhao Y, Sun L, Jiang M, Wang Q, Wang Q, Yang W & Wu Y (2019) Crystal structure of CagV, the *Helicobacter pylori* homologue of the T4 SS protein VirB8. *FEBS J* 286: 4294–4309

Xu Q, Abdubek P, Astakhova T, Axelrod HL, Bakolitsa C, Cai X, Carlton D, Chen C, Chiu H-J, Chiu M, *et al* (2010) Structure of the γ- D -glutamyl- L -diamino acid endopeptidase YkfC from *Bacillus cereus* in complex with L -Ala-γ- D -Glu: insights into substrate recognition by NlpC/P60 cysteine peptidases. *Acta Crystallograph Sect F Struct Biol Cryst Commun* 66: 1354–1364

Yang F, Jiang Y-L, Zhang J-T, Zhu J, Du K, Yu R-C, Wei Z-L, Kong W-W, Cui N, Li W-F, *et al* (2023) Fine structure and assembly pattern of a minimal myophage Pam3. *Proc Natl Acad Sci* 120: e2213727120

Yu R-C, Yang F, Zhang H-Y, Hou P, Du K, Zhu J, Cui N, Xu X, Chen Y, Li Q, *et al* (2024) Structure of the intact tail machine of Anabaena myophage A-1(L). *Nat Commun* 15: 2654
